# High Throughput Characterization of Eukaryotic 2A-Like Peptides Identifies Novel Leucine-Associated Reduction in Protein Abundance

**DOI:** 10.64898/2026.06.17.732966

**Authors:** John C. Snell, Kenneth A. Matreyek

## Abstract

Virally-derived ribosomal skipping 2A peptides are a popular tool for protein co-expression. Despite their use in over 9,000 publications, the biochemical and biophysical properties underlying the skipping mechanism remain largely unexplored. We identified 4,218 2A-like peptides originating from non-viral organisms. We developed and utilized the Trifluorescent Reporter fluorescent tool for high-throughput multiplexable analysis of ribosomal skipping, and tested 3,271 2A-like peptide sequences. We identified peptides that skipped, failed to skip, and skipped but failed to restart translation, in addition to peptides that induced a reduction in protein abundance. Peptides that skipped and induced reductions in protein abundance largely originated from eukaryotes. A poly-leucine stretch in an alpha-helix N-terminal to the conserved GDxExNPGP motif drove both skipping and the reduction in protein abundance. Analysis of the native eukaryotic protein contexts revealed that reduction may be harnessed as an expression regulator. The high-throughput approach used in this work greatly expands the functional knowledge of what biophysical and biochemical characteristics lead to ribosomal skipping, including an apparent latent eukaryotic ‘leucine stall-helix’ motif.

**Graphical Abstract:** 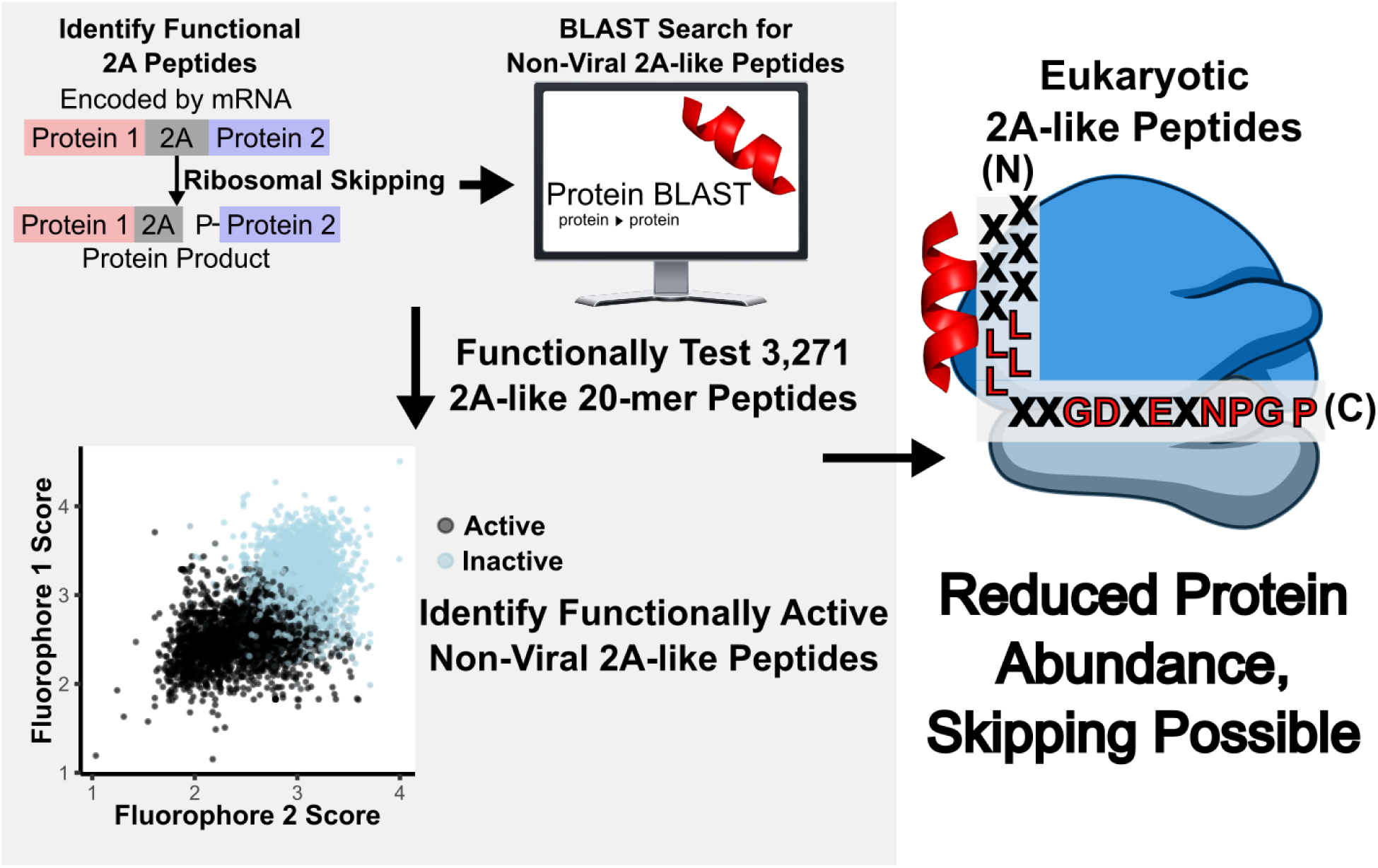

## Introduction

Ribosomal skipping sequences were first discovered in positive-stranded RNA viruses where they aid in separating viral polyproteins. First described in Foot-and-Mouth Disease virus*^1,2^*, translation events of the 2A protein (F2A) fail to form the peptide bond between its C-terminal glycine and proline residues, producing two distinct protein products without proteolytic cleavage.

Outside of their viral context, 2A proteins (henceforth 2A peptides) maintain the capacity to skip, but do so incompletely*^3^*. 2A peptides have since become an essential component of recombinant DNA constructs due to the ability to co-translationally produce separate proteins. Aside from F2A, other commonly used 2A peptides include 2A from equine rhinitis A virus (E2A), porcine teschovirus-1 (P2A), and *Thosea asigna* virus (T2A)*^3^*. Functional 2A peptides have been found in other related viruses*^2,4–6^*, though these are not typically used in engineering contexts. Functional 2A peptides typically contain either a ‘canonical’ core C-terminal _-8_GDxExNPGP_+1_ motif (indexing relative to the skipped peptide bond) or a ‘non-canonical’ core C-terminal _-7_GxExNPGP_+1_ motif*^1,7,8^* and are defined as being 18-23 residues in length*^9,10^*.

Failure to skip yields one of two possible outcomes. The first, ribosomal ‘readthrough’, is where the peptide bond forms normally and a fusion protein is produced. The second, ribosomal ‘falloff’, is a hypothesized event where only the protein N-terminal to the 2A is produced*^2,11^*. The relative ratio of the expected outcomes comprises the ribosomal skipping efficiency of a given peptide, and is considered to be related to the specific order of upstream residues*^3,12^*.

The overall prevalence of ribosomal skipping sequences across life remains undetermined. There are dozens of experimentally characterized ribosomal skipping sequences of viral origin*^1,6,12^*. A handful of eukaryotic-origin ribosomal skipping sequences have been experimentally tested and shown to skip*^1,4,13–15^*, and ribosomal skipping has occurred in all tested eukaryotic systems*^1,3,16,17^*. Some putative skipping sequences of bacterial origin skip in eukaryotic cells*^15^*, but ribosomal skipping does not occur in bacterial cells*^1,15^*.

Experimental validation is the primary bottleneck to identifying ribosomal skipping sequences. The RefSeq database currently contains more than 470 million complete accessions, representing a more than 10-fold increase over the past decade*^18^*. Only high throughput experimental approaches are capable of testing the thousands of putative skipping sequences that are bioinformatically identifiable^15^.

We paired bioinformatic analysis with a novel high-throughput fluorescent assay to test for skipping in a library of 3,271 putative ribosomal skipping peptide sequences in cultured mammalian cells. We further tested a library of 218 synonymous variants of P2A, T2A, E2A, and F2A to resolve conflicting reports regarding the impact of codon sequence on skipping. We identified samples that perform each of the hypothesized skipping outcomes, as well as a number of peptides that induced reductions in protein abundance. We performed residue-position analysis and identified residue usage trends that correlate with skipping. We finally performed structural analysis to look at the native roles of 2A-like peptide sequences.

## Results

### Identification of Non-Viral 2A-Like Sequences

It has been unclear whether ribosomal skipping is dependent on the properties of the 2A-like residue sequence, the mRNA molecule being translated, or both due to conflicting reports in the literature*^19,20^*. Accordingly, we identified and tested candidate ribosomal skipping sequences using the original coding sequences. We identified 45 peptides that were previously reported to exhibit ribosomal skipping, and noted the 20 residues spanning from position −19 to position +1 relative to the skipped peptide bond for each. There were 36 viral and 9 non-viral peptides (Table S1). These peptides had a high bias for the canonical core motif, as well as a moderate bias of leucine from position −13 to −10 (Figure 1A).

**Figure 1.**
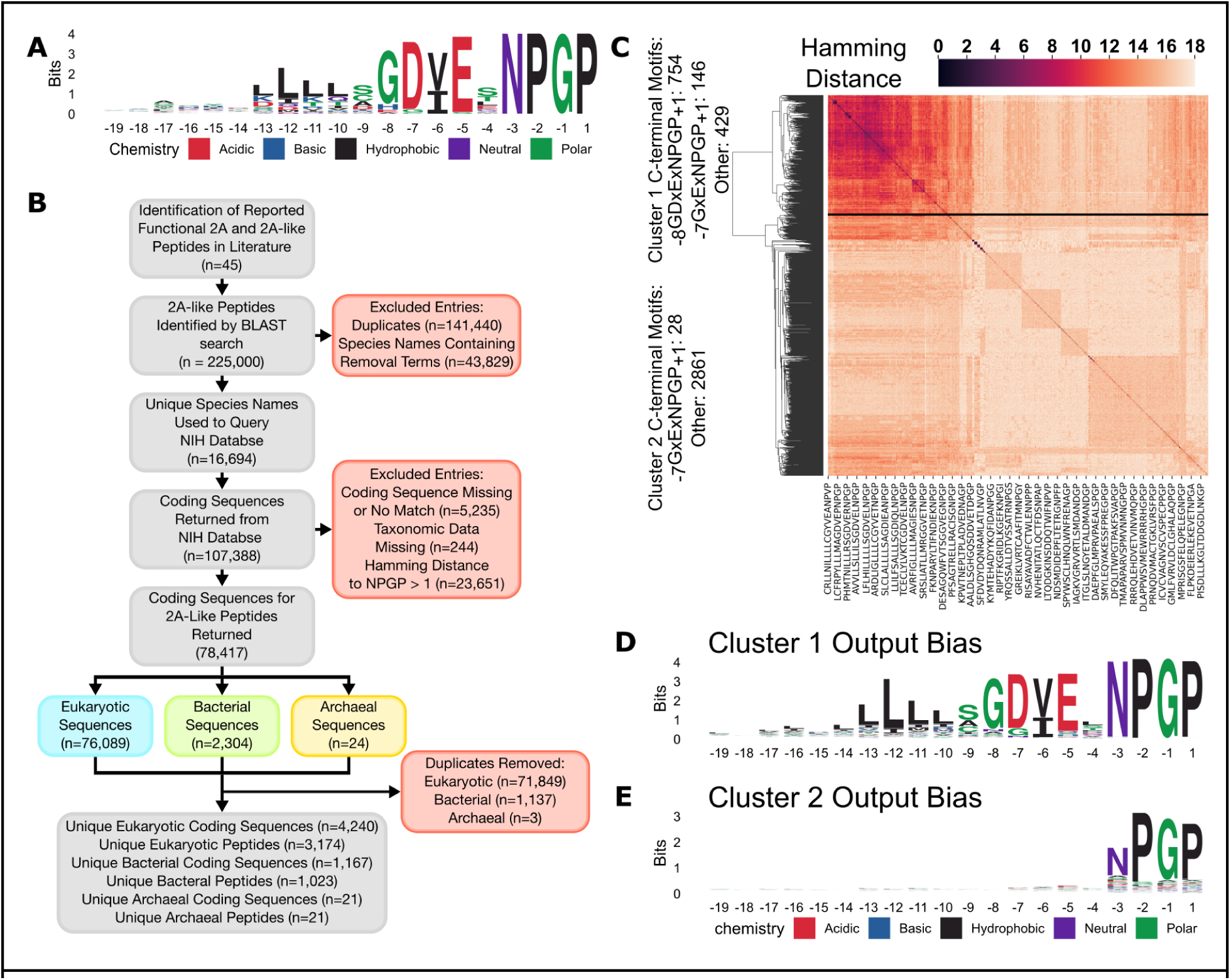
Identification of Non-Viral 2A like Peptides for Library Construction. (A) Positional residue biases of 45 previously identified functional skipping peptides (Table S1) used for seeding BLAST searches. (B) Search workflow for identification of candidate 2A-like peptides. BLAST was used to identify proteins that contain 2A-like peptides. Genomes of species the proteins originated from were queried for further 2A-like peptides, and original coding sequences were identified for use as library members. (C) Identity matrix of 20-mer 2A-like peptide sequences that were bioinformatically identified. Samples broadly populate 2 clusters. Cluster 1 contained samples with the canonical C-terminal motif _-8_GDxExNPGP_+1_, the non-canonical C-terminal motif _-7_GxExNPGP_+1_, and samples that are slightly deviant from these motifs. Cluster 2 contained highly deviant peptides. (D, E) Positional residue biases in output samples in (D) Cluster 1 and (E) Cluster 2.

Each peptide was used as the input for a Basic Local Alignment Search Tool (BLAST)*^21^* protein query. The top 5,000 matches by BLAST were recovered for each query, yielding 44,484 unique entries. Coding DNA for 8,487 species harboring 2A-like peptides was retrieved from NCBI datasets*^22^*. We found and recorded all native coding sequences for 20-mer peptides terminating proximal to ‘NPGP’ (Table S2) and identified 5,428 unique 2A-like coding sequences for 4,218 peptide sequences of eukaryotic, bacterial, and archaeal origin (Figure 1B) (Table S3).

The output peptide sequences exhibited high sequence diversity (Figure 1C). Hierarchical clustering identified that in the 2 major clusters determined by peptide sequence identity, one cluster contained peptides with both the canonical and non-canonical core motifs, and the other did not. Peptides in the latter cluster frequently had 6 or fewer residues in common with other peptides. Peptides in the cluster containing the canonical and non-canonical core motifs showed a similar residue-position bias as the input for the bioinformatic search (Figure 1D), while peptides in the other cluster showed only bias toward _-3_NPGP_+1_ (Figure 1E), largely imposed by the filtering step of our bioinformatic pipeline.Development of the Trifluorescent Reporter for Skipping

### Development of the Trifluorescent Reporter for Skipping

Testing the thousands of identified 2A-like peptides required a scalable, multiplexable assay for identifying ribosomal skipping. We developed the Trifluorescent protein Reporter (TR) to facilitate such high-throughput analysis of ribosomal skipping. The construct consists of a central green fluorescent protein, monomeric Kusabira Green (mKG), with the red fluorescent protein mCherry fused N-terminally and the near infrared fluorescent protein miRFP670 fused C-terminally. There is also an N-terminal HA tag and a C-terminal DYKDDDDK (FLAG) tag to allow for western blotting of either ends of the construct (Figure 2A, Figure S1). We engineered two BsaI Type IIS restriction enzyme sites flanking a mutable flexible loop in mKG*^23^*, enabling insertion of 2A-like peptides using golden-gate cloning*^24^*. The TR is encoded in a promoterless plasmid immediately following an attB Bxb1 serine recombination site. The TR plasmid is compatible with cells containing an attP Bxb1 recombination site, permitting stable expression of a single site-specific genomically-integrated plasmid copy per cell*^25,26^*. A strict genotype-phenotype link is established for each cell expressing the TR, allowing for high-throughput multiplexable measurement of skipping.

**Figure 2.**
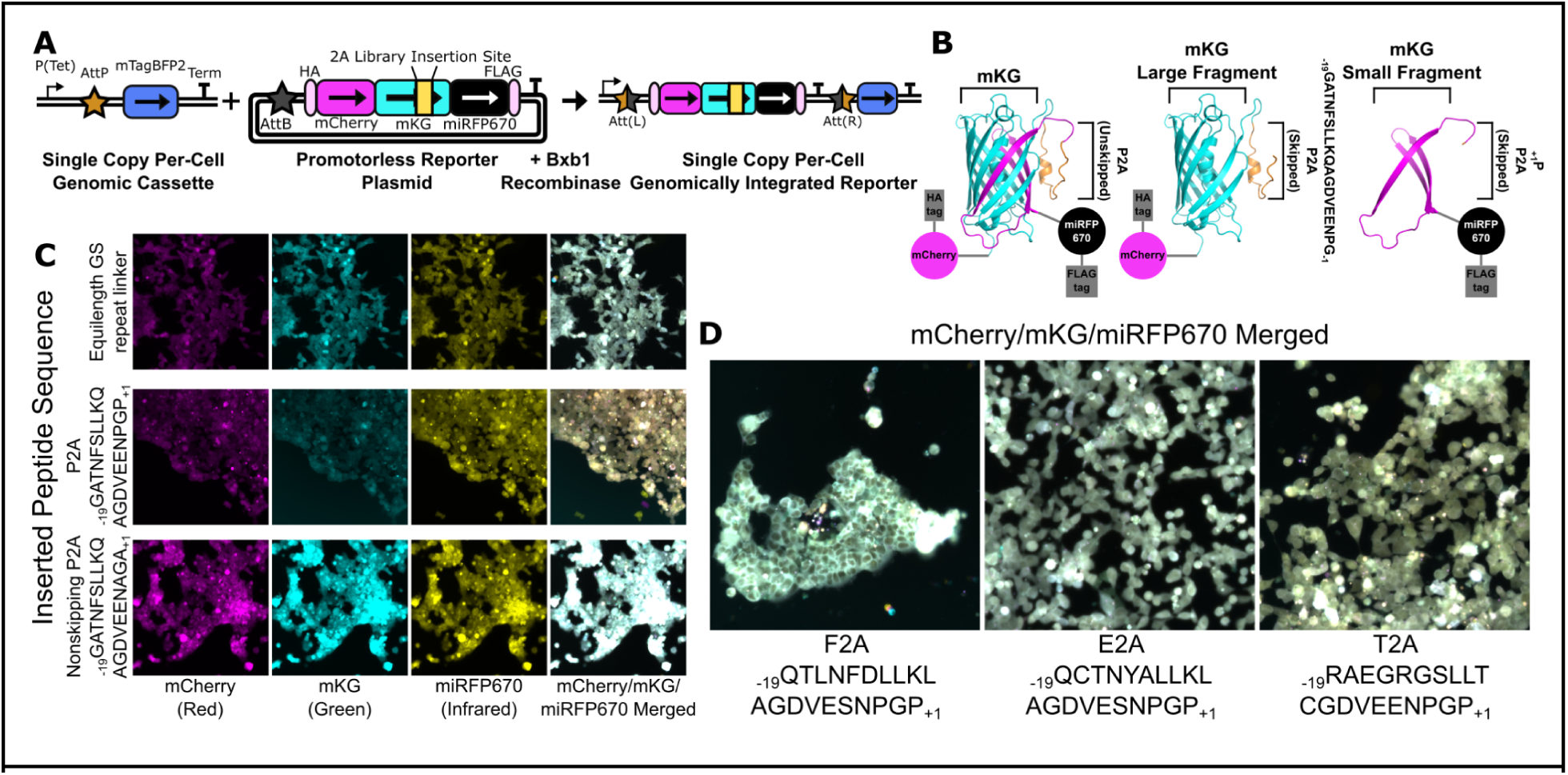
Design and Function of the Trifluorescent Reporter Assay System. (A) A single-site genomic integration ‘Landing Pad’ system for controlled expression of a Trifluorescent Reporter (TR) for study of 2A-like peptides. Utilization of promoterless Bxb1 attB plasmids and integration into an engineered site with a single promoter preceding a Bxb1 attP site ensures phenotypic effects only result from a single integrated plasmid copy per cell. (B) Representative diagram of the TR protein. Left: Structure of monomeric Kusabira Green (mKG) by AlphaFold 3 with 2A insertion site and P2A inserted into the flexible loop. Structure representative of failure to skip. Center: structure of mKG with P2A N-terminally resulting from either a skipping event or a falloff event. Right: structure of mKG with P2A C-terminally resulting from a skipping event. (C) Fluorescent images of HEK293T cells expressing the TR and containing an equilength GS repeat (top), P2A (middle) or nonskipping P2A (bottom) approximately 3 weeks post transfection. (D) Merged fluorescent images for TR containing F2A, E2A, and T2A.

By utilizing the known mutable flexible loop in mKG, the reporter was expected to allow for measurement of the different skipping outcomes (Figure 2B, Figure S1, S2). In the case of a readthrough event, the beta-barrel of mKG would not be disturbed as shown by AlphaFold 3 structural prediction*^27^*, and the protein products would be red, green, and infrared fluorescent.

In the case of a skipping event, 3 beta-strands of the beta-barrel of mKG remain attached to miRFP670, and the remainder of the beta-barrel is attached to mCherry, producing an infrared and red fluorescent protein, respectively. During a falloff event, the 3 beta-strands of mKG and miRFP670 are not produced, producing only a red fluorescent protein.

We constructed a set of TR plasmids containing a 20 residue GS repeat ((GS)x10), P2A, and a non-skipping version of P2A where the C-terminal _-3_NPGP_+1_ is mutated to _-3_NAGA_+1_. When transfected into HEK293T cells and maintained for 3 weeks, we observed fluorescence for all fluorophores, with a relative decrease in green fluorescence in P2A consistent with the occurrence of skipping (Figure 2C). We further transfected TR plasmids containing F2A, T2A, and E2A, their relative _-3_NAGA_+1_ nonskipping variants, and TR without an insert into cells to confirm fluorescence with a range of samples (Figure S3-S13). Both E2A and T2A operated similarly to P2A (Figure 2D). F2A and its _-3_NAGA_+1_ variant exhibited nuclear exclusion in all fluorescent channels (Figure S7, S11), which is consistent with recent reports*^28^* and likely results from _-17_LNFDLLKLAG_-8_ acting as a CRM1 recognition motif. Nuclear exclusion did not appear to affect overall fluorescent intensity.

### Efficacy of the Trifluorescent Reporter with Standard 2A Peptides

We first established that fluorescence could be used to measure ribosomal skipping with the TR (Table S4). We performed western blotting, the standard for measuring skipping, on P2A, T2A, E2A, F2A, the nonskipping _-3_NAGA_+1_ variants, the (GS)x10 equilength control, and the TR without an insert at the 2A insertion site (Figure 3A, S14). Blotting for the N-terminal HA tag identified the nonskipping band in the nonskipping controls, a mixture of skipping and non-skipping bands in E2A, F2A, and T2A, and predominantly the skipping band in P2A. When measuring the comparative intensity of the skipped and the nonskipped bands, we observe that P2A and T2A skipped in accordance with the prior results*^3^*, while F2A and E2A had marginally lower skipping than expected. When blotting for the C-terminal FLAG tag, we observed similar results for P2A and T2A, with E2A and F2A reporting skipping more akin to prior results.

**Figure 3.**
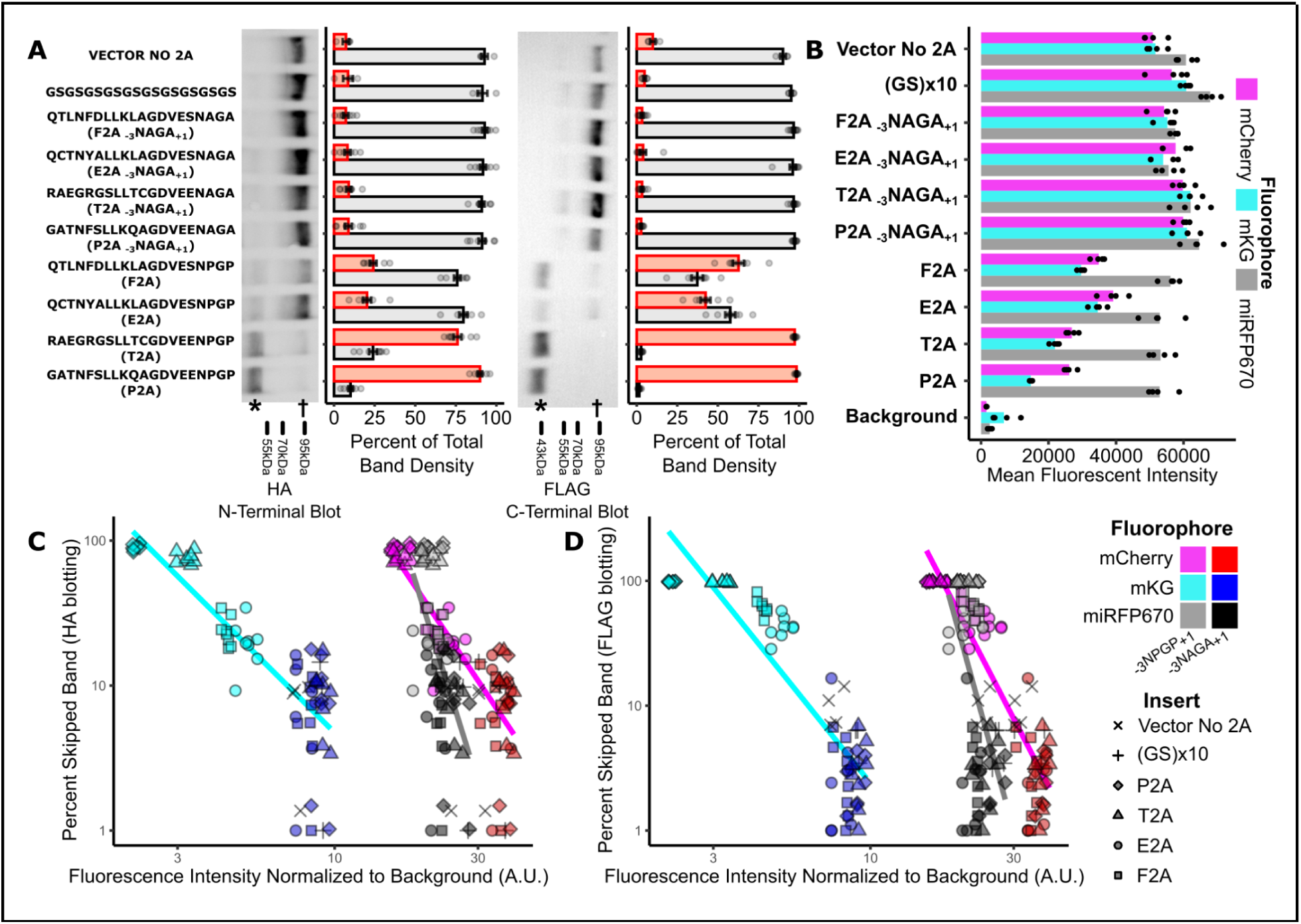
Trifluorescent Reporter Skipping Assay Efficacy. (A) Representative western blots for HA (N-terminal tag, left) and FLAG (C-terminal tag, right) in trifluorescent reporter controls. Relative band intensity of skipped band (orange) and unskipped band (grey) in repeat blots with independently transfected population represented in bar graphs. *, skipped protein bands. †, unskipped protein bands. (n = 4) (B) Geometric means of flow cytometry MFI measurements for each fluorophore in trifluorescent reporter for controls. (n = 4) (C) Relationship between fluorescence intensity (normalized to cell background) and percentage of skipped band as measured by blotting for HA N-terminal tag of the TR. Spearman ρ for mCherry: −0.815. Spearman ρ for mKG: −0.807. Spearman ρ for miRFP670: −0.560. (D) Relationship between fluorescence intensity (normalized to cell background) and percentage of skipped band as measured by blotting for FLAG C-terminal tag of the TR. Spearman ρ for mCherry: −0.842. Spearman ρ for mKG: −0.845. Spearman ρ for miRFP670: −0.487.

We next measured the Mean Fluorescence Intensity (MFI) of each construct by flow cytometry (Figure 3B, S15) and observed a relative decrease in mKG fluorescence concurrent with prior reported skipping functionality in P2A, T2A, E2A, and F2A. We also observed a concurrent decrease in mCherry fluorescence, indicating that mCherry fluorescence was acting as an unintended secondary indicator of skipping. There was minimal decrease in miRFP670 fluorescence in the skipping samples relative to non-skipping samples. We also measured fluorescence in bulk by plate fluorimetry, and observed similar results to the flow cytometry (Figure S16, Table S4).

We determined that fluorescent measurements approximated western blotting readouts for each construct. We first compared the MFI to the percentage of N-terminal skipping product (Figure 3C). We calculated a Spearman ρ of −0.815, −0.807, and −0.560 for mCherry, mKG, and miRFP670, respectively. When we compared MFI to the percentage of C-terminal skipping product (Figure 3D), we calculated a Spearman ρ of −0.842, −0.845, and −0.487 for mCherry, mKG, and miRFP670, respectively. This indicates that, while western blotting remains the higher fidelity measurement approach, fluorescence as measured in the TR is sufficiently capable of identifying the ribosomal skipping phenotype. We also tested an alternative TR design where mCherry and miRFP670 were swapped, and observed the same pattern of relative fluorescent changes (Figure S17, S18).

### Sort-Seq of Library of Non-Viral 2A-Like Peptides

Having established that fluorescent intensity in the TR changes proportionally to skipping efficiency, we proceeded to our high-throughput assay. We shuttled an oligopool library encoding our bioinformatically identified candidate non-viral 2A-like coding sequences, the previously tested samples as a control panel, a sizable number of synonymous variants of P2A, T2A, E2A, and F2A, and a small set of targeted mutants into the TR using Golden Gate Cloning*^24^*. Sequencing of the plasmid preparation indicated that 86% of the library was successfully shuttled into the TR vector.

The plasmid library was recombined into HEK293T “landing pad” cells^25,26^. We selected for recombined cells and grew the cells out for 3 weeks to ensure stable TR expression. We performed quartile sorts for each fluorophore, and Illumina sequenced each quartile to quantify the frequency of each library member in each quartile. We calculated a weighted average ‘Sort-Seq’ score for each library member using the read frequency*^29^* (Figure 4A). This was repeated across 7 experiments. 78% of the library had at least one Sort-Seq score for each fluorophore, and 68% of the library had at least two scores for each fluorophore (Table S5).

**Figure 4.**
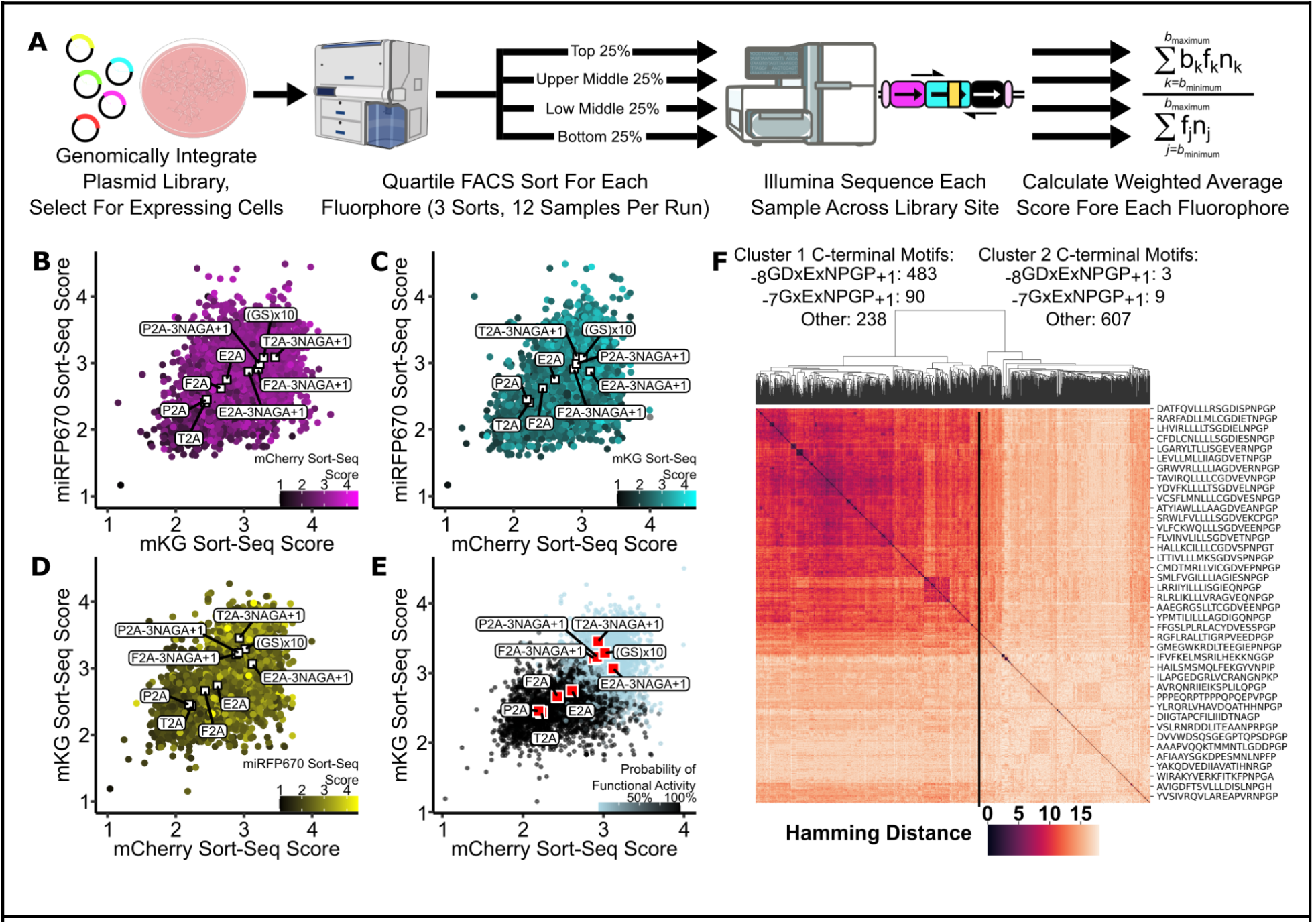
Sort-Seq and Initial Classification of Library. (A) Schematic representation of Sort-Seq experimental workflow. Cells were transfected with the plasmid library and selected for single-copy stable expression. The cells expressing the library were sorted into quartiles for each fluorophore. Each quartile was Illumina sequenced across the library site to determine library member frequency. A weighted average score was calculated for each library member for each fluorophore. (B, C, D) Sort-Seq results for library samples. In each pairwise comparison, skipping controls localized in the lower quadrant, and nonskipping controls localized in the upper quadrant. (E) Results of fuzzy-clustering machine learning algorithm to predict skipping classification using Sort-Seq scores. Light-colored points were classified as functionally inactive, while dark-colored points were classified as functionally active. (F) Identity matrix of samples that were classified as functionally active by the fuzzy-clustering algorithm. Most peptides containing the canonical and non-canonical C-terminal core motifs were within cluster 1, and most peptides containing divergent C-termini were within cluster 2.

We compared the Sort-Seq scores for the library to check for clustering of controls. We observed that the P2A, T2A, E2A, and F2A _-3_NAGA_+1_ nonskipping variants and (GS)x10 all clustered around a Sort-Seq score of 3 for each fluorophore. We also observed that P2A, T2A, E2A, and F2A observed proportionately decreasing Sort-Seq scores approximately in rank order based on the observed skipping functionality in the TR (Figure 4B,C,D, Figure 3A, Figure S19).

We trained a series of classification models to classify samples as functionally active or inactive based on the Sort-Seq scores using the controls as the training data (Figure S20). We utilized a Fuzzy Clustering (FC) model for final classification due to its reporting probability of class inclusion in the prediction and correctly reclassifying all controls. The model broadly determined that as Sort-Seq scores decrease, the probability that a peptide is functionally active increases (Figure 4E).

Peptides classified as functionally active frequently exhibited higher sequence identity than did the entire library (Figure 4F). Only two-fifths of the peptides classified as functionally active had either the canonical or non-canonical C-terminal core motifs. The diversity of peptides classified as functionally active indicates that there may be more ribosome-altering nascent peptides than previously thought.

### Impacts of Synonymous Codons on Canonical Skipping

The relative contribution of the nascent peptide compared with the mRNA or codons to ribosomal skipping remains unclear. Most reports show little impact of synonymous codon substitutions on skipping*^1,2,12,19^*, though others argue that targeted codon deoptimization can completely terminate skipping functionality*^20^*. To determine how codons impact skipping, we measured sub-libraries for P2A, T2A, E2A, and F2A encoding synonymous codons at each position, as well as a fully recoded variant where every codon was replaced with a synonymous codon.

When we compared the Sort-Seq scores of the synonymous variants with the parent coding sequences, we observed that synonymous variants clustered around centroids near the parent coding sequences (Figure 5A, Table S5). We identified each centroid and determined the distance in standard deviations for each sample relative to the centroid. Most single-codon synonymous variants lay within three standard deviations of the centroid (Figure 5B). The fully recoded variants were toward the maximal number of standard deviations from the centroid, while the parent coding sequences were approximately one standard deviation from the centroid. Nonskipping _-3_NAGA_+1_ variants were the greatest number of standard deviations from the centroid. We did not observe any positional or codon specific patterns that influenced the distance of synonymous variants from the population centroid (Figure S21).

**Figure 5.**
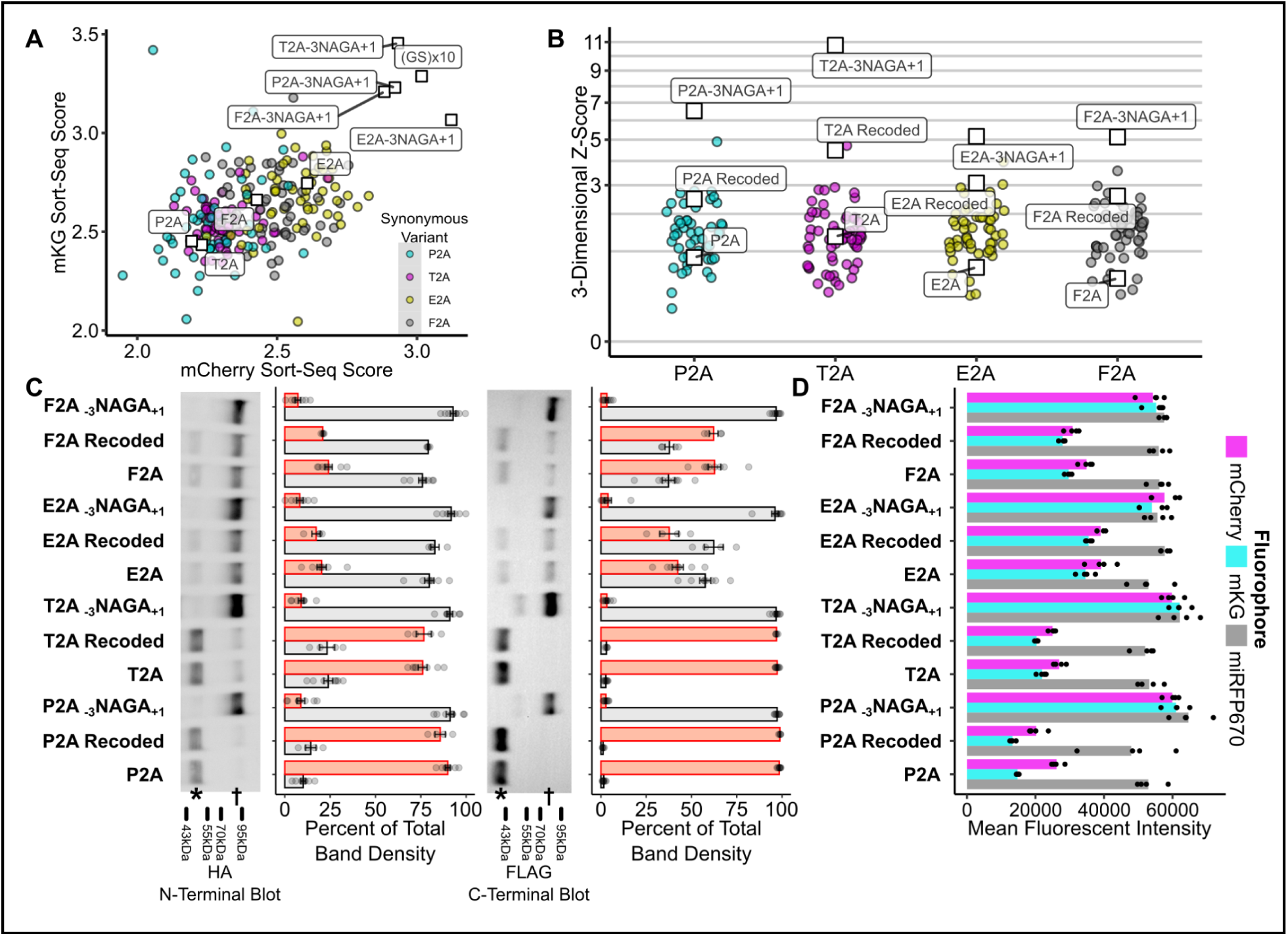
Effects of Synonymous Variants on Ribosomal Skipping. (A) Distribution of Sort-Seq scores of synonymous variants of P2A, T2A, E2A, and F2A. (B) Number of standard deviations from the centroid of the population of synonymous variants. (C) Representative western blots for HA (N-terminal tag, left) and FLAG (C-terminal tag, right) for control coding sequence and coding sequences where each codon is a synonymous codon for P2A, T2A, E2A, and F2A. *, skipped protein bands. †, unskipped protein bands. (n = 4) (D) Geometric means of flow cytometry measurements for each fluorophore for each sample. (n = 4)

Having not observed any variant spread indicative of changes in skipping functionality, we individually generated clonal lines expressing the fully recoded variants of P2A, T2A, E2A, and F2A. We observed that the recoded variants were indistinguishable from the parent coding sequences by microscopy (Figure S22-S25). We measured skipping for the recoded variants by western blotting (Figure 5C, Figure S26) and flow cytometry (Figure 5D, Figure S27, S28, Table S4). We observed little, if any, difference between the recoded variants and the parent coding sequences.

Having observed no difference in the ribosomal skipping efficiencies of peptides with synonymous mutations, we attempted to replicate the prior reported loss of skipping function result. The report was that P2A lost the ability to skip when the _-3_NPGP_+1_ coding sequence was mutated from AACCCTGGACCT to AATCCGGGTCCG*^20^*. We were unable to replicate the reported loss in skipping (Figure S29). Our results support peptide sequences, rather than mRNA sequence and codon choice, being the driver of 2A-mediated ribosomal skipping.

### Evidence for Active 2A-like Peptides in Eukaryotes

Ribosomal skipping sequences function in diverse eukaryotic contexts including yeast, plant, insect, and mammalian cells, but do not function in bacterial cells*^1,3,15^*. Unlike traditionally monocistronic eukaryotic mRNA translation, bacterial mRNA is often polycistronic with multiple internal translation initiation sites. Furthermore, the structure, size, and shape of bacterial and eukaryotic ribosomes differ significantly*^30,31^*. Bacterial proteins are unlikely to experience selective pressure for functionally active 2A-like ribosomal skipping sequences akin to eukaryotes, as any such peptide would likely be incapable of skipping in its native context. Bacterial sequences consequently represent a stochastic distribution of 2A functionality.

We experimentally tested approximately three times as many eukaryotic-origin peptides than bacterial-origin peptides (Figure 6A), though this likely reflects the dominance of eukaryotic genomes in the queried NCBI databases. Despite their identification by identical search criteria, we observed differing distributions of Sort-Seq scores between eukaryotic and bacterial peptides (Figures 6B,C). A greater proportion of eukaryotic peptides exhibited lower Sort-Seq scores than did bacterial peptides (Figure 6D). Accordingly, a greater proportion of eukaryotic peptides were classified as functionally active (Figure 6E). Considering the collective lack of evidence for 2A-like peptide functionality in bacteria, this supports the hypothesis that there are evolutionary pressures surrounding active 2A-like peptides in eukaryotic proteins.

**Figure 6.**
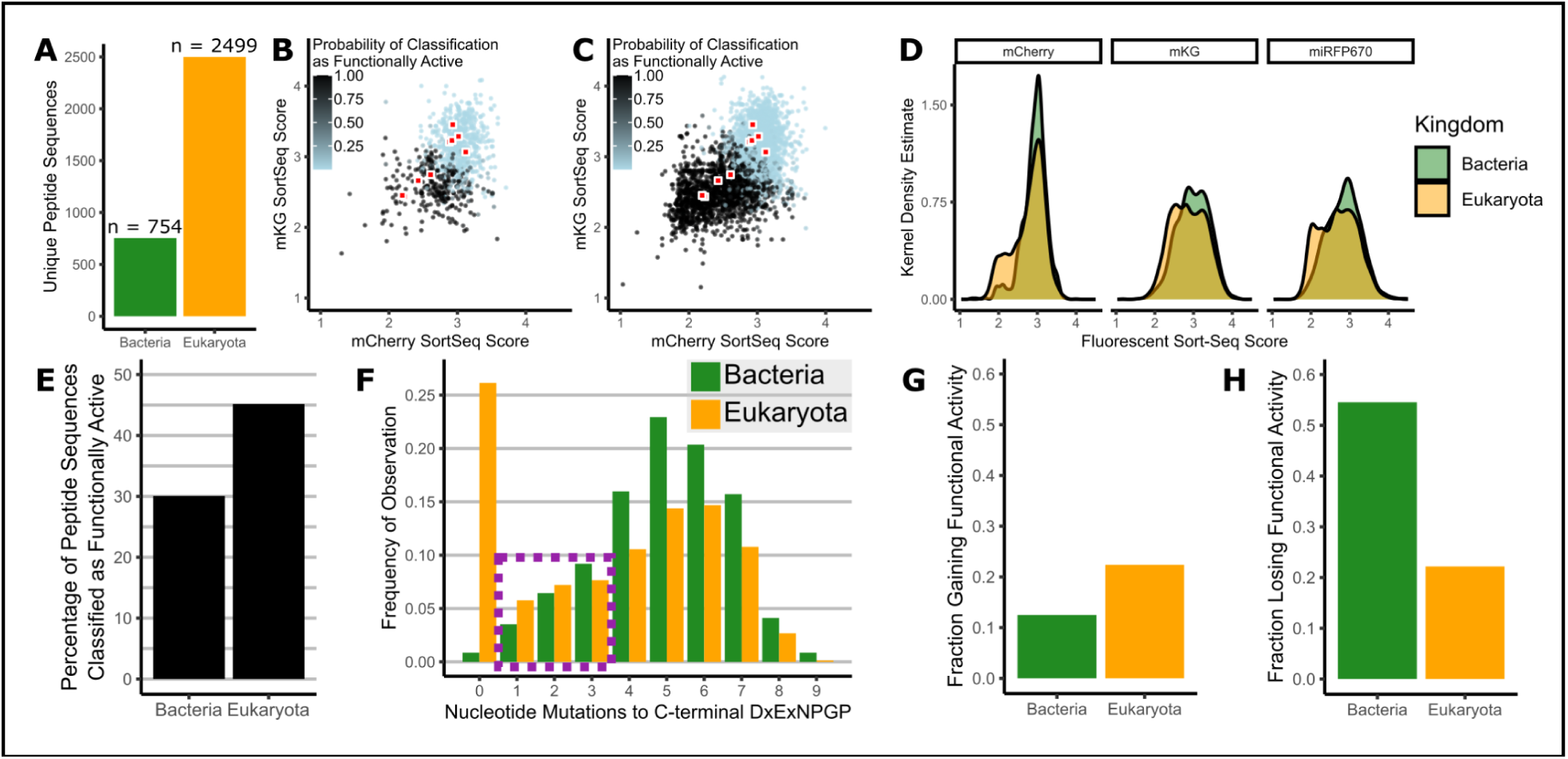
Active 2A-like Peptides are More Prevalent in Eukaryotes than Bacteria. (A) Difference in number of bacterial and eukaryotic peptides tested by Sort-Seq. Scatter plots of mCherry and mKG Sort-Seq scores and classifications for activity for (B) bacterial and (C) eukaryotic peptides. (D) Kernel density estimates of Sort-Seq score distributions for bacterial and eukaryotic-origin peptides. (E) Bar plot of percentage of 20-mer peptide sequences classified as functionally active in bacteria and eukaryotes. (F) The minimum number of substitution mutations needed to generate the C-terminal _-7_DxExNPGP_+1_ motif, excluding peptides with C-terminal _-7_GxExNPGP_+1_. Coding sequences with 3 or fewer nucleotide changes to generate the C-terminal _-7_DxExNPGP_+1_ motif (dotted box) were generated and tested by Sort-Seq to capture the lower mode in eukaryotic peptides. (G) Fraction of peptides mutated to generate _-7_DxExNPGP_+1_ that went from functionally inactive to functionally active classification upon mutation. (H) Fraction of peptides mutated to generate _-7_DxExNPGP_+1_ that went from functionally active to functionally inactive classification upon mutation.

There were differences in the 2A-like C-terminus depending on genomic origin. Eukaryotic peptides more frequently contained the _-7_DxExNPGP_+1_ canonical C-terminal core motif than did bacterial peptides (Figure 6F). We identified the shortest evolutionary path in substitution mutations that would generate _-7_DxExNPGP_+1_. We determined that there was a mode among non-_-7_GxExNPGP_+1_ eukaryotic coding sequences at 6 mutations, and at 5 mutations for bacterial coding sequences. A greater proportion of eukaryotic coding sequences were proximal to the canonical _-7_DxExNPGP_+1_ core motif and only deviated from it by 3 or fewer nucleotides (Figure 6F).

This relative enrichment of eukaryotic _-7_DxExNPGP_+1_-proximal coding sequences may implicate evolutionarily accessible functional activity. To test this, for all coding sequences that differed from _-7_DxExNPGP_+1_ by 3 or fewer nucleotides, we generated a paired artificially-derived coding sequence where the _-7_DxExNPGP_+1_ was constituted through the fewest number of substitution mutations. This sublibrary of constituted peptides was tested via Sort-Seq and paired with the extant 2A-like peptides.

Mutations to the _-7_DxExNPGP_+1_ motif had no effect on the FC classification for three-fourths of peptides of either bacterial or eukaryotic origin. A greater proportion of inactive peptides of eukaryotic origin gained activity upon mutation as compared to bacteria (Figure 6G). Conversely, a smaller proportion of active peptides of eukaryotic origin became inactive upon mutation (Figure 6H). We used McNemar’s test to test the likelihood of functionally active peptides losing active classification and functionally inactive peptides gaining active classification being the same in both domains. We calculated that p=0.083 for all peptides, p=0.033 for eukaryotic peptides, and p=0.513 for bacterial peptides. These results indicate that extant eukaryotic peptides are more likely to be capable of mutating in to functionally active peptides than are bacterial peptides. Such peptides could become functionally active with evolutionary selective pressures.

### Phenotypic Diversity of 2A-Like Peptides

We individually tested a diverse subset of the identified 2A-like peptides to verify the Sort-Seq results. We selected 11 library members that were frequently observed in the Sort-Seq data and classified as functionally active, alongside 10 additional peptides from the library. We also utilized relative residue frequencies in classified functionally active peptides to design a novel 2A-like peptide, AFCFFLLLMLLCDVEINPGP, which we anticipated would be capable of skipping.

We created clonal cell lines expressing these 22 novel peptides. We first examined fluorescence via microscopy, plate-based fluorimetry, and flow cytometry (Figure S30-S38). We observed high correlation between individual fluorescent measurements and Sort-Seq scores (Figure 7A) and determined that the multiplex assay effectively captured the fluorescent phenotypes.

**Figure 7.**
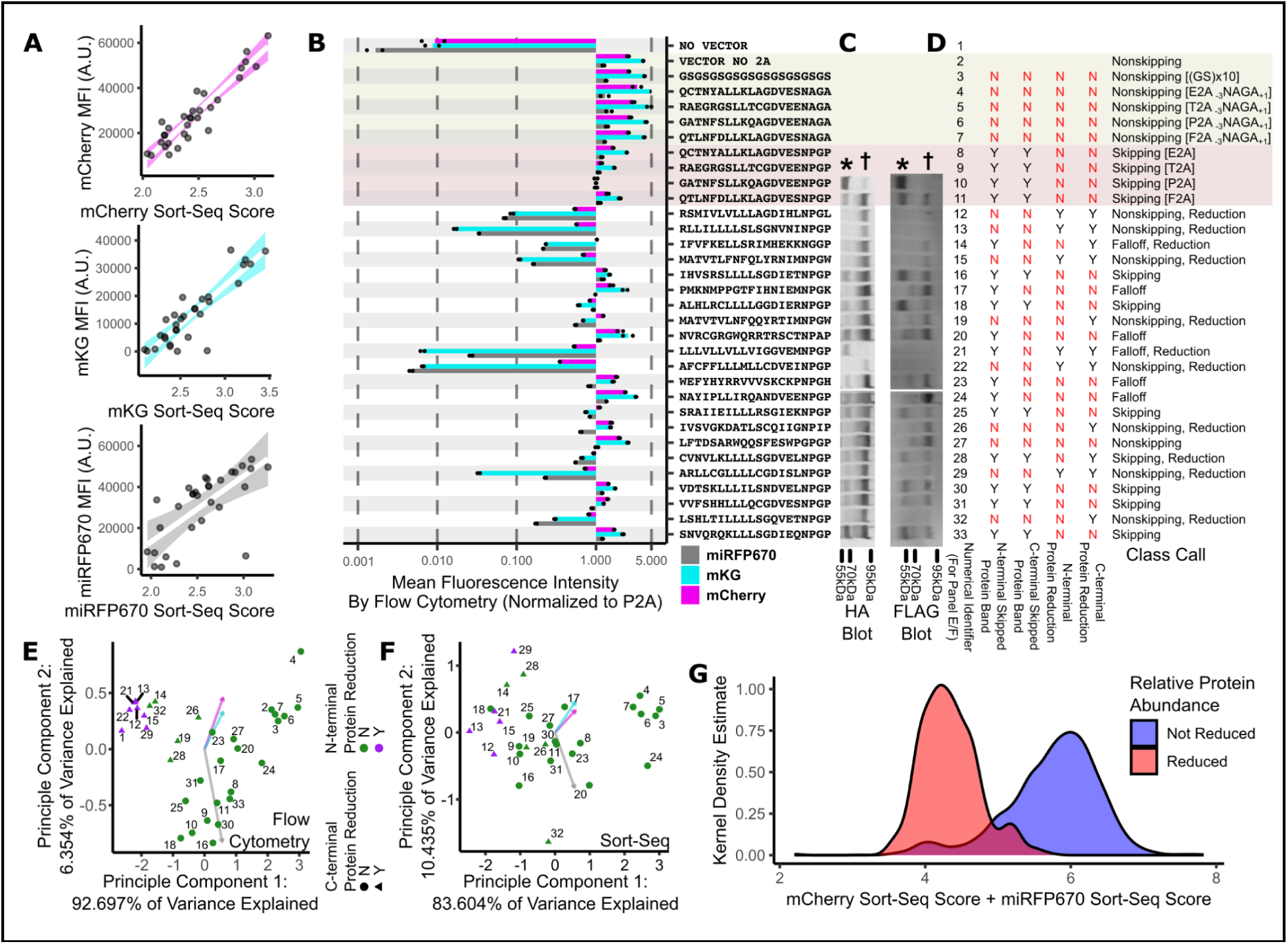
Individual Testing of Select 2A-like Peptides and Reclassification of the Sort-Seq Library. (A) Direct comparison of Sort-Seq Scores and MFI for controls and individually tested samples. Pearson r for mCherry: 0.93; Pearson r for mKG: 0.89; Pearson r for miRFP670: 0.70. Colored bands represent 95% confidence intervals. (B) Flow cytometry results for samples that were individually tested, normalized to P2A. (C) Western blot results for individually tested samples. Left: N-terminal HA tag. Right: C-terminal FLAG tag. *, skipped protein bands. †, unskipped protein bands. (D) Manual classifications made for functional 2A-like peptide phenotypes. Presence of N-terminal skipped protein band (HA), C-terminal skipped protein band (FLAG), N-terminal protein reduction (80% or less of P2A mCherry fluorescence), and C-terminal protein reduction (80% or less of P2A miRFP670 fluorescence) were identified as a binary for each peptide. Presence of no skipped bands determined nonskipping, presence of only N-terminal skipped bands determined falloff, and presence of both N-terminal and C-terminal skipped bands determined skipping. Either N-terminal or C-terminal protein reduction determined a reduction call. All samples were reclassified as one of the appropriate 6 classes based on these observations. (E) Principal component analysis of MFI from flow cytometry allows for separation of all protein reduction samples. (F) Principal component analysis of Sort-Seq scores for individually tested samples and controls. (G) Density plot of samples grouped in reduction classes and non-reduction classes by SVM reclassification models, plotted for their summed mCherry and miRFP670 Sort-Seq scores.

The novel peptides yielded a wide range of fluorescent values distinct from the skipping and nonskipping controls (Figure 7B). Eight peptides exhibited fluorescence values within the range of the controls, while the remaining thirteen had at least one color below that of P2A. Of these, eight peptides exhibited greater than 3-fold reductions to green and near-infrared fluorescence relative to P2A alongside mild reductions to red fluorescence.

We again utilized western blotting to test for the presence of skipped protein fragments using the N-terminal HA and C-terminal FLAG tags. Only 7 of the tested peptides exhibited classical skipping functionality indicated by the presence of ~50 kDa fragments in both blots (Figure 7C). We observed six peptides where only the N-terminal ~50 kDa band was present, which we reclassified as “falloff” peptides. This included peptides like LLLVLLVLLVIGGVEMNPGP, which only had the N-terminal skipped band, and WEFYHYRRVVVSKCKPNPGH, which had a weak N-terminal skipped band alongside unskipped bands, with neither of these demonstrating C-terminal skipped bands. Akin to the nonskipping controls, we observed nine inactive peptides that only exhibited the ~100 kDa full-length bands, which we reclassified as “nonskipping”.

Despite using pure populations of nearly isogenic cells and fractionating similar amounts of total protein through SDS-PAGE, we observed reductions in band intensities that mirrored the reductions in fluorescence intensity. To accommodate this unanticipated reduced protein phenotype, and due to the difficulties of cross-sample comparisons via western blotting, we labeled peptides inducing protein reduction based on fluorescence measurements. We defined N-terminal protein reduction as 80% of the mCherry fluorescence of P2A, and C-terminal reduction as 80% of the miRFP670 fluorescence of P2A (Figure 7D). This definition was used because P2A is the best known ribosomal skipping sequence, and a 20% reduction in fluorescence in either mCherry or miRFP670 relative to P2A would be beyond any expected skipping-induced decreases in a better performing skipping peptide. Half of the tested samples exhibited reduced protein abundance by this definition. Thus, combined phenotypic analysis revealed an unexpected range of phenotypic diversity, including alternative fragment accumulations and overall reductions to protein steady-state amounts, resulting in six distinct phenotypic classes (Figure 7D).The individual peptide reclassification scheme required protein fractionation and western blotting, which is not possible in multiplex. While skipping outcomes directly affect fluorescence, the TR was not designed to accommodate the reductions in protein abundance, which now had to be disentangled. Principal component analysis of the individual samples using MFIs from flow cytometry was sufficient to isolate samples exhibiting both C-terminal and N-terminal protein reduction (Figure 7E). Principal component analysis with the Sort-Seq scores of the same samples was also capable of separating peptides based on reduction, albeit less effectively (Figure 7F).

To better phenotypically describe the peptides tested in the library, we trained a support vector machine (SVM) model to classify the library into the aforementioned 6 classes using only their Sort-Seq scores. For simplicity, we collapsed N-terminal and C-terminal reduction into a broad ‘reduction’ subclassification, since N-terminal reduction always co-occurred with C-terminal reduction (Figure 7D). We condensed synonymous variants, and obtained a new classification for each peptide (Table S6). The model accurately reclassified 86% of samples used to train it. The SVM model successfully distinguished the classes exhibiting reduction, as the reduced classes had a modal combined mCherry and miRFP670 Sort-Seq score of approximately 4, as opposed to non-reduction classes having a modal score of approximately 6 (Figure 7G). While the phenotype of any specific 2A-like sample should be confirmed individually, the general accuracy of our classifications enables broader analyses of the characteristics and properties underlying each phenotypic group.

### Sequence Characteristics of Phenotypic Classes

The N-terminal residues of 2A-like peptides are postulated to strongly affect ribosomal skipping efficacy*^1,11,12^*. The sequence diversity of historically tested peptides has been insufficient to properly explore what characteristics of the 11 residues preceding the canonical C-terminal core motif potentiate ribosomal skipping. Having experimentally tested a diverse set of extant peptides that differ in functional outcome, we examined what amino acid differences may underlie the differential phenotypes.

Skipping and reduction class samples from eukaryotes share positional residue bias trends with viral peptides. We calculated and compared residue frequencies at each position among the phenotypically classified eukaryotic peptides with bacterial peptides and the viral peptides that were removed in the bioinformatic search. The skipping and reduction class samples in eukaryotes hierarchically clustered with the viral peptides, and the nonskipping and falloff samples from eukaryotes clustered with bacteria (Figure 8A). This indicates a commonality between the reduction and classical ribosomal skipping phenotypes.

**Figure 8.**
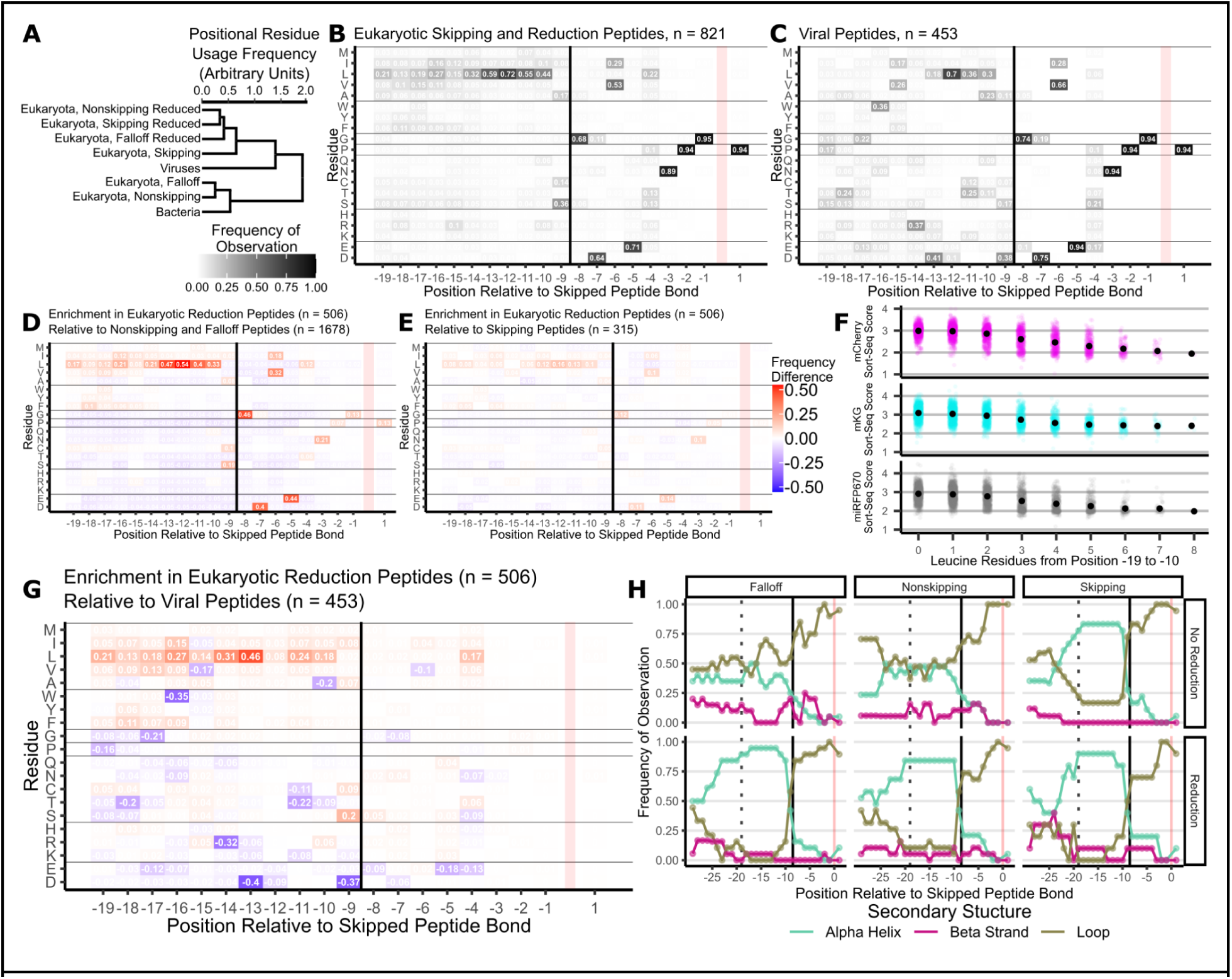
Functional Eukaryotic 2A-like Peptides Exhibit Distinct N-terminal Leucine Enrichment and Alpha-Helical Secondary Structure. (A) Hierarchical Clustering of peptide classes based on positional frequencies of residues. (B) Positional frequencies of residues in eukaryotic skipping and reduction class peptides. (C) Positional frequencies of residues in 2A-like viral peptides removed during bioinformatic search. (D) Residue enrichment of eukaryotic classified reduction peptides relative to eukaryotic classified nonskipping and falloff peptides. (E) Residue enrichment of eukaryotic classified reduction peptides relative to eukaryotic classified skipping peptides. (F) Relationship between number of leucine residues from position −19 to −10 and Sort-Seq score. Pearson r for mCherry: −0.606. Pearson r for mKG: −0.490. Pearson r for miRFP670: −0.495. (G) Residue enrichment of eukaryotic classified reduction peptides relative to 2A-like viral peptides removed during bioinformatic search. (H) Local secondary structure of eukaryotic 2A-like peptides across all 6 phenotypic classes. Dotted vertical line marks the most N-terminal residue tested (position −19 relative to skipped peptide bond). Vertical black line marks the N-terminus of the canonical core motif. Vertical red line marks the skipped peptide bond.

We observed an expansion of hydrophobic, predominantly leucine residues in the N-terminal region of eukaryotic skipping and reduction class peptides (Figure S41, S42). The skipping class and all reduction classes showed similar residue usage patterns, with preference for the canonical C-terminal core motif and enrichment of leucine residues from position −13 to position −10, and a general enrichment of hydrophobic residues from position −19 to −10 (Figure 8B, Figure S39). Eukaryotic nonskipping and falloff peptides showed slight preference for the canonical C-terminal core motif with limited enrichment in the N-terminal region (Figure S39). Only 14.86% of tested bacterial peptides were classified as skipping or in a reduction class, limiting interpretation of their sequence characteristics (Figure S40).

Viral peptides that were removed in the bioinformatic search had a similar residue usage pattern to the skipping and reduction samples in Eukaryotes (Figure 8C). Though we did not experimentally test the viral peptides for functionality, we expect they will exhibit similar frequency of functionality as prior reports, and are thus a sufficient proxy for the residue usage patterns of viral ribosomal skipping sequences. We observed no enrichment of the proposed ‘Class B’ 2A-like peptides^15^. The reported hallmark, _-15_W, showed almost no enrichment signal, only being present in 27 of the 2,499 unique tested eukaryotic peptides (Figure S39, S40). Such ‘Class B’ peptides appear to be rare in eukaryotic 2A-like peptides from our results.

We compared residue use patterns to identify enrichments which may contribute to the differences between classes. When we compared the pooled reduction classes with the pooled nonskipping and falloff classes, we observed an enrichment of residues associated with the canonical and noncanonical C-terminal core motifs and of N-terminal hydrophobic residues alongside leucine (Figure 8D). A nearly identical pattern was observed when the skipping class was compared with the pooled nonskipping and falloff classes. We observed minimal positional enrichment of residues when comparing the pooled reduction and skipping classes (Figure 8E).

Leucine was globally enriched in active 2A-like sequences, so we determined how the number of leucine residues between position −19 to −10 impacted Sort-Seq scores for all sequences. We observed that when there were 3 or more leucine residues in the N-terminus, the Sort-Seq score decreased, with Pearson r of −0.606, −0.490, and −0.495 for mCherry, mKG, and miRFP670 respectively (Figure 8F).

The leucine expansion is likely a differentiator between skipping and reduction. Since 2A-like viral peptides are largely expected to skip and reduction has not been previously reported to be a prominent viral phenotype, differences between the eukaryotic reduction peptides and viral peptides should identify the boundary between skipping and reductions in protein abundance. We thus compared residue usage among the pooled reduction classes with the viral 2A-like peptides (Figure 8G). Hydrophobic residues were N-terminally enriched in eukaryotic reduction peptides, with a striking enrichment of _-13_L and _-14_L relative to viruses. In contrast, viruses more frequently utilized a broader range of residues in the N-terminal region, including _-16_W, _-14_R, and _-13_D. The enrichment of hydrophobic residues, and leucine specifically, appears to determine reduced protein abundance, with viral peptides containing 1-3 leucines, and eukaryotic peptides that induce protein reduction containing 3-6 leucines (Figure 8F, Figure S42). As throttling overall protein expression would likely be evolutionarily disadvantageous for many viruses, the residue differentiation we observe is most likely driven by viral evolutionary pressure to avoid reducing viral protein expression.

The N-terminus of 2A-like peptides is considered to form an alpha-helix in the ribosomal exit tunnel*^2,15,19^*, but where the alpha helix typically begins and ends relative to the skipped peptide bond has not been clearly defined. We considered the local secondary structure of 2A-like peptides in the context of their native proteins to identify the boundaries of the alpha helix. We identified 267 unique eukaryotic proteins and 77 unique non-eukaryotic proteins with 2A-like peptides that were well-represented in the Sort-Seq data. We utilized AlphaFold 3 to predict those structures*^27^*, and implemented DSSP*^32^* to identify the predicted local secondary structure of the 2A-like peptides (Figure 8H, Figure S43, Table S7). Among eukaryotic peptides, we observed that the reduction and the skipping classes showed strong propensity for alpha helices from position −19 to −10 and for free loops from position −9 to +1. Alpha helices began between position −25 and −20. The nonskipping and falloff classes showed more mixed distribution of alpha helices and free loops from position −19 to −10. The regularity of position −10 being the cutoff for the alpha helix is likely mechanistically important for reduction and skipping.

The consistent hydrophobic residues along the N-terminal alpha helix indicates that specific geometric presentation of these residues contributes to both skipping and reduction. The positioning of leucines from position −14 to −10 on the putative alpha-helix likely induces specific interactions with the ribosomal exit tunnel necessary for both skipping and reduction. Paired with the prior observation that removal of N-terminal residues hinders skipping*^1,10,15^*, these data indicate that formation of an alpha-helix with specific positional hydrophobic residues in the exit tunnel is mechanistically necessary to prime the ribosome so that skipping can occur. Increased leucines at the N-terminal regions of the helix appear to preferentially bias the translated protein to become reduced in abundance.

### 2A-like Peptides in Eukaryotic Protein Contexts

While the utility of protein 2A in viruses is well understood, the functional role of 2A-like peptides in eukaryotic proteins remains unclear. There are prior reports of functionally active 2A-like peptides in select eukaryotic contexts, such as APE-type non-LTR retrotransposons*^4^* and NOD-like receptor proteins*^33^*. Most proteins are first identified en masse through genome sequencing and automated annotation, resulting in minimal functional or structural data and little contextual information of their roles. The majority of studies that consider the native context of 2A-like peptides also predate the major protein structure prediction models*^34^*, further limiting past structural insights and data annotation for proteins with 2A-like peptides. Within our dataset, most proteins still retained ambiguous designations, limiting interpretation of native 2A-like peptide functionality.

Having already predicted structures for 344 proteins, we determined that 2A-like peptides occupy a wide variety of protein structural families. Following AlphaFold 3 predictions of the protein tertiary structures, we used Foldseek*^35^* to perform structural database similarity searches. This allowed us to group proteins based on identifiable domains within their predicted tertiary structures (Figure S44). The structures grouped into 75 distinct annotations. 11 annotations were represented by 3 or more unique 2A-like peptides (Figure 9A, Table S7).

**Figure 9.**
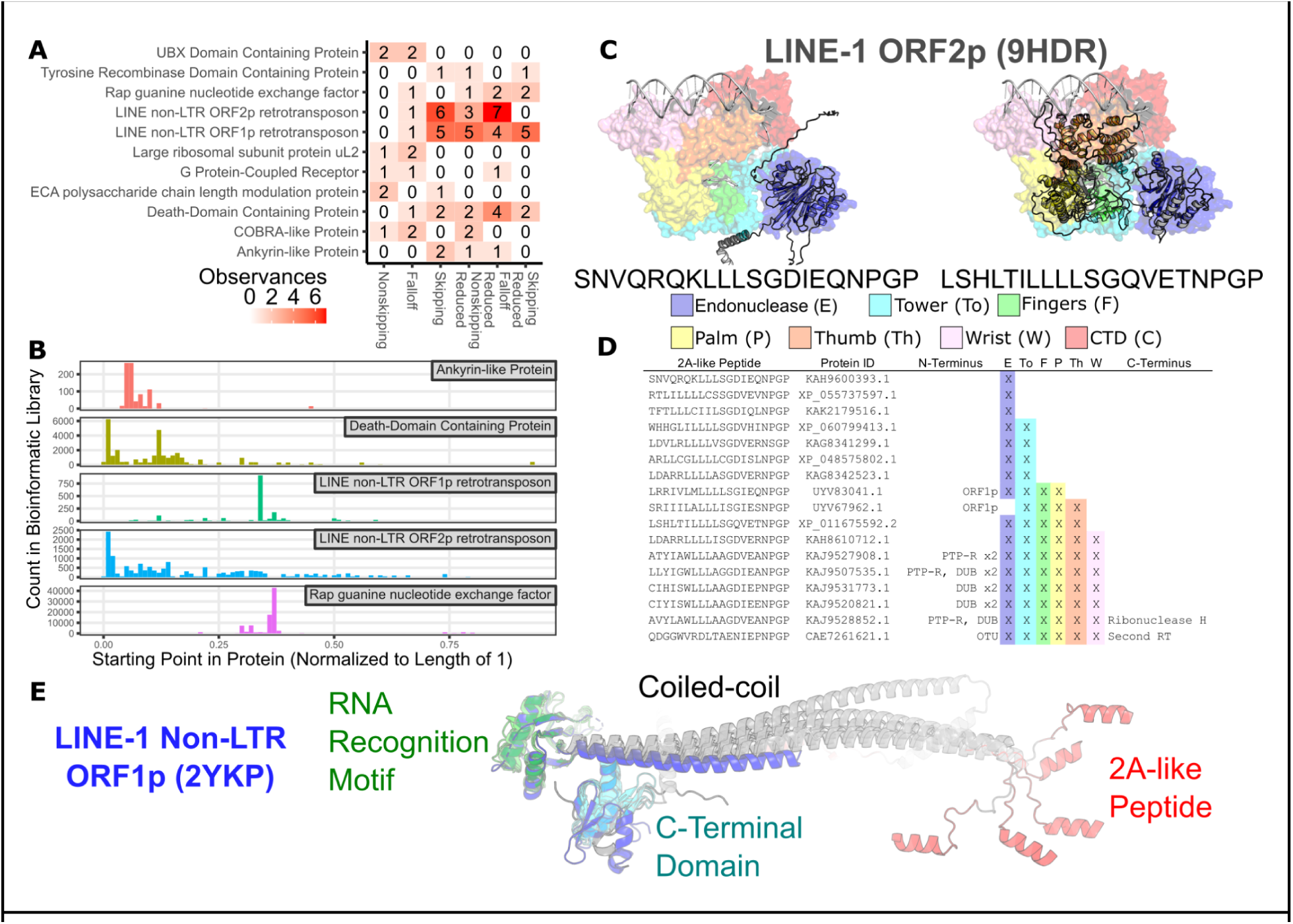
Functional 2A-like Peptides are Associated with Various Genes and Transposable Elements. (A) Structure annotations that were observed 3 or more times among AlphaFold 3 structural predictions of 344 full-length proteins containing 2A-like peptides. (B) Histogram of the relative position of residue −19 of 2A-like peptides in the 5 most enriched structure annotations. (C) Representative structures of LINE non-LTR ORF2p-like proteins (left: KAH9600393.1, right: XP_011675592.2) carry differential amounts of the reverse transcriptase domains when compared to LINE-1 ORF2p (9hdr). (D) Table of structural domains present in LINE non-LTR ORF2p proteins as described in (A). DUB: Deubiqutinase; PTP-R: Pentacotripeptide-Repeat; OTU: OTU Deubiquitinase. (E) Representative structures of LINE-1 ORF1p-like proteins aligned with LINE-1 ORF1p (2ykp). Overlaid structures are: GFS00708.1, KAH9368516.1, XP_037274885.1, XP_037556826.2, XP_042143744.1, and XP_042149113.1.

Of the well represented annotations, 5 stood out for being enriched for skipping and reduction classifications. 2A-like peptides associated with Rap Guanine nucleotide Exchange Factor (RapGEF), death-domain containing proteins, and ankyrin-like proteins were related to autosomally encoded proteins, and 2A-like peptides associated with LINE1 non-LTR ORF1p and ORF2p proteins were instead associated with genes from transposable elements. We observed that 50% or more of 2A-like peptides were in a reduction class for each of these annotations (Figure 9A). When we returned to the full bioinformatic dataset (Table S2), we observed that peptides of the same annotation were regularly in the same position within their respective proteins (Figure 9B).

The structural annotations we identified reflected patterns in annotated proteins in the greater bioinformatic dataset. There, we observed 1 cluster (here defined as hamming identity distance of 3 or less) of 2A-like peptides in RapGEF across 34 species of teleost fish. The singular protein family and high relatedness of these peptides indicates that they may originate from a shared acquisition event. We also observed 1 cluster of 7 peptides in ankyrin-like proteins in *Halichondria panicea* and 4 clusters in death-domain containing proteins among 4 unrelated marine species, including 2 peptides in NOD-like receptors in *Strongylocentrotus purpuratus*, one of which was observed prior^33^. The ankyrin-like proteins may share common ancestry, but it is not immediately obvious how evolutionarily inter-related either of these annotations is. Importantly, due to the large number of unannotated proteins in the bioinformatic dataset, our results here both for structural annotations we made and for analysis based on existing annotations only represents a fraction of the identified proteins containing 2A-like peptides.

Diverse non-LTR retrotransposons proteins were linked to both skipping and reduction class 2A-like peptides. Functional 2A-like peptides in ORF2p-like reverse transcriptase (RT) enzymes were previously identified in the Crack, L2, L2A, CR1, Ingi, and L1Tc clades^4^, which all possess an N-terminal APE-type DNA endonuclease domain. We likewise observed ORF2p-like proteins with 16 distinct 2A-like peptide clusters across 16 different species, ranging from algae to fish. Most instances of ORF2p we observed included the APE-type endonuclease domain, but had differing amounts of the subsequent RT domains and preceding N-terminal sequence(Figure 9C, D), resulting in more variable 2A locations in the predicted protein product (Figure 9B), though the 2A-like peptide regularly preceded the endonuclease domain. Approximately half of instances were sufficiently complete to be capable of self replication.

Some complete instances of ORF2p were fused to other functionally meaningful protein domains such as deubiquitinase domains (DUBs). Within *Haematococcus lacustris*, we observed five unique 2A-like peptides for ORF2p including those from the distant Dualen/RandI clade^36^. Among these, two had N-terminal Josephin DUB domains, one had an N-terminal pentacotripeptide repeat domain, and two had both. We separately observed a *Symbiodinium necroappetens* ORF2p with an N-terminal OTU DUB domain, alongside a C-terminal fusion of a second RT domain. In the bioinformatic dataset, we observed 16 clusters of 2A-like peptides across kingdoms, including plants and animals, indicating likely repeated independent acquisitions.

There were nine distinct 2A-like peptide clusters (hamming identity distance of 3 or less) with non-LTR retrotransposon ORF1p-like nucleic acid binding proteins across one species of limpet and five different ticks. These proteins encode a coiled-coil domain, an RNA-recognition motif, and a C-terminal domain of unclear function. The 2A-like peptides always preceded the coiled-coil domain (Figure 9E). Despite low sequence similarities, the predicted structures of these proteins were nearly identical, and the locations of the 2A peptides were largely consistent (Figure 9B).

2A-like peptides also appeared in more novel and unexpected retrotransposon forms. We observed two instances of 2A-like peptides between N-terminal ORF1p and C-terminal ORF2p domains in *Cordylochernes scorpioides*, and the APE-type domain only appeared to be present in one of these (UYV83041.1). We also found a 2A-like peptide attached to the C-terminal end of a Ty3-like LTR retrotransposon reverse transcriptase. Altogether, the majority of retrotransposon-associated 2A-like peptides exhibited reduced protein abundance, making up 70% of ORF1p occurrences and 59% of ORF2p occurrences we tested, indicating that reduction may be directly related to the primary function of 2A-like peptides in retrotransposons.

2A-like peptides are found in both autosomal proteins and retrotransposons within diverse eukaryotic organisms. They exhibit a range of skipping outcomes, but also frequently exhibit reduced protein abundance. The properties and secondary structure along the nascent peptide sequence distinctly parallel what is seen in viral 2A peptides. Altogether, it appears that 2A-like peptides in eukaryotic proteins natively operate as specialized regulatory motifs that regulate protein expression through the induction of peptide skipping, protein reduction, or both.

## Discussion

In this study, we developed the TR, bioinformatically identified 4,218 non-viral 2A-like peptides, experimentally tested 3,271 of these peptides, and found that 821 exhibited 2A-like mechanistic phenotypes. We identified that a large number of peptides had reduced protein abundances. The combined consideration of both reduction and skipping allowed us to identify that 2A-like peptides undertake a 10-residue alpha-helical secondary structure followed by 10 residues of free loops. We identified that specific positional leucine residues appear necessary for the skipping phenotype to occur, and that a poly-leucine expansion co-occurs with the protein reduction phenotype. These findings improve the understanding of which N-terminal residues contribute to ribosomal skipping, and provide an improved framework for understanding the function of 2A-like peptides in native eukaryotic protein contexts.

2A-mediated skipping could only be tested in low-throughput prior, but the TR as a tool permits high-throughput screening. While the TR functions well for its purpose, the functional design has limitations. First, fluorescence does not always scale proportionally per fluorophore*^37^*, so the relationship between protein quantity and fluorescence intensity would need to be calibrated for exact quantitation. Second, housing 2As in the flexible loop of mKG produces a fragmented and potentially unstable beta-barrel attached to mCherry, as realized by the co-ordinate decrease in red fluorescence observed with skipping sequences such as P2A. Third, a subset of viral 2A peptides mediate skipping through a frameshifting mechanism*^38^*, which was not tested for here but could be by frameshifting miRFP670 in the TR. Fourth, the engineered loop of mKG, and our data resulting from a 20-residue insertion into it, will likely not completely represent the properties of any given 2A-like peptide in its native context. Despite these limitations, the TR permits bulk canvassing of 2A-like peptides for activity.

We originally expected to observe a mixture of the skipping, nonskipping, and falloff outcomes based on the known functionality of the ribosomal skipping mechanism*^2,11^*. The observation of reduced protein abundance paired with each of the other skipping outcomes was unexpected based on prior reports of 2A peptides. It is unclear whether reductions in protein abundance are rare with viral 2A-like peptides, or whether they exist but were not captured in the in vitro translation systems commonly used, which likely differ in their rates of ribosomal loading and proteasomal activities.

For the structures we predicted and annotated, the major groups we identified were LINE1 non-LTR ORF1p and ORF2p, death-domain containing proteins, ankyrin-like proteins, and RapGEF. While we have not verified the function of the 2A-like peptides in their native contexts, we hypothesize that protein expression regulation via reductions in protein abundance is a central phenotype of many such 2A-like peptides. Among 2A-like peptides from death-domain containing proteins, 73% were in a reduction class. Such proteins regulate cell death, and overexpression can lead to cell death^39–41^, evolution of a peptide motif that mediates protein reduction could contribute to cell homeostasis. Meanwhile, 83% of 2A-like peptides from RapGEF were in a reduction class. Notably, RapGEF is involved in neural progenitor development^42^, which requires tight temporal regulation of protein expression. Some neurodevelopmental proteins are regulated through the induction and resolving of stalled ribosomes^43–45^. Consequently, regulation of RapGEF expression via a protein reduction peptide motif is plausible.

50% of 2A-like peptides from Ankyrin-like proteins were in a reduction class, and it is less clear how these proteins might utilize reduction. Among other non-retrotransposon annotations, observation of skipping or reduction peptides was rare, and there were no annotations with strong signal from skipping class peptides. Functional 2A-like peptides appear to be non-randomly harnessed by evolution and appear to enrich for reduction specifically, indicating that this may be the primary phenotype, and that skipping is only harnessed when reduction is not the main beneficial outcome.

Our observation of ORF1p and ORF2p mirrors prior reports*^4,14^*, though we observed instances in a wider range of clades, mostly due to the expanded number of sequenced genomes in the past decade. ORF1p is generally thought to be regulated by host cells through mechanisms such as splicing*^46,47^*. ORF1p and ORF2p can be observed together in some retrotransposon lineages, and the expression of ORF2p is controlled alongside ORF1p in such cases. Orphan ORF2p are also regularly observed, though the expression regulation mechanism is unclear*^48^* and retrotransposition function is still dependent on ORF1p*^49^*. 2A-like peptides were regularly observed in the N-termini of both ORF1p and ORF2p across multiple retrotransposon clades. If reduction is indeed a central phenotype of eukaryotic 2A-like peptides, LINE1 elements may use them to aid in evasion of the innate immune response*^50^* until certain intracellular signals or conditions are encountered.

Skipping and reduction class 2A-like peptides shared significant sequence-position commonalities. In analyzing both groups, we observed a regular pattern of N-terminal leucine and hydrophobic residues paired with the expected C-terminal canonical and non-canonical core motifs. Relatedly, we observed an enrichment of alpha-helical secondary structure from position −19 to −10 specific to the skipping and reduction classes. 2A alpha-helical formation has been hypothesized to occur in the ribosomal exit tunnel*^2,15,19^*. The specific positioning of repeat leucines and the regular terminated position of the alpha-helix immediately following the leucine residues are likely mechanistically important.

Recent work indicates that alpha-helix formation in the ribosomal tunnel is broadly possible*^51^*, and facilitates interactions of the nascent peptide with the ribosomal exit tunnel which occasionally cause stalling*^52–55^*, though it remains difficult to predict*^56^*. How many types of ribosomal stalling-mediating alpha-helices (‘stall-helices’) exist in proteomes remains unclear. From our results, 2A-like peptides consist of a 2A-like C-terminus and a leucine-rich N-terminal alpha-helix. Prior reports indicate that 2A-like peptides can induce ribosomal stalling*^38,55^*, which could result in protein reduction if ribosomal escape is not achieved^55,57^. Protein reduction from stalling at an engineered stop codon at the C-terminus of a viral 2A peptide was overcome by co-expression of the viral cytosolic OTU DUB from Congo hemorrhagic fever virus^55^, potentially related to the Josephin and OTU DUBs observed associated with ORF2p. Consequently, our results indicate that 2A-like peptides function as leucine-centric stall-helices, where the correct C-terminal residues induce a ribosomal skip to mediate ribosomal release from the stalled state.

The stall-helix model of 2A ribosomal skipping functionality likely works through the steric effects of leucine side chains interacting with the ribosomal exit tunnel (Figure 10). Though we have not determined what atomic interactions would mediate stalling, this model incidentally explains the differential functionality in bacteria and eukaryotes. Bacterial ribosomal exit tunnels are longer than in eukaryotes, with larger cross-sectional areas further from the peptidyl transferase center (PTC)*^31,58^*. While both bacteria and eukaryotes have constriction sites 30Å from the PTC, eukaryotes also have a second constriction site that is 50Å from the PTC*^58^*.

**Figure 10.**
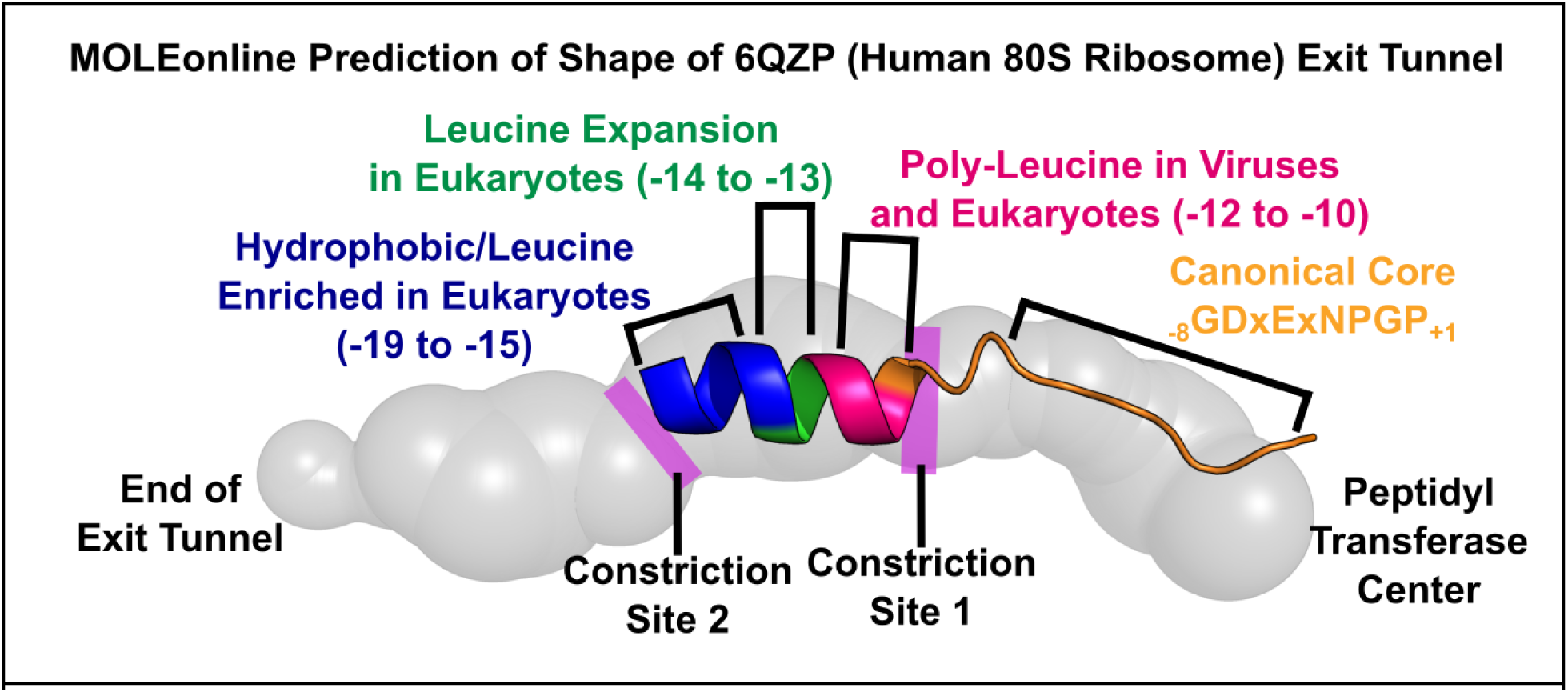
The N-Terminal Alpha Helix of 2A-like Peptides Fits Between the Constriction Sites of the Eukaryotic Ribosomal Exit Tunnel. The alpha helix in 2A-like peptides fits between the two constriction sites of eukaryotic ribosomes, and the 10 C-terminal residues in free-loop conformation extend to the peptidyl transferase center. Exit tunnel geometry of 6qzp was estimated using MOLEonline^59^. Blue (position −19 to −15) represents the region enriched in hydrophobic and leucine residues in eukaryotes. Green (position −14 to −13) represents the position most enriched for leucines in eukaryotic 2A-like peptides but not viral 2A peptides. Pink (position −12 to −10) represents the position enriched for leucines in both eukaryotic 2A-like peptides and viral 2A peptides. Orange represents the location of the canonical core C-terminal motif and the residue at position −9 which bridges the alpha helix and the canonical core C-terminal motif.

Assuming a fully extended conformation, the 9 residues preceding the skipped bond would have an approximate length of 34.2Å, just following the first constriction site in both taxonomic domains. The N-terminal 10 residues in an alpha helical formation would constitute 2.8 turns of the alpha-helix spanning 15Å, for a total distance of 49.2Å from the PTC, corresponding to the site of the second constriction site in eukaryotes (Figure 10). Residues on the alpha helix would be geometrically positioned to interact with the exit tunnel. The enriched leucine residues from position −14 to −11 are likely the major interacting residues, though cumulative interactions along the alpha-helix likely still cumulatively impact skipping.

The stall-helix model incidentally provides a possible explanation for how the glycine-serine GSG N-terminal linker improves skipping function*^10,60^*. 2A peptides are considered to be 18-21 residues, though some can retain function at 16 residues*^15^*. A GSG linker introduces highly flexible, helix-interrupting residues*^61^* at the N-terminus of the 2A at the second constriction site, ensuring that the alpha-helices of engineered viral 2A peptides terminate between the constriction sites. It is also notable that viral 2A peptides and active eukaryotic 2A-like peptides frequently have a _-9_SG_-8_ flexible dipeptide approximately aligning with the first constriction site (Figure 8). This is also consistent with the pattern of the alpha helices of the eukaryotic protein structures we predicted starting between −25 and −19, which would be near the second constriction site (Figure 8H).

The lack of a second constriction site in bacteria would mean a wider range of sampled residue positions in the exit tunnel. As the hypothesized leucine sidechain geometries would not be enforced by the second constriction site, 2A-like skipping would necessarily be impossible in bacteria, which is consistent with existing studies. This would not preclude the possibility of a different bacteria-specific mechanism, though to our knowledge no such mechanism has been described.

This model of 2A-like peptides functioning as stall-helices helps explain how a peptide sequence imparts ribosomal skipping functionality. Peptide sequences that are alpha-helical, highly enriched in leucine, and containing a C-terminus similar to either the canonical or non-canonical core motif will function as a stall-helix, and either ribosomally skip or mediate protein reduction as the ribosomal stall is cleared through degradation. Though this work identifies sequence trends that are likely to contribute to the mechanism, the exact contributions of any given residue at any given position remain unclear. Structural modeling of eukaryotic ribosomes considering the 2A as a stall-helix will be important for determining which atomic interactions are necessary to induce the stalling phenotype. Continuing exploration of the mutational space around functionally active peptides will be necessary for identifying 2A-like peptides with improved functionality. Continued use of the TR for further high-throughput studies will be important in the process of identifying 2A-like peptides that outperform P2A.

## Materials and Methods

### Identification of Candidate 2A-like Sequences

45 functional 2A and 2A-like peptides were identified in the literature, and a 20 amino acid string for each was identified in the source paper (Table S1). Each peptide was utilized as a query sequence for a BLASTp search of the NIH NCBI database of non-redundant protein sequences. An EXPECT threshold of 200,000 and a word size of 2 to maximize search capacity. A PAM30 matrix was utilized for the search. An Existence of 9 and an Extension of 1 were used as gap costs for the search. No compositional adjustments were utilized. The top 5000 proteins for each BLAST search were downloaded as text files in FASTA format and used for downstream processing.

All resulting data was combined, and species names were extracted from the headers. Any headers containing a viral or non-species term for removal were diverted into a file as removed entries. The genus and species were then extracted for each maintained entry, as was the term Candidatus where relevant. Each associated species was checked against the NCBI database for species where genome assemblies are available (https://www.ncbi.nlm.nih.gov/genome/browse<u>&/</u>) to confirm that only one genome entry would be called to, and the coding DNA was downloaded from NCBI datasets*^22^* (https://www.ncbi.nlm.nih.gov/datasets/) via the command line tool.

We next translated the coding DNA for each species, and identified each instance of the ‘NPGP’ amino acid motif, searching for any peptide ending in xNGP, NxGP, NPxP, or NPGx, and noted the 20-mer residue sequence, the full protein, the coding sequence, and the transcript ID as a candidate entry.

To retrieve the taxonomic domain for any species containing a putative 2A-like peptide, we downloaded the taxonomy dump from NCBI on June 24, 2024. We subsequently utilized the NCBItax2lin tool (https://github.com/zyxue/ncbitax2lin)*^62^* to extract the domain for each taxonomy dump, and attached the domain to the associated 2A-like peptide (Table S3).

### Comparison to Previous Datasets

As others have recently performed large bioinformatic screens searching for 2A-like peptides using an alternative approach*^15^*, we tested how much overlap existed between our bioinformatic datasets. To maximize comparability of the data, we removed all Viral peptides from the existing dataset. Since current reporting identifies a 2A peptide as between 18 and 23 amino acids*^9,10^*, we removed all samples shorter than 18 amino acids from the existing dataset. This only resulted in 157 unique peptides. Of these, 152 had no matches within our identified dataset and 5 had 1 peptide match. No peptides with reported activity passed the threshold of 18 amino acids in length.

While the existing data considered shorter peptides, local context of the remaining 2 to 5 preceding residues likely impacts skipping, so generality of skipping functionality in these shorter peptides is unclear. Further, most data presented here is functional while most pre-existing bioinformatic data is not, making the data mostly incomparable. As such, further comparisons were not performed.

### Library Design

To prevent unintended cuts by BsaI, all bacterial, eukaryotic, and archaeal coding sequences identified were given synonymous mutations to remove BsaI recognition motifs. Eukaryotic and Archaeal coding sequences containing BsaI recognition sites were included with and without these domesticating mutations

All 2A-like peptides were checked for the _-7_DxExNPGP_+1_ conserved terminal motif. All peptides without the motif were checked for the minimum number of nucleotide substitutions to reconstruct a _-7_DxExNPGP_+1_ motif, using nucleotide frequency to determine what nucleotide to use in the case of ties. Entries requiring 3 or fewer mutations were generated and included in the library.

The nucleotide sequences for Thosea Asigna Virus 2A, Equine Rhinitis A Virus 2A, Porcine Teschovirus 2A, and Foot and Mouth Disease Virus 2A were retrieved, and a synonymous variant sublibrary was created for each, with one entry included for each possible single-residue synonymous codon variant. One additional entry was included, where every codon was simultaneously mutated to a synonymous codon.

For controls, the base coding sequences for Thosea Asigna Virus 2A, Equine Rhinitis A Virus 2A, Porcine Teschovirus 2A, and Foot and Mouth Disease Virus 2A were included, as well as 3 backup controls identified by Donnelly et al*^1^* which are similar to P2A, T2A, and E2A. We included variants of Thosea Asigna Virus 2A, Equine Rhinitis A Virus 2A, Porcine Teschovirus 2A, and Foot and Mouth Disease Virus 2A where the terminal motif coding _-3_NPGP_+1_ was mutated to instead translate for _-3_NAGA_+1_, as well as a repeated GS linker that is 20 amino acids long.

Eukaryotic, bacterial, and archaeal coding sequences were each included once in the library. The minimum mutation entries were each included once in the library. Synonymous variants of F2A, P2A, T2A, and E2A were each included twice in the library. Controls were included in a cycling pattern until 7500 members had been included.

A sequence of ATCACGCGGGGTCTCAAGCTG was appended to the beginning (5’) of each library entry, and GGCTCTGAGACCGCAGCTG was appended to the end (3’). These contain a pair of inward-facing BsaI recognition sites, allowing library entries to integrate with scar-free repair into the TR. The library was ordered from Agilent Technologies as a HiFi Oligo Library of 7500 members.

### Library and Sample Cloning

To prepare the oligo insert for insertion into the plasmid, we combined 1 μL of 10 nM oligo, 1 μL of 10 nM reverse primer (CAGCTGCGGTCTCAGAGC) for the oligo, 3 μL of nuclease free water, and 5 μL of KAPA HIFI polymerase (Fisher Scientific, 50-196-5299). PCR was performed with 5 minutes at 95°C, 10 cycles of 20 seconds at 98°C, 15 seconds at 55°C, and 30 seconds at 72°C, followed by 5 minutes at 72°C. This was supplemented with an addition of 1.25 μL each of 10 uM forward (ATCACGCGGGGTCTCAAGCT) and reverse primer for the oligo, and 2.5 μL of KAPA HIFI polymerase. PCR was performed with 5 minutes at 95°C, 20 cycles of 20 seconds at 98°C, 15°C, followed by 5 minutes at 72°C. 4 such tubes were produced in parallel.

Samples were run on a 1% agarose TE buffer gel. Bands were excised from the gel, and extracted with a Bio-Rad Freeze ‘N Squeeze DNA Gel Extraction Spin Column (Bio-Rad, cat. no. 7326165) and cleaned with a Zymo DNA Clean & Concentrator-5 kit (Zymo Research, cat. no. D4004) and pooled.

75 ng of target plasmid, 4 molar equivalents of the PCR prep, 1 μL of T4 DNA Ligase buffer (New England Biolabs, B0202S), 0.25 μL of T4 DNA Ligase (New England Biolabs, M0202M), and 0.75 μL of BsaI HiFi V2 (New England Biolabs, R3733S) were combined with nuclease free water to a final volume of 10 μL and incubated for 60 minutes at 37°C, followed by 5 minutes at 60°C following standard practice of golden gate cloning*^24^*. 4 tubes were prepared in parallel and combined for library samples, and single tubes were prepared for individual samples.

For library samples, NEB 3020 electrocompetent bacteria were prepared in house at high concentration. 25 μL of bacteria were added to a 0.1cm electroporation cuvette alongside 1μL of plasmid library prep. Bacteria were pulsed at 1.8 kV, 200 Ω, and 25 μF. Cells were resuspended in 1 mL of LB media and allowed to recover for 1 hour at 37°C. 8 such cuvettes were prepped and pooled after recovery. 10% of cells were plated on LB media plates spiked with 150 μg/mL Ampicillin, and the remaining 90% were suspended in 850 mL of LB liquid media spiked with 150 μg/mL Ampicillin, and all bacteria were grown for 18 hours. After 18 hours, plates indicated a minimum of 30,000 transformants, indicating a minimum 4-fold coverage of the expected library. The plasmid library was extracted from bacteria grown in LB liquid media using the Promega PureYield Plasmid Midiprep System (Promega, A2492), and final library concentration was confirmed via Broad Range dsDNA qubit (Thermo Fisher Scientific, Q32853).

For individual samples, chemical-competent bacteria were prepared in house. 99 μL of cells combined with 1 μL of plasmid and let rest on wet ice for 30 minutes, heat-shocked at 42°C for 1 minute, rested on wet ice for 5 minutes, and allowed to rest at 37°C for 30 minutes. Samples were plated on LB media plates spiked with 150 μg/mL Ampicillin and grown for 18 hours. Individual colonies were then picked and grown overnight in 3 mL of LB media with 150 μg/mL ampicillin, and subsequently miniprepped using either the Zymo Zyppy miniprep kit (Zymo, 11-309) or the Thermo GeneJet miniprep kit (Thermo Fisher Scientific, FERK0503). Final sample concentration was confirmed with the ThermoFisher Qubit Broad Range dsDNA quantitation kit.

### Cell Culture

Cell lines were cultured in media comprising Dulbecco’s Modified Eagle’s Medium containing L-glutamine and 4.5 g/L glucose and not containing sodium pyruvate (Corning, 10-017-CV), 10% Fetal Bovine Serum (Gibco, 10437028), 100 μg/mL penicillin and 0.1 mg/mL streptomycin (Corning, 30-002-CI), and 20 μg/mL doxycycline (Fisher Scientific, AAJ67043AD). Passaging was performed via detachment utilizing trypsin with ethylenediaminetetraacetic acid (EDTA) at 0.25% (Corning, 25-053-CI). Landing pad cells derived from G417A (G417A_pLenti-Tet-BattP(GA)-BFP-2A-iCasp9-2A-Blast-rtTA3; Addgene plasmid number 200631) as described previously*^63^* were used for all experiments. For long term passaging of cells, 20μg/mL blasticidin (InvivoGen, ANT-BL-1) was regularly pulsed on cells to ensure constitutive transcription of the Landing Pad site in untransfected cells.

### Recombination of Plasmids into Landing Pad Cells

HEK293T G417A Landing Pad cells were plated on 6 well plates at 200,000 cells per well. Cells were allowed to grow for 24 hours, and then transfected using either xFect (Takara Bio, 631318) or Fugene 6 (Promega, E2691), with 100 ng of a vector containing eeBxb1^64^ (Addgene plasmid A222339). After 24 hours, media was changed, and cells were transfected again using either the same transfection reagent as the day prior, as well as 100 ng of the vector containing eeBxb1 and 1000 ng of library-containing vector, with another media change 24 hours later. After 3 days, media was changed to media containing 10 nM AP1903 and 1 μg/mL puromycin. Cells were allowed to grow in puromycin and AP1903-containing media for two weeks, with media changes and splits as necessary. Cells were maintained, and changed to cell culture media without puromycin and AP1903 at least 3 days prior to any experiments as puromycin affects the ribosome, and may impact homeostatic levels of skipped protein products.

### Trifluorescent Reporter Construct

A plasmid construct coding for an AttB(GA) site, an HA tag-mCherry-monomeric Kusabira Green-miRFP670-DYKDDDDK tag fusion protein, and the EMCV IRES followed by a Puromycin resistance gene was developed for this work. BsaI recognition sites were added within the mutable flexible loop within mKG, oriented facing outwards. The plasmid was domesticated to remove all other BsaI recognition sites. The plasmid was deposited with Addgene as pJCS068c3 (pJCS068c3_AttB_HA-mCherry-mKG1-BsaI-mKG2-miRFP670-FLAG_IRES-PuroR) under Addgene ID 258519 A flipped version of the construct where mCherry and miRFP670 were swapped was also developed, and deposited with Addgene as pJCS108C (pJCS108C_AttB_HA-miRFP670-mKG1-BsaI-mKG2-mCherry-FLAG_IRES-PuroR) under Addgene ID 258520. Both plasmids are compatible with HEK293T G417A Landing Pad cells.

### Western Blotting

Samples were prepared by measuring concentration using a Bicinchoninic Acid (BCA) assay, then mixing 8 μg of protein in radioimmunoprecipitation assay (RIPA) buffer (Thermo Scientific, 89901) with 4X LDS loading dye (GenScript, M00676-10). Samples were then run on BioRad Mini-PROTEAN TGX Precast protein gels (cat. No. 4561096) at 40V. Samples were then transferred to PVDF and blocked with 5% powdered milk in Tris-buffered saline comprising 250 mM TRIS Base, 27 mM KCl, and 1.37 M NaCl (Dot scientific, DST60075-4000) at 1/10 dilution with 1 mL/L Tween 20 (MP Biomedicals, TWEEN201) for 1 hour. Samples were incubated with 1:1000 dilutions of anti-HA tag antibody (Cell Signaling Technologies, catalog no. 3724S) and anti-DYKDDDDK antibody (Cell Signaling Technologies, catalog no. 8146S) for 48 hours. For figure 2, Figure 4, Figure S13, and Figure S25, 1:5000 dilution of Goat-anti-Rabbit secondary conjugated to Alexafluor 647 (Invitrogen, a21246) was used as secondary for the HA antibody and 1:5000 dilution of Goat-anti-Mouse secondary conjugated to HRP (Jackson Immuno, 115-035-166) was used for the FLAG antibody, with SuperSignal West Femto Luminol (Thermo Fisher Scientific, 34096) used as the reagent for HRP.

For the reclassification blots, sample prep was identical. We first blotted with 1:1,000 dilution of mouse anti-DYKDDDDK tag antibody, followed by a 1:5,000 dilution of goat-anti-mouse secondary conjugated to Alexafluor 594 (Invitrogen, a11005) and imaged at 594nm due to the secondary being cross-reactive with rabbit antibodies. We then applied 1:1000 rabbit anti-HA tag antibody, followed by 1:5000 dilution of goat-anti-rabbit secondary conjugated to Alexafluor 647 and imaged at 647nm. We then blotted with anti-beta-actin conjugated to HRP (Thermo Fisher, HRP-60008), which was mildly cross reactive with the mouse anti-DYKDDDDK tag antibody. Relative differences in actin signal correlated with relative levels of FLAG signal that were observed in those samples, in addition to the expected actin band.

### Microscopy

Microscope images were taken with a Nikon Eclipse Ti2-e fluorescent microscope at 40x magnification, outfitted with a SOLA SM II 365 light engine (Lumencor), a CFI Plan Apochromat DM Lambda 20X objective or a NIKON Plan Fluor 4X objective, GFP (#96392), Texas Red (#96395), or Cy5 (#96396) filter sets, and imaged with a DS-QI2 monochrome CMOS camera. For each sample, mTagBFP2 was imaged for 744ms at 11.4x gain, mCherry was imaged for 605ms at 4.1x gain, mKG was imaged for 857ms at 4.1x gain, and miRFP670 was imaged at for 605ms at 9.3x gain. Images were pseudocolored and merged using Fiji imageJ version 2.16.0/1.54p.

### Flow Cytometry Fluorescence Measurement

Samples were detached with trypsin-EDTA and resuspended in Fluorescence Activated Cell Sorting (FACS) media comprising Phosphate Buffered Saline (Fisher Scientific, MT21031CV), 5% fetal bovine serum, and 1mM EDTA (Thermo Fisher Scientific, 15575020). Samples were processed on a BD LSRFortessa Cell Analyzer. For fluorophores, mTagBFP2 was excited with a 405 nm laser and emitted light was collected with a V450/50 nm filter, mCherry was excited with a 532 nm laser and emitted light was collected with a G610/20 nm filter, mKG was excited with a 488 nm laser and emitted light was collected with a B515/20 nm filter, and miRFP670 was excited with a 640 nm laser and emitted light was collected with a R670/30 nm filter. FCS files were collected, and data was analyzed using BD FlowJo V10.10.0. Samples were processed to isolate live cells and singlet cells, and data was exported into CSV format for subsequent processing.

### Plate Reader Bulk Fluorescence Measurement

Samples were detached with trypsin-EDTA and resuspended in FACS media and loaded as a 100 μL volume within a 96-well clear bottom plate. A cell concentration curve comprising wells with 100,000 to 1,000,000 cells was prepared in parallel in 100 μL of FACS media and loaded on the 96-well plate. The plate was read out on a BioTek Synergy H1 microplate reader using absorbance at 520 nm, 620 nm, and 720 nm. Fluorescence was measured for mTagBFP2 with an excitation of 399 nm and an emission of 454 nm, for mCherry with an excitation of 587 nm and an emission of 610 nm, for mKG with an excitation of 494 nm and an emission of 507 nm, and for miRFP670 with an excitation of 642 nm and an emission of 670 nm. The absorbance values for the concentration curve were used to generate a linear equation relating cell count to absorbance, and cell count for each sample was estimated using each absorbance measurement. Fluorescence per cell was then calculated for each sample for each fluorophor

### Sort-Sequencing of Transfected Library

60 minutes prior to sorting, cells were washed with PBS, incubated with trypsin-EDTA for 2 minutes, resuspended with 4 volumes of cell culture media, and counted on a BioRad TC-20 cell counter. Cells were then spun down at 300xg for 3 minutes, and resuspended in FACS media to a concentration between 5 and 10 million cells/mL, with a minimum volume of 3 mL, which was split into 3 12×75 mm tubes (Falcon, 352052) and strained using a strainer lid to further disassociate cells.

For each run, cells were sorted once on each fluorophore using a BD FACS Symphony S6. For fluorophores, mTagBFP2 was excited with a 405 nm laser and emitted light was collected with a V431/28 nm filter, mCherry was excited with a 561 nm laser and emitted light was collected with a YG610/20 nm filter, mKG was excited with a 488 nm laser and emitted light was collected with a B515/20 nm filter, and miRFP670 was excited with a 561 nm laser emitted light was collected with a YG670/30 nm filter. Live singlet cells were isolated, and a single fluorescent color was chosen as the sort condition. Gates were set to sort 25% of cells into each tube. Fluorescent measures were taken for the first 100,000 events for each color, and sorting was continued until each collection tube had collected a minimum of 500,000 cells. This was repeated for each of mKG, mCherry, and miRFP670.

DNA was extracted from the cells using a Qiagen DNeasy Blood and Tissue Kit (Qiagen, 69506), and DNA concentrations were then determined via broad-range dsDNA Qubit. The region of interest was then amplified from genomic DNA as previously described*^29^*. Each individual sort group was sequenced using a unique barcode pair using the NextSeq 550 Illumina platform. 8 sorts were performed and 10 sequencing runs were performed. Data for 5 sorts and 7 sequencing runs was kept for mCherry and mKG, and data for 4 sorts and 6 sequencing runs was kept for miRFP670.

### Analysis of Illumina Reads

Illumina read 1 and read 2 FastQ files were recovered for 120 total samples. We generated paired reads for each sample using PEAR*^65^*, isolated reads containing a library member, and combined samples containing the same library member. We then calculated a weighted average score for each library member in the sample, using the equation:

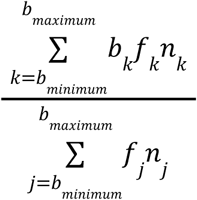

where b is the designated bin value (1, 2, 3, or 4, ordered as lowest to highest fluorescence), f is the frequency of reads in bin b, and n is the number of cells that were sorted in bin b. Scores for each library member were then averaged across sorts of the library to control for inter-sort variability.

### Classification of Skipping Sequences

To determine which samples skipped, the controls P2A, T2A, E2A, F2A, P2A_-3_NAGA_+1_, T2A_-3_NAGA_+1_, E2A_-3_NAGA_+1_, F2A_-3_NAGA_+1_, and (GS)x10 were used to train machine learning models. We utilized the Fuzzy Clustering (FC), Support Vector Machine (SVM), and Logistic Regression models of scikit-learn for all machine learning models. Models for Fuzzy Clustering, and SVM with a linear, polynomial, and radial basis function kernel were trained with mCherry, mKG, and miRFP670 Sort-Seq scores as inputs and skipping as a binary output. We also calculated the kernel density estimate in 3-dimensional space for all samples, then trained an additional logistic model using the mCherry, mKG, and miRFP670 Sort-Seq scores as well as the kernel density estimate value as inputs and skipping as a binary output. The FC model was used as the final classifier model due to the probability of class inclusion provided by the model, as well as properly sorting all controls when classes of all samples were predicted.

For reclassification, we utilized the original control panel as well as all individually tested samples from FIgure 7 to train a series of SVM models to to predict class based on Sort-Seq scores. Each input peptide was classified as either skipping, non-skipping, falloff, skipping with protein reduction, non-skipping with protein reduction, or falloff with protein reduction, with protein reduction defined as either less than 80% of the mCherry signal of P2A or less than 80% of the miRFP670 signal of P2A, though all samples in the mCherry reduction group were also in the miRFP670 reduction group. The SVM model with the polynomial kernel, followed by the SVM model with the radial basis function kernel, performed best at reclassifying the training samples into the 6 classes. Consequently, all samples measured by Sort-Seq were reclassified into the 6 classes by the SVM model with the polynomial kernel (Table S6).

### Synonymous Variant Analysis

After Sort-Seq scores were calculated for each library member, synonymous variants of P2A, T2A, E2A, and F2A were isolated. For each peptide, the arithmetic mean scores of all coding variants for mCherry, mKG, and miRFP670 were calculated from the Sort-Seq scores. The standard error of the mean and the standard deviation for the distribution of all coding variants for each peptide were also calculated based on the mCherry, mKG, and miRFP670 Sort-Seq scores. The Z-score (or number of standard deviations) of distance from the centroid was then calculated for each synonymous variant, the parent, and the _-3_NAGA_+1_ control. The 3-dimensional Z-score was then calculated as 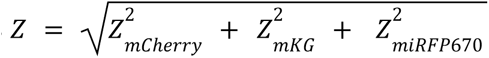. The 3-dimensional Z-score only indicates magnitude of deviation, and not direction relative to the centroid.

### Structural Prediction and Analysis

Many proteins in the bioinformatic search returned as hypothetical or unknown proteins. We identified samples that were represented at least 14 times across the mCherry, mKG, and miRFP670 Sort-Seq data, representing either one coding sequence that was well represented in all sorts, or peptides with alternative coding sequences that were reasonably represented broadly. We randomly sampled up to 20 Eukaryotic and 10 non-Eukaryotic peptide sequences for each of the 6 skipping classes. We identified the full-proteins associated with the 2A-like sequences, collapsed down to only one isoform per protein, and removed full-protein sequences over 5,000 residues in length, leaving 344 sequences for structural predictions. Full protein sequences were submitted as jobs on Google’s AlphaFold Server using AlphaFold 3*^27^* between February 21, 2026 and March 26, 2026.

We utilized PyDSSP (https://github.com/ShintaroMinami/PyDSSP)*^32,66^*to determine the secondary structure for model 0 of each AlphaFold structural prediction. We submitted model 0 for each structural prediction to Foldseek^35^ to identify proteins with similar structures using the Foldseek API (see Table S7 for search tickets). We took the top 10 results from the AlphaFold Database (AFDB) clustered to 50% sequence identity (AFDB50), AFDB for the manually curated UniProt dataset (AFDB swissprot), AFDB for the proteomes of key organisms to research and public health (AFDB proteome), and of the Protein Data Bank clustered to 100% sequence identity (PDB100) by E-value. We counted word occurrences in the top 10 results and identified the top matching structure by maximum average word usage. Authors independently curated structure match calls for each protein, and calls were consolidated to determine the likely structure and purpose of each protein. Structures were visualized using PyMOL version 3.1.6.1.

## Supporting information

Supplemental Figures 1 through 44

## Data Analysis and Data Availability

Scripts used to perform data analysis are available at https://github.com/MatreyekLab/Eukaryotic-2A-Peptides. A freeze of the data analysis and associated data files will be available at Zenodo upon acceptance for publication.

Both the TR and the flipped TR will be available as plasmids from Addgene. The TR plasmid is available as Addgene plasmid number 258519. The flipped TR plasmid is available as Addgene plasmid number 258520.

Raw and processed high-throughput sequencing data from the Sort-Seq experiments is available from the Gene Expression Omnibus database at GEO accession number GSE334901.

## Acknowledgements

We would like to thank the team at the Genomics Core Facility at the CWRU School of Medicine Genetics and Genome Sciences Department for their assistance in this work.

We would like to thank the Case Comprehensive Cancer Center Cytometry and Microscopy Shared Resource and NIH grant 1-S10-OD-026841-01A1 for their assistance in this work.

## Funding

Funder

National Institutes of Health

Grant reference Number

GM142886

Author

Kenneth A Matreyek

The funders played no part in study design, data collection, formal analysis, interpretation of results, or in the decision to submit this work for publication.

## Author Contributions

John Snell: Methodology, Data Curation, Investigation, Formal Analysis, Writing - Original Draft, Review, and Editing.

Kenneth Matreyek: Conceptualization, Methodology, Data Curation, Writing - Review and Editing, Funding Acquisition, Project Administration.

