## Supplemental Figures 1 through 44 for "High Throughput Characterization of Eukaryotic 2A-Like Peptides Identifies Novel Leucine-Associated Reduction in Protein Abundance"

**Table of Contents**

| Supplemental Figure | Description |
|---|---|
| 1 | Diagram of Landing Pad, Golden Gate Cloning Schema, and Fluorescent Outcomes in Library Context. |
| 2 | AlphaFold 3 Structural predictions of mKG, of mKG with the Golden Gate cassette, and of mKG with the Golden Gate cassette with P2A inserted. |
| 3 | Microscope images of Landing Pad cells without an inserted vector. |
| 4 | Microscope images of Landing Pad cells containing TR with P2A. |
| 5 | Microscope images of Landing Pad cells containing TR with T2A. |
| 6 | Microscope images of Landing Pad cells containing TR with E2A. |
| 7 | Microscope images of Landing Pad cells containing TR with F2A. |
| 8 | Microscope images of Landing Pad cells containing TR with P2A $_{-3}$NAGA$_{+1}$. |
| 9 | Microscope images of Landing Pad cells containing TR with T2A $_{-3}$NAGA$_{+1}$. |
| 10 | Microscope images of Landing Pad cells containing TR with E2A $_{-3}$NAGA$_{+1}$. |
| 11 | Microscope images of Landing Pad cells containing TR with F2A $_{-3}$NAGA$_{+1}$. |
| 12 | Microscope images of Landing Pad cells containing TR with (GS)x10. |
| 13 | Microscope images of Landing Pad cells containing TR with no insert. |
| 14 | Western blots testing skipping for independently transfected populations of each control vector as tested in figure 3. |
| 15 | Flow cytometry distributions for independently transfected populations for each control vector as tested in figure 3. |
| 16 | Bulk fluorescent processing of samples tested in figure 3 and principal |

| | |
|---|---|
| | component analysis testing for fluorescent similarity among independently transfected populations. |
| 17 | Flow cytometry distributions for each control insert in a flipped (miRFP670-mKG-mCherry) version of the TR. |
| 18 | Bulk fluorescent processing of controls in flipped TR, and principal component analysis testing for fluorescent similarity among constructs. |
| 19 | Sort-Seq data separated by peptide domain of origin. |
| 20 | Model classification overview for figure 4E. |
| 21 | Positional distance from centroids among single-codon synonymous variants of P2A, T2A, E2A, and F2A. |
| 22 | Microscope images of Landing Pad cells containing TR with fully recoded P2A. |
| 23 | Microscope images of Landing Pad cells containing TR with fully recoded T2A. |
| 24 | Microscope images of Landing Pad cells containing TR with fully recoded E2A. |
| 25 | Microscope images of Landing Pad cells containing TR with fully recoded F2A. |
| 26 | Western blots testing skipping for independently transfected populations of each vector as tested in figure 5. |
| 27 | Flow cytometry distributions for independently transfected populations for each vector as tested in figure 5. |
| 28 | Bulk fluorescent processing of each vector tested in figure 5, and principal component analysis testing for fluorescent similarity among constructs. |
| 29 | Testing of P2A coding sequence reported to be incapable of skipping. |
| 30 | Microscope images of Landing Pad cells containing TR with samples from figure 7, part 1. |
| 31 | Microscope images of Landing Pad cells containing TR with samples from figure 7, part 2. |
| 32 | Microscope images of Landing Pad cells containing TR with samples from figure 7, part 3. |
| 33 | Microscope images of Landing Pad cells containing TR with samples from figure 7, part 4. |
| 34 | Western blots testing skipping for each vector as tested in figure 7. |

| | |
|---|---|
| 35 | mCherry flow cytometry distributions for each vector as tested in figure 7. |
| 36 | mKG flow cytometry distributions for each vector as tested in figure 7. |
| 37 | miRFP670 flow cytometry distributions for each vector as tested in figure 7. |
| 38 | Bulk fluorescent processing of each vector as tested in figure 7, and relationship between bulk and individual fluorimetry data for sample population. |
| 39 | Residue-usage heatmaps for each reclassified class among eukaryotic-origin 2A-like peptides. |
| 40 | Residue-usage heatmaps for each reclassified class among bacterial-origin 2A-like peptides. |
| 41 | Biochemical properties by-position for each reclassified class. |
| 42 | Residue usage count in the N-terminal half of eukaryotic 2A-like peptides by residue type. |
| 43 | Secondary structures from position -19 to +1 for bacterial-origin 2A-like peptides and for all 2A-like peptides based on AlphaFold 3 structural predictions. |
| 44 | Full heatmap from figure 9A, including annotations with fewer than 3 observations. |

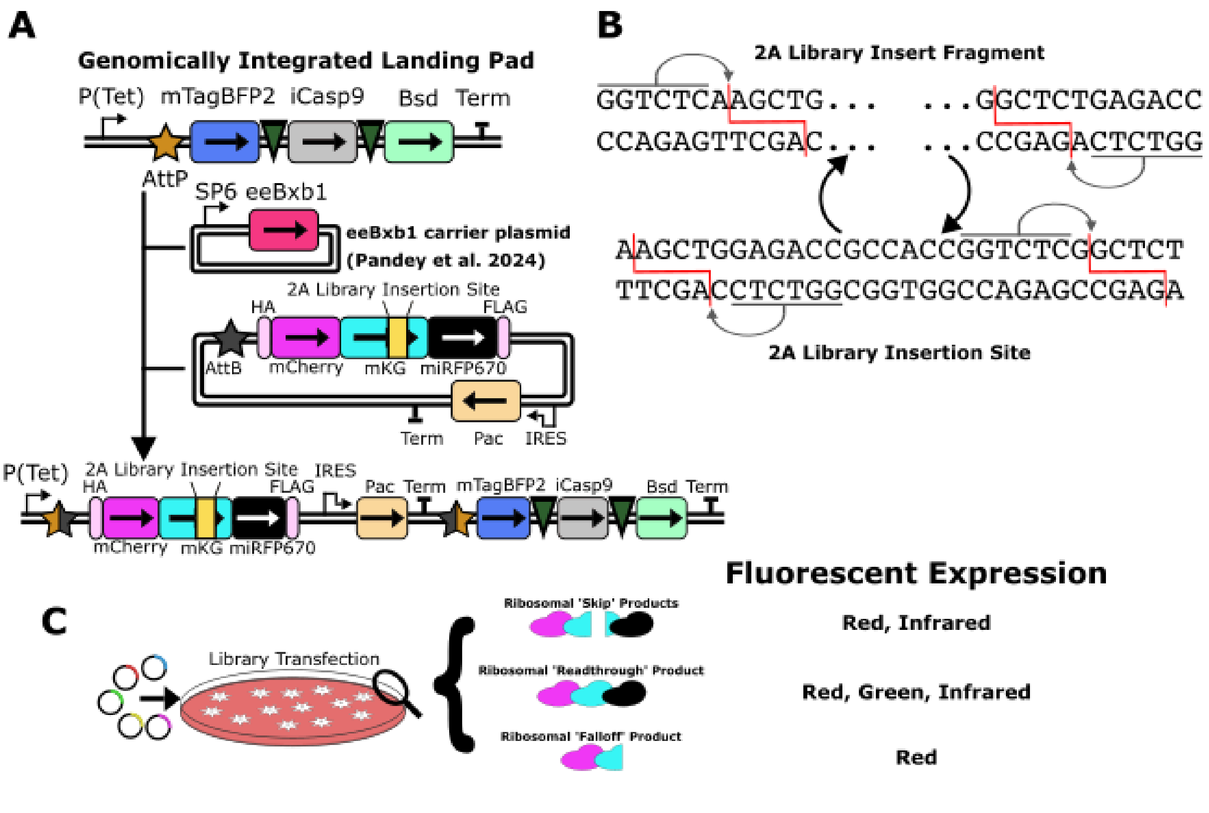


Supplemental Figure 1.

(A) Comprehensive Diagram of 'Landing Pad' genomic integration cassette (top), Trifluorescent Reporter (middle), and integrated reporter (bottom). (B) Local sequence of trifluorescent reporter (bottom) and library oligonucleotides (top) used for library insertion into vector through Golden Gate Cloning. (C) Expected behavior of peptides in library based on design. A skipped translation is expected to produce a red fluorescent protein and an infrared fluorescent protein. A readthrough translation is expected to produce a red, green, and infrared fluorescent protein. A falloff translation is expected to produce an infrared fluorescent protein. Our results indicate that the red-only fluorescent protein is non-fluorescent, but is present.

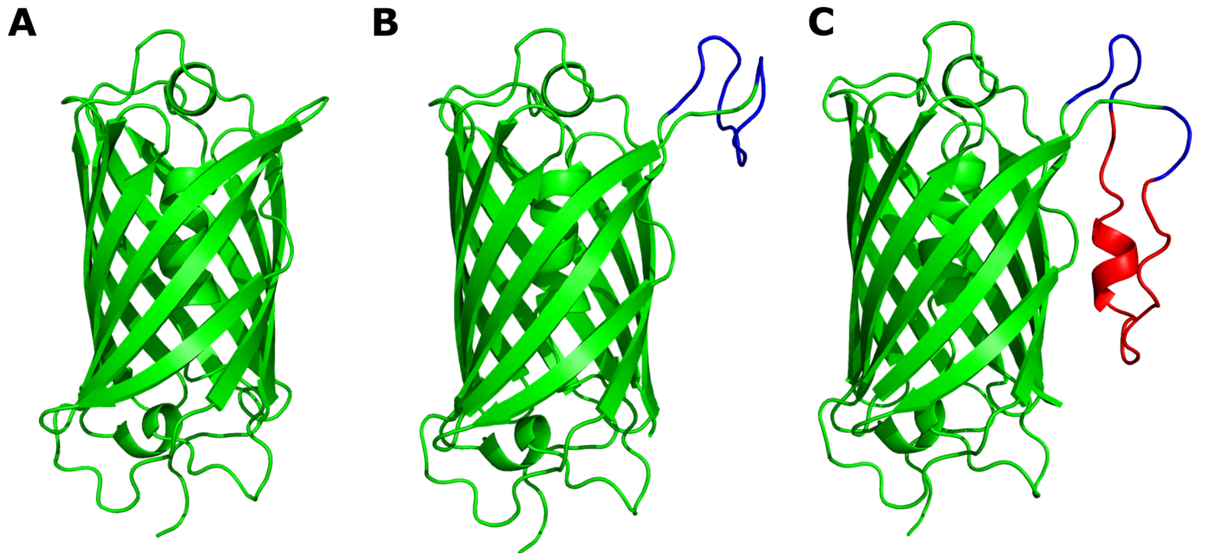


Supplemental Figure 2.

Structural predictions of monomeric Kusabira Green (mKG) using Alphafold 3 show that the Golden Gate recombination cassette does not interfere with beta-barrel structure. (A) mKG baseline structure. (B) mKG with Golden Gate recombination cassette (blue) inserted into mutable flexible loop. (C) mKG with P2A (red) recombined into Golden Gate recombination cassette in mutable flexible loop.

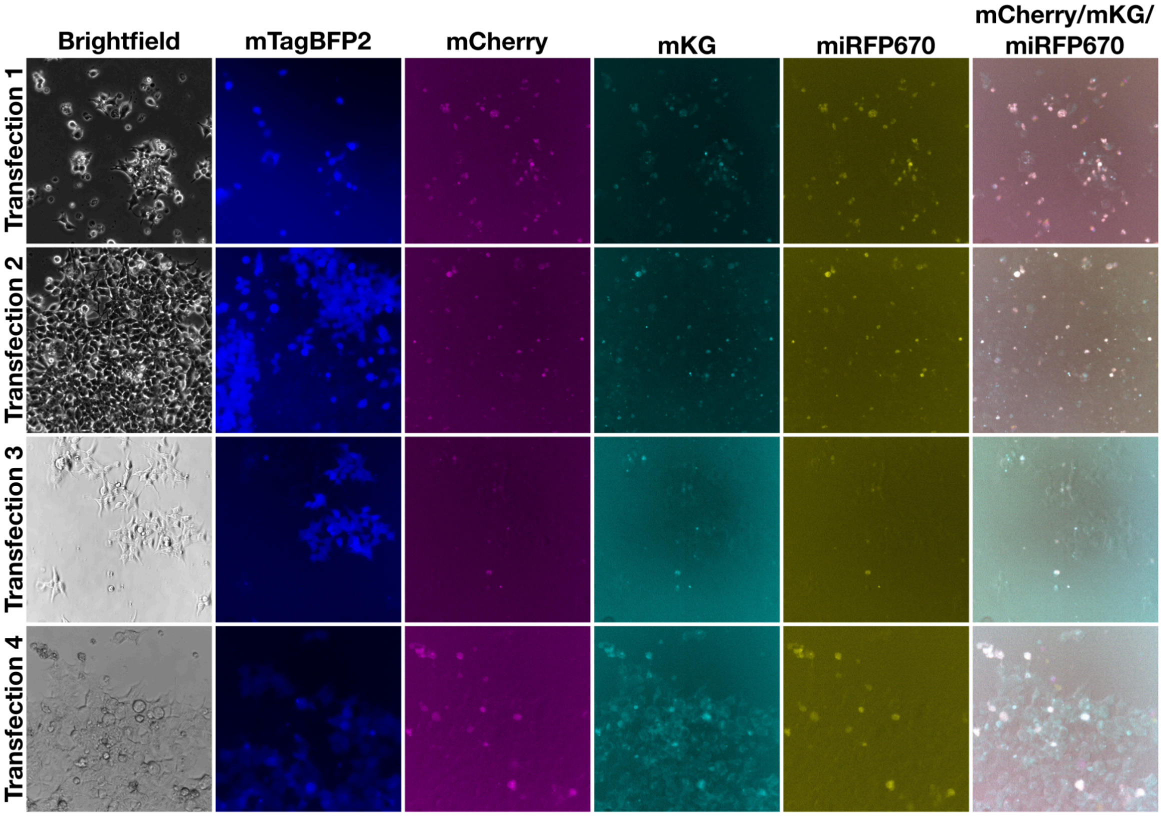


Supplemental Figure 3.

Microscope images of cells without transfected and integrated plasmid.

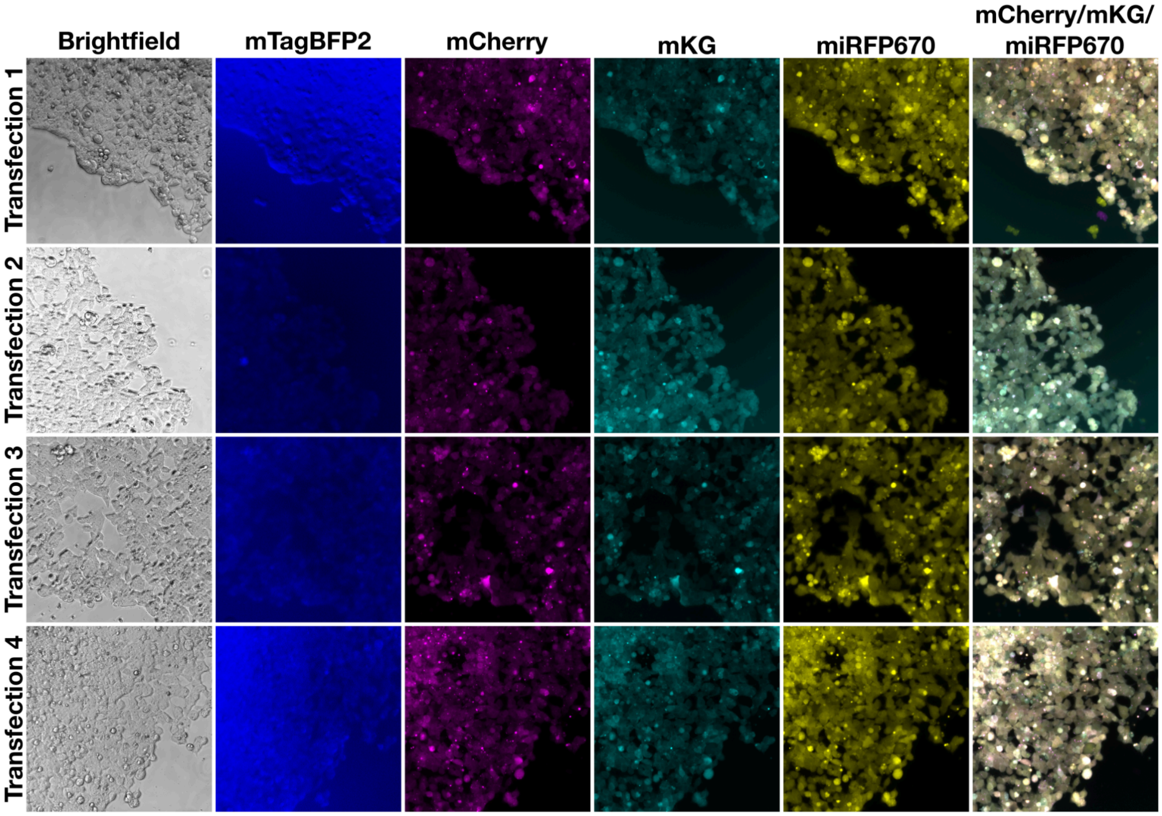


Supplemental Figure 4.

Microscope images of cells with integrated TR containing GATNFSLLKQAGDVEENPGP (P2A) inserted at the 2A insertion site.

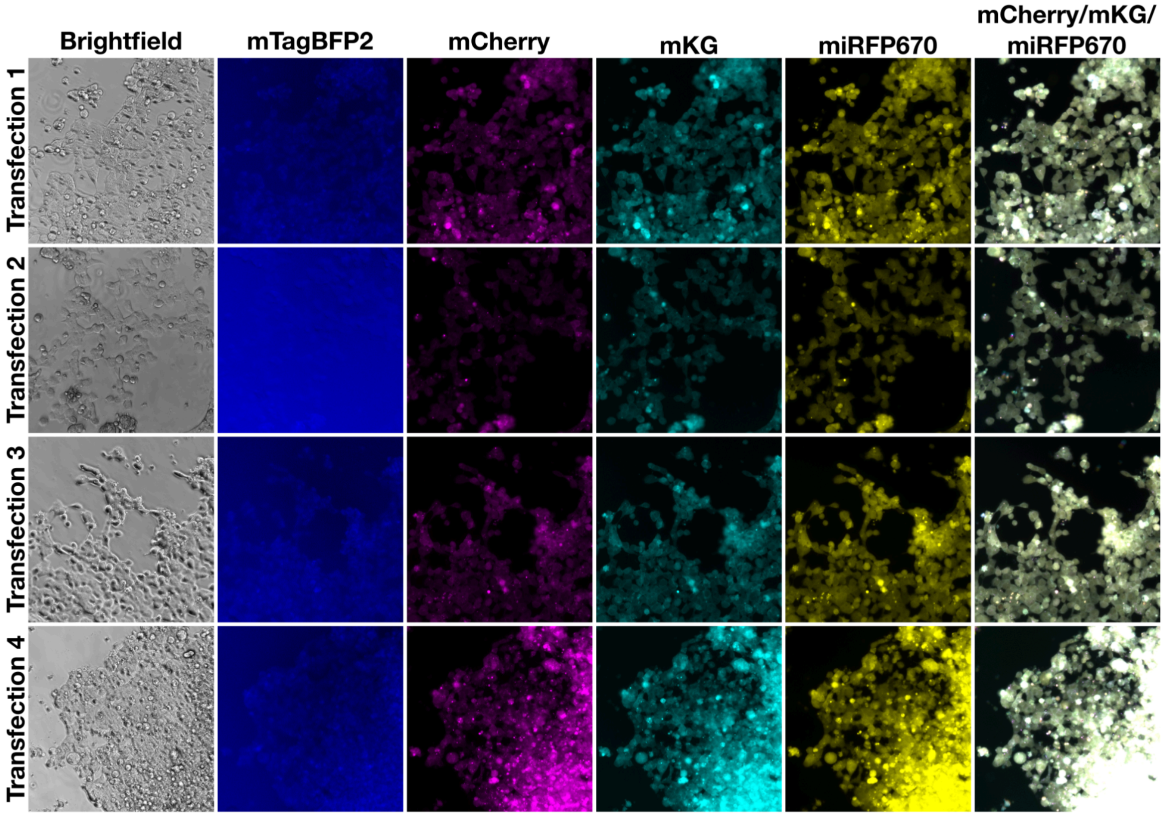


Supplemental Figure 5.

Microscope images of cells with integrated TR containing RAEGRGSLLTCGDVEENPGP (T2A) inserted at the 2A insertion site.

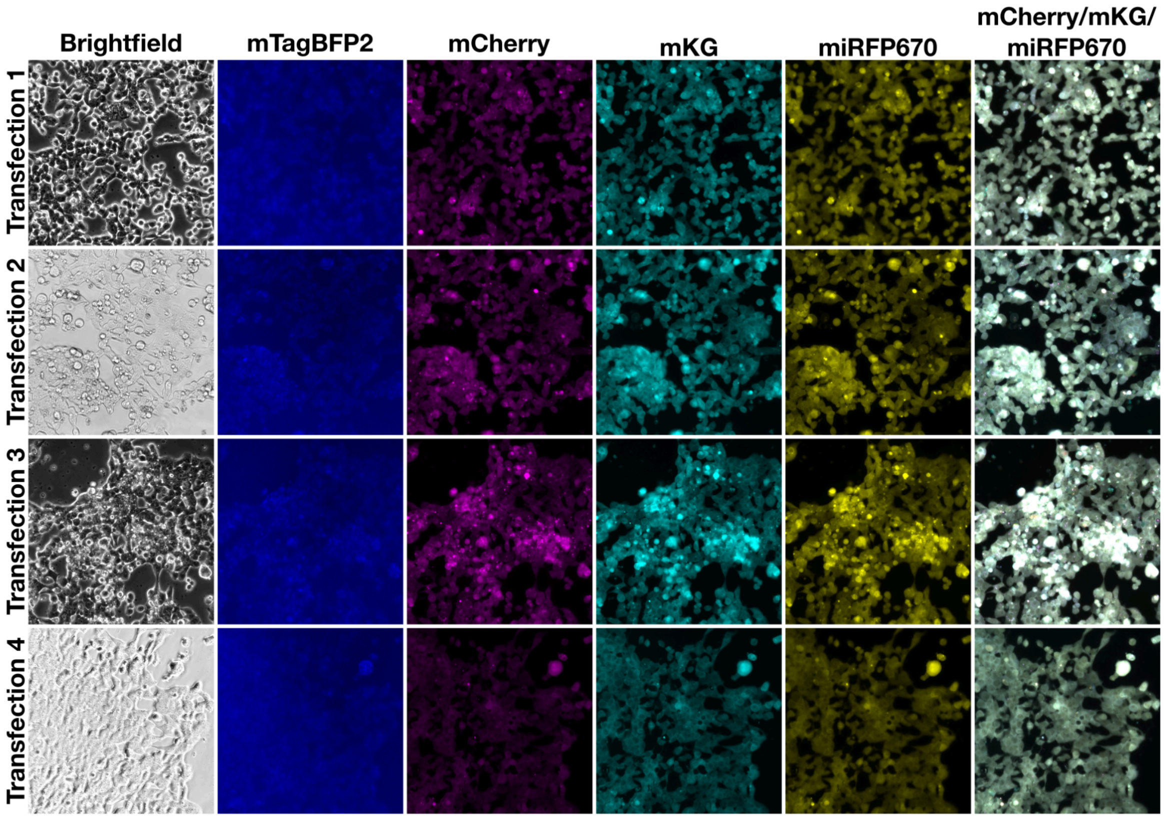


Supplemental Figure 6.

Microscope images of cells with integrated TR containing QCTNYALLKLAGDVESNPGP (E2A) inserted at the 2A insertion site.

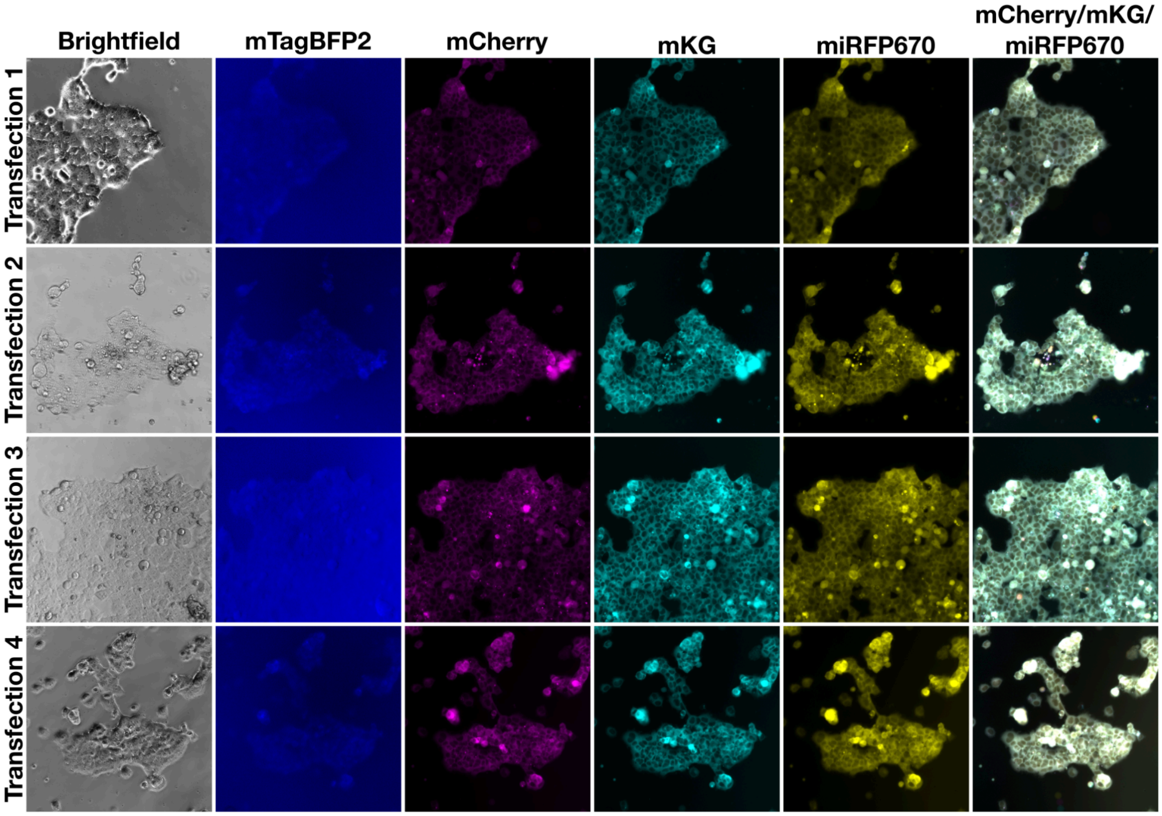


Supplemental Figure 7.

Microscope images of cells with integrated TR containing QTLNFDLLKLAGDVESNPGP (F2A) inserted at the 2A insertion site.

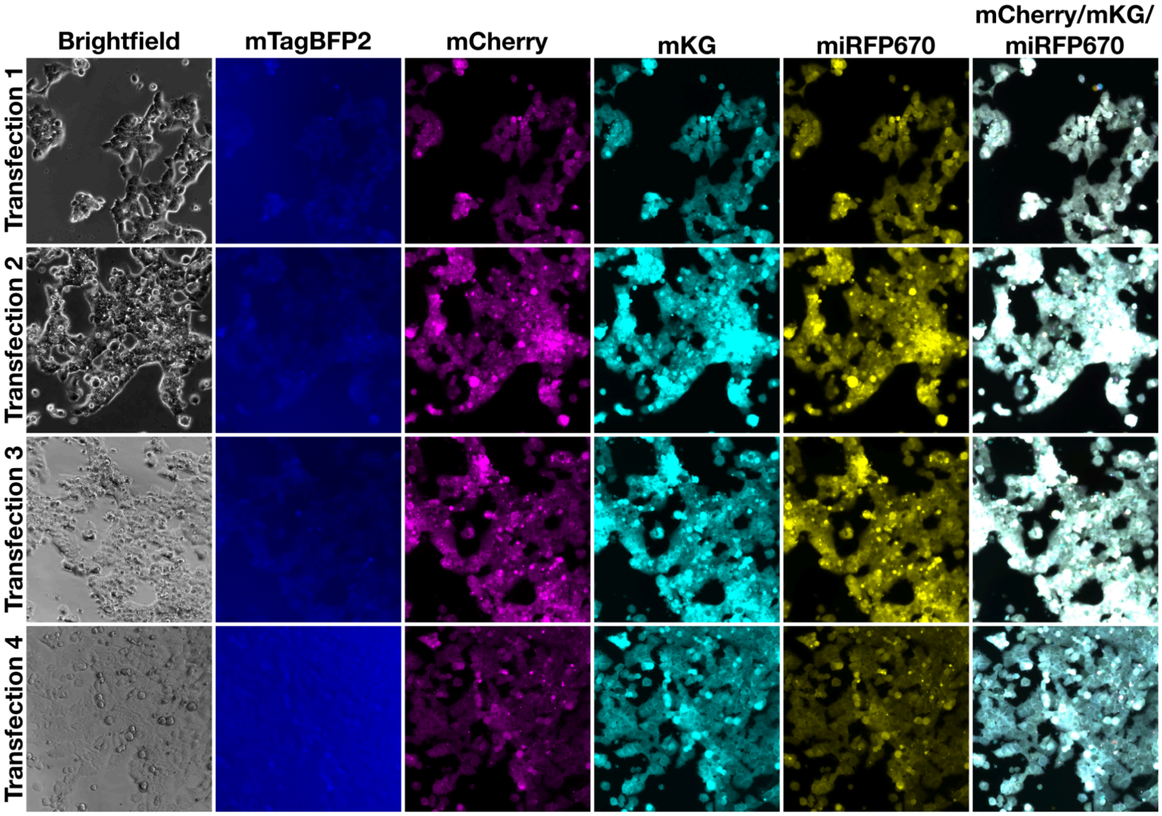


Supplemental Figure 8.

Microscope images of cells with integrated TR containing GATNFSLLKQAGDVEENAGA ($P2A_{-4}NAGA_{+1}$) inserted at the 2A insertion site.

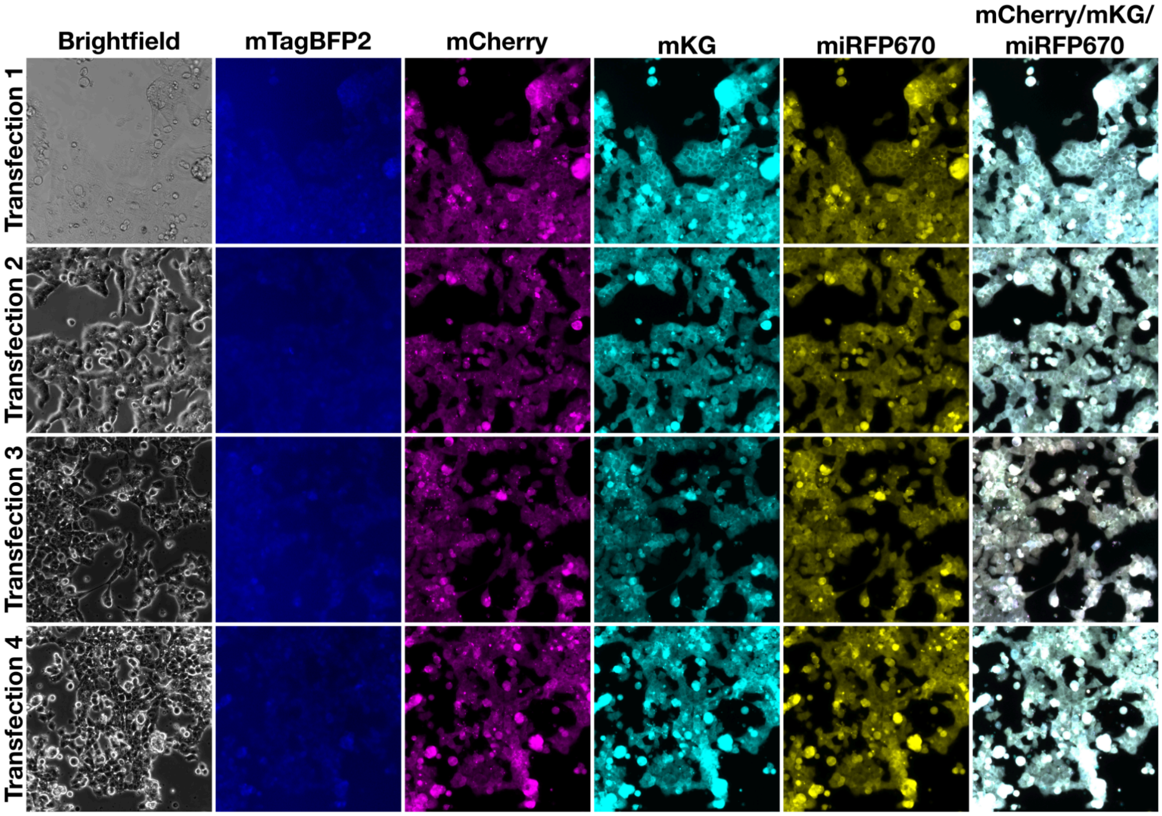


Supplemental Figure 9.

Microscope images of cells with integrated TR containing RAEGRGSLLTCGDVEENAGA (T2A$_{-4}$NAGA$_{+1}$) inserted at the 2A insertion site.

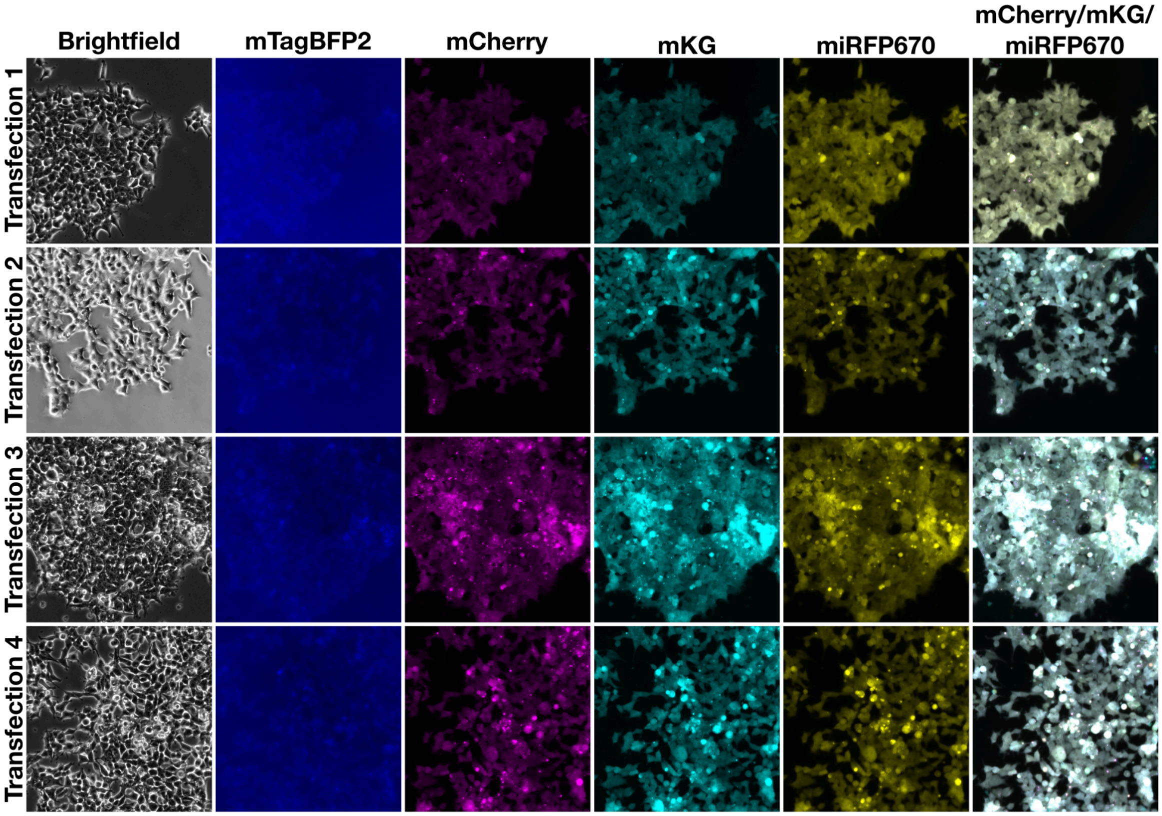


Supplemental Figure 10.

Microscope images of cells with integrated TR containing QCTNYALLKLAGDVESNAGA ($E2A_{-4}NAGA_{+1}$) inserted at the 2A insertion site.

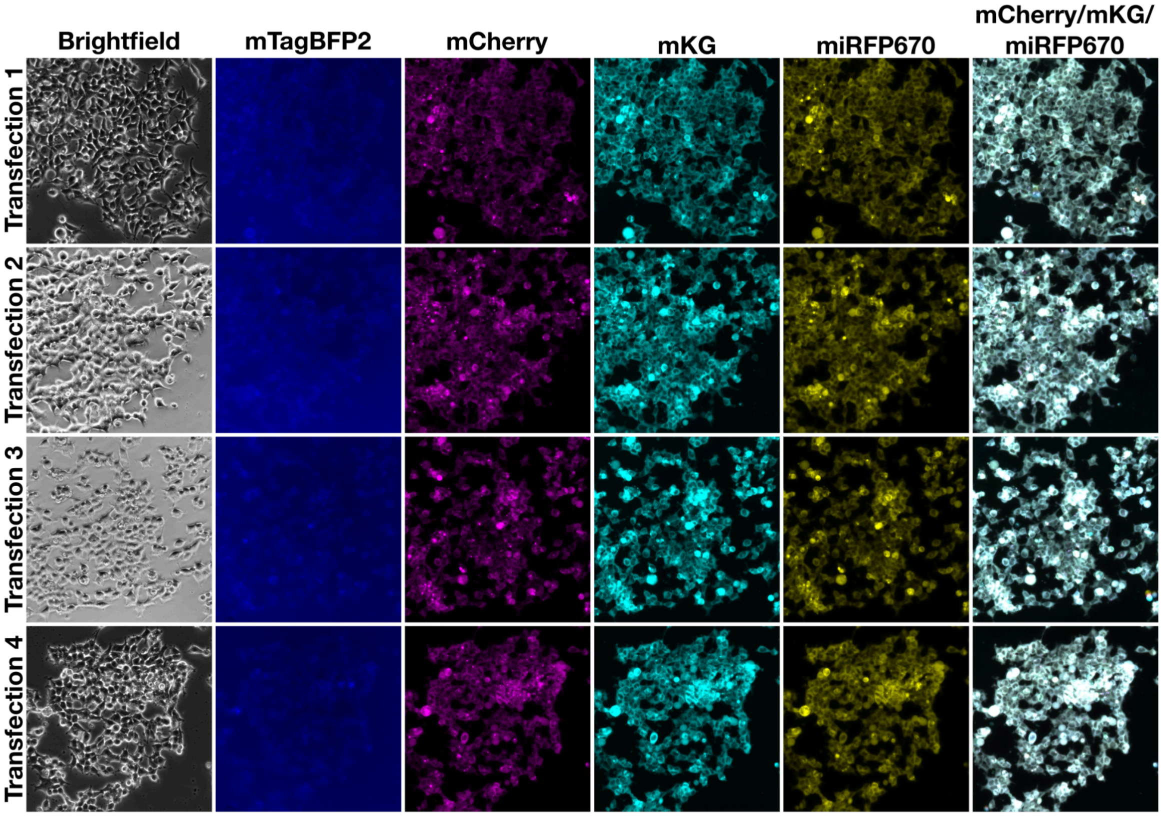


Supplemental Figure 11.

Microscope images of cells with integrated TR containing QTLNFDLLKLAGDVESNAGA ($F2A_{-4}NAGA_{+1}$) inserted at the 2A insertion site.

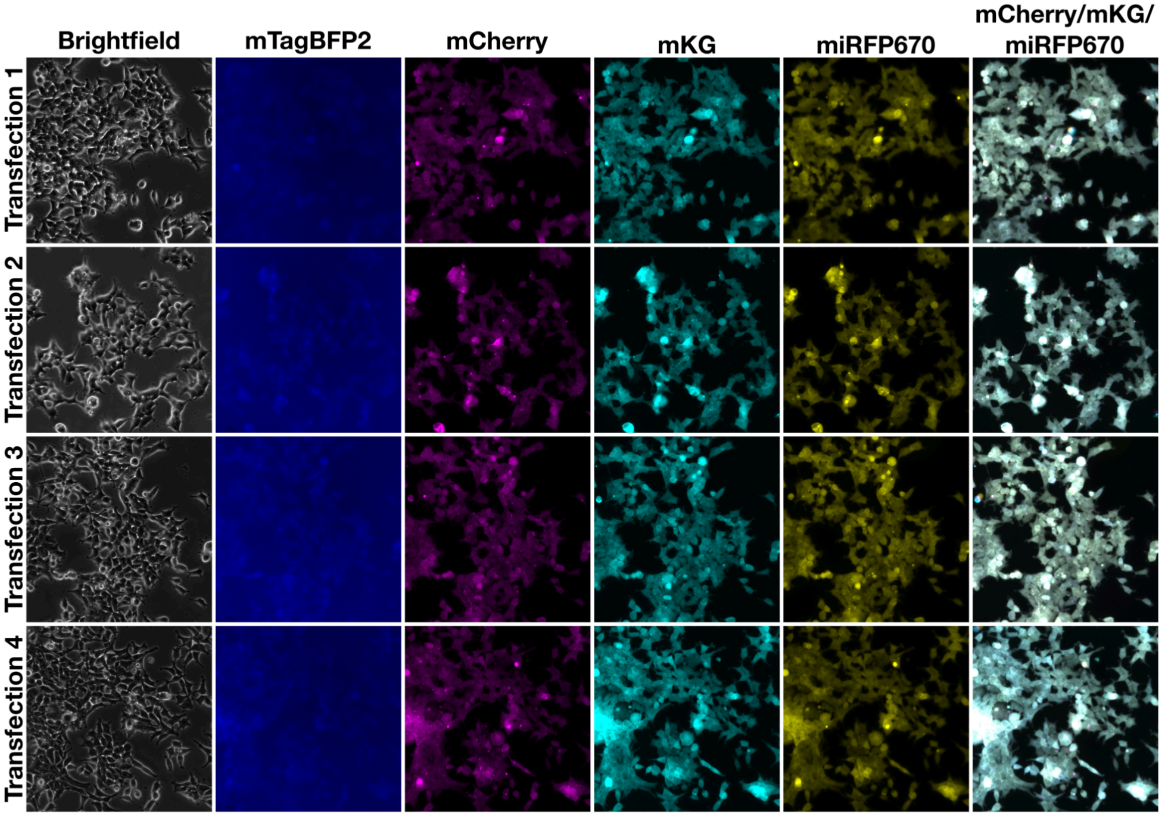


Supplemental Figure 12.

Microscope images of cells with integrated TR containing GSGSGSGSGSGSGSGSGSGS ((GS)x10) inserted at the 2A insertion site.

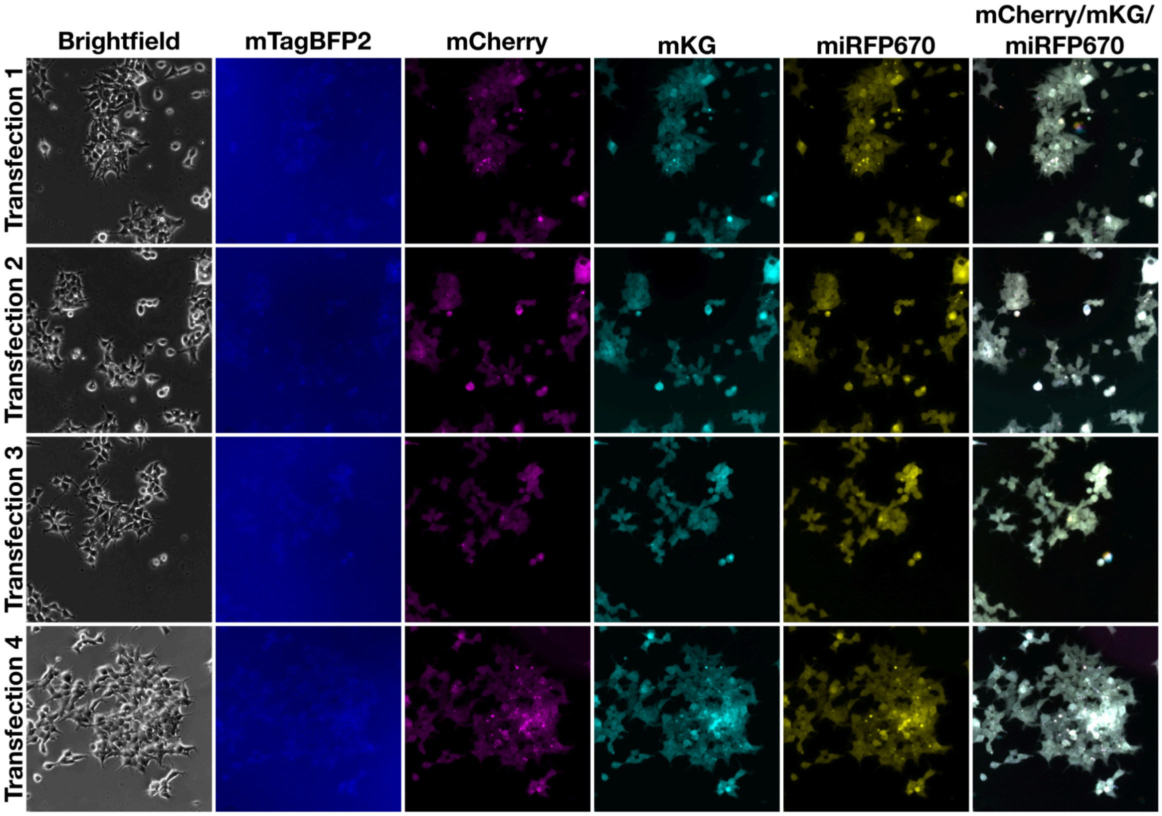


Supplemental Figure 13.

Microscope images of cells with integrated TR with no sequence inserted at the 2A insertion site.

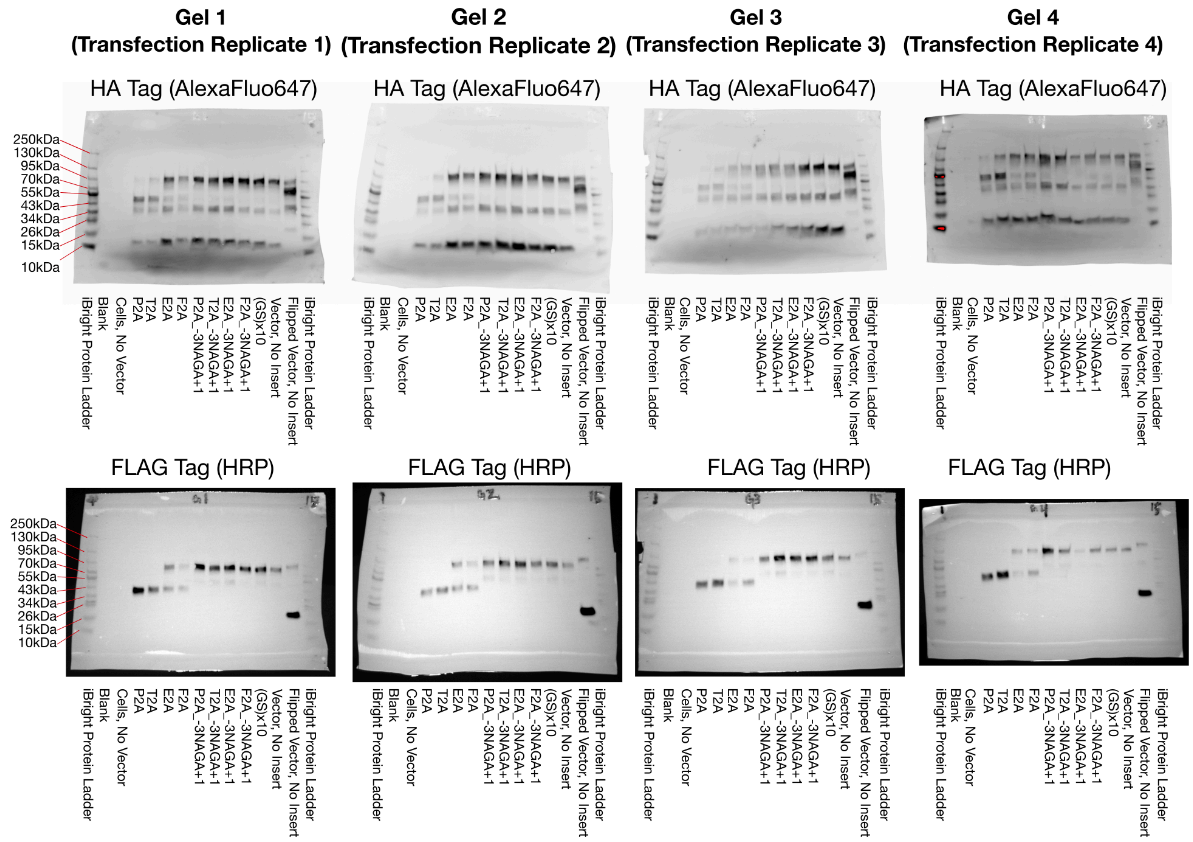


Supplemental Figure 14.

Western blots for samples from Figure 3 and Supplemental figures 3-13.

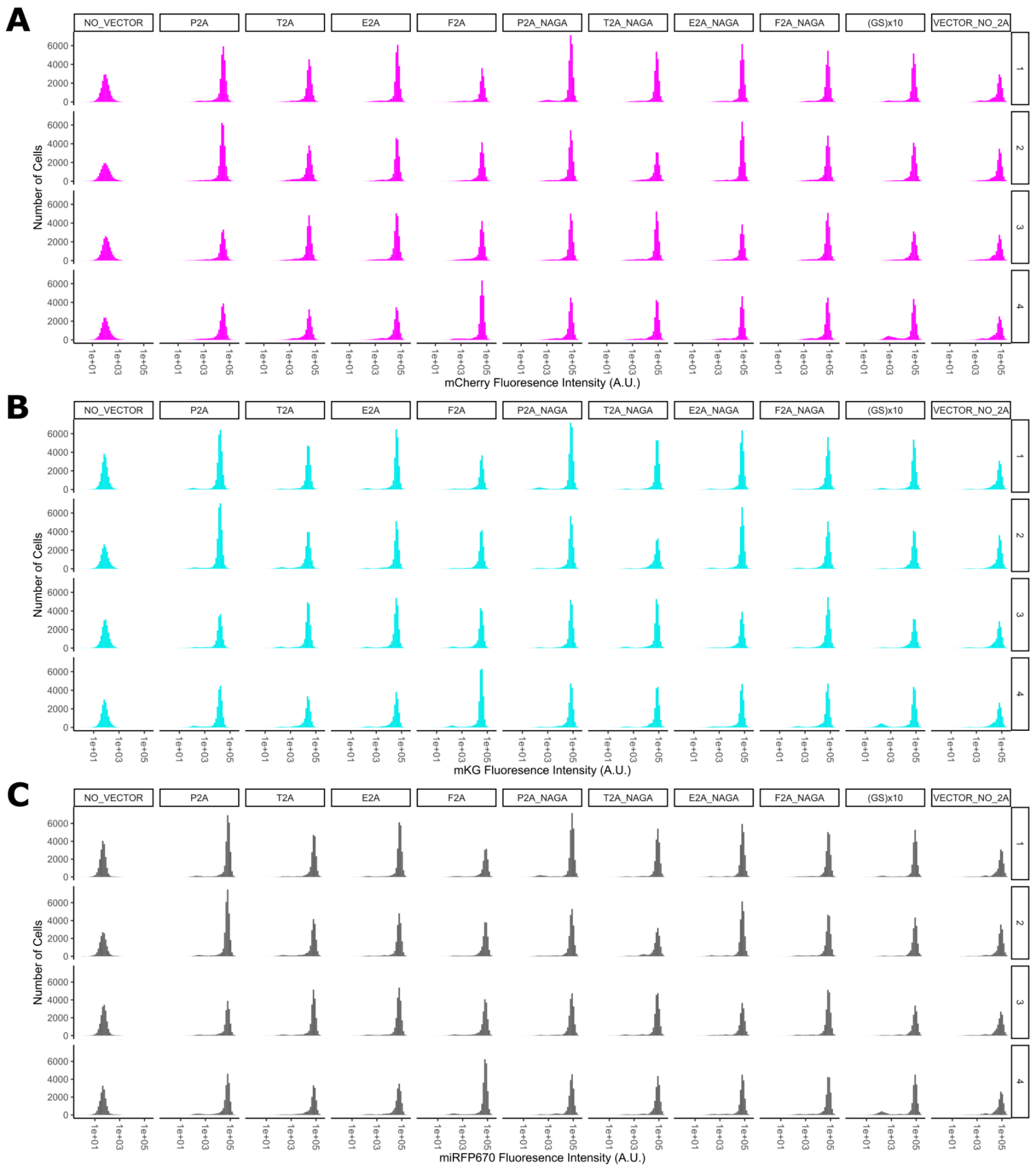


Supplemental Figure 15.

Flow Cytometry distributions for samples and replicates. (A) mCherry distribution, (B) mKG distribution, and (C) miRFP670 distribution for each sample used in Figure 3, and Supplemental figures 3-13. Replicates are independently transfected and maintained populations.

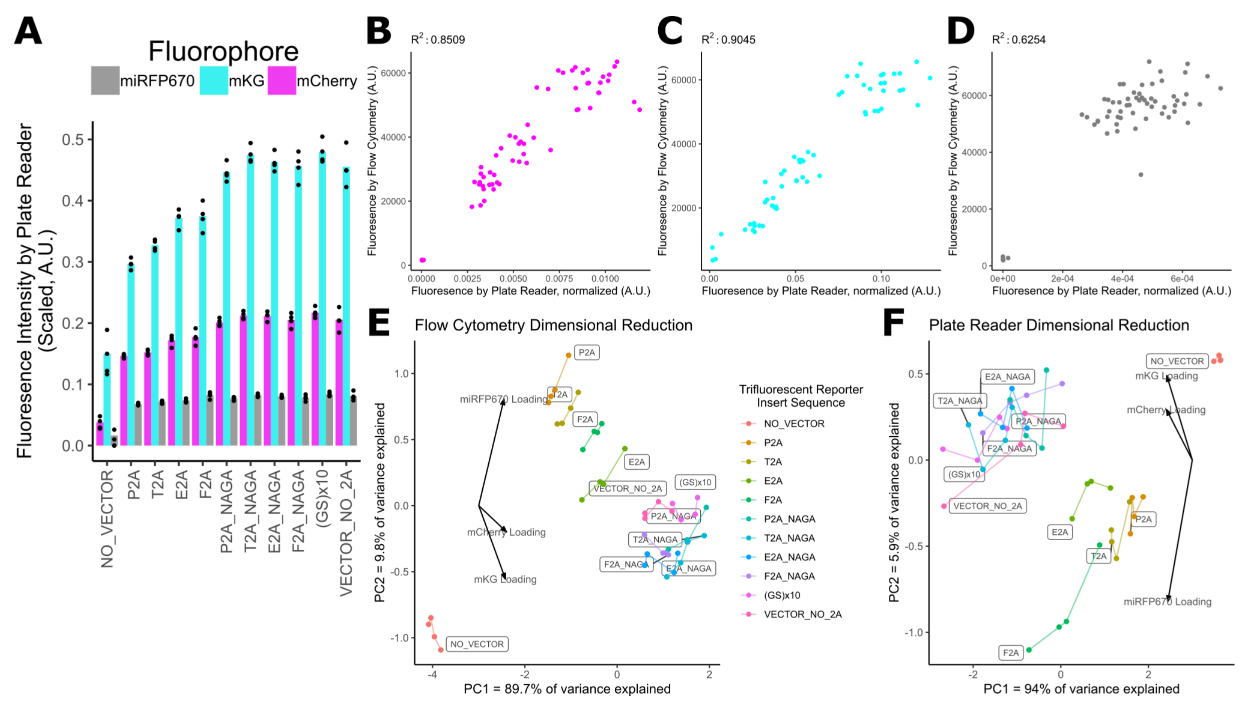


Supplemental Figure 16.

Measurement of Fluorescence through bulk population normalized to cell count on plate reader yields similar results to flow cytometry results, and fluorescent signal alone can identify skipping function. (A) Relative fluorescence of fluorescent intensity between samples, normalized to cell count. Scaled to cube root due to relatively lower sensitivity of instrument for miRFP670. Relationship between paired plate reader and flow cytometry measurements for (B) mCherry ($R^2$ = .8509), (C) mKG ($R^2$ = .9045), and (D) miRFP670 ($R^2$ = .6254) shows general correlation between the two methods, though lower correlation in miRFP670 is thought to be due to relative insensitivity of plate reader to miRFP670 fluorescent signal. (E) Dimensional reduction of flow cytometry measurements by Principle Component Analysis cleanly separates P2A, T2A, E2A, and F2A from non-skipping controls, with distance from non-skipping controls correlating with skipping efficacy. (F) Dimensional reduction of plate reader measurements by Principle Component Analysis cleanly separates P2A, T2A, E2A, and F2A from non-skipping controls, though distance from non-skipping controls does not directly correlate with skipping efficacy.

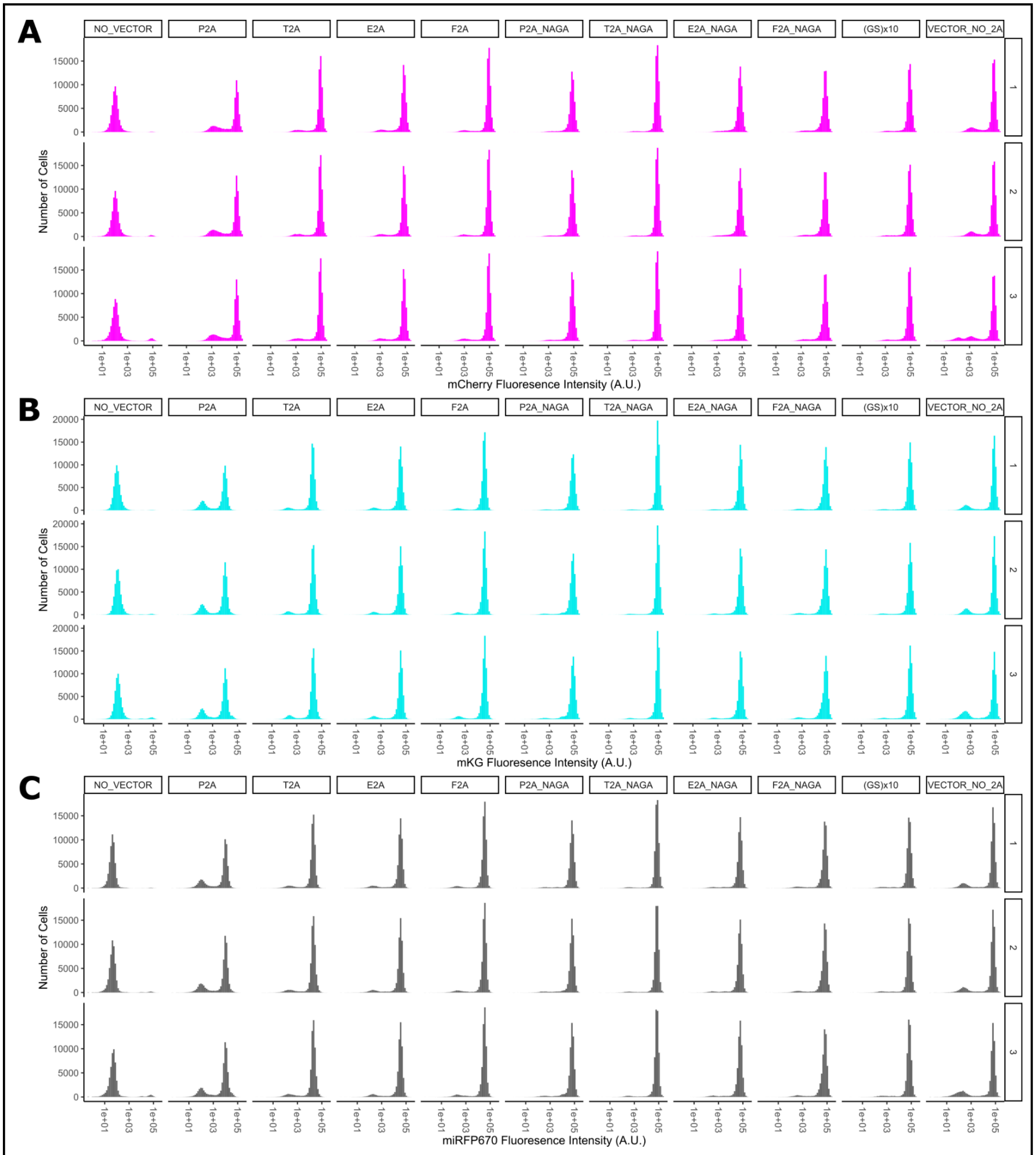


Supplemental Figure 17.

Flow Cytometry distributions for flipped trifluorescent reporter construct (miRFP670-mKG-mCherry). (A) mCherry distribution, (B) mKG distribution, and (C) miRFP670 distribution for flipped versions of constructs as used in figure 2. Replicates are technical replicates.

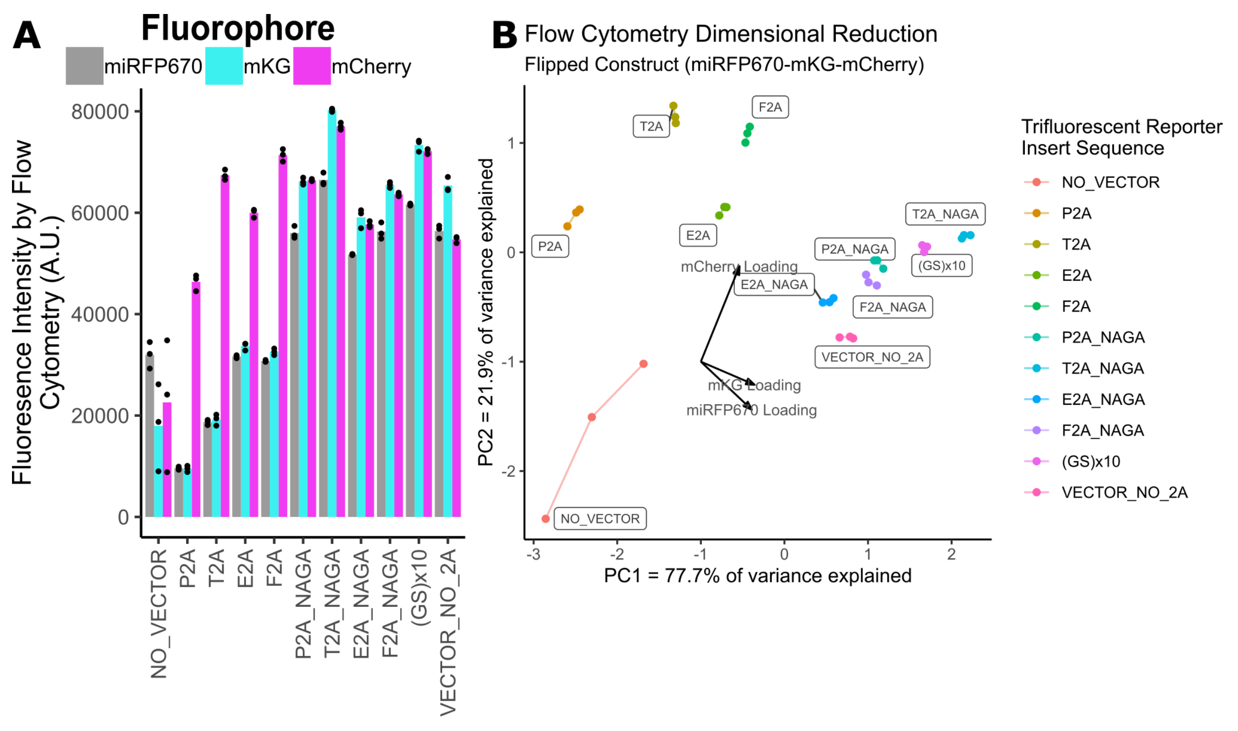


Supplemental Figure 18. Measurement and comparison of fluorescent signal from 2A insert panel from Figure 2 as applied to flipped trifluorescent reporter construct (miRFP670-mKG-mCherry). (A) Relative fluorescence of fluorescent intensity between samples. (B) Dimensional reduction of plate reader measurements by Principle Component Analysis. Distance of skipping samples (P2A, T2A, E2A, F2A) from non-skipping samples approximately correlates with expected skipping functionality.

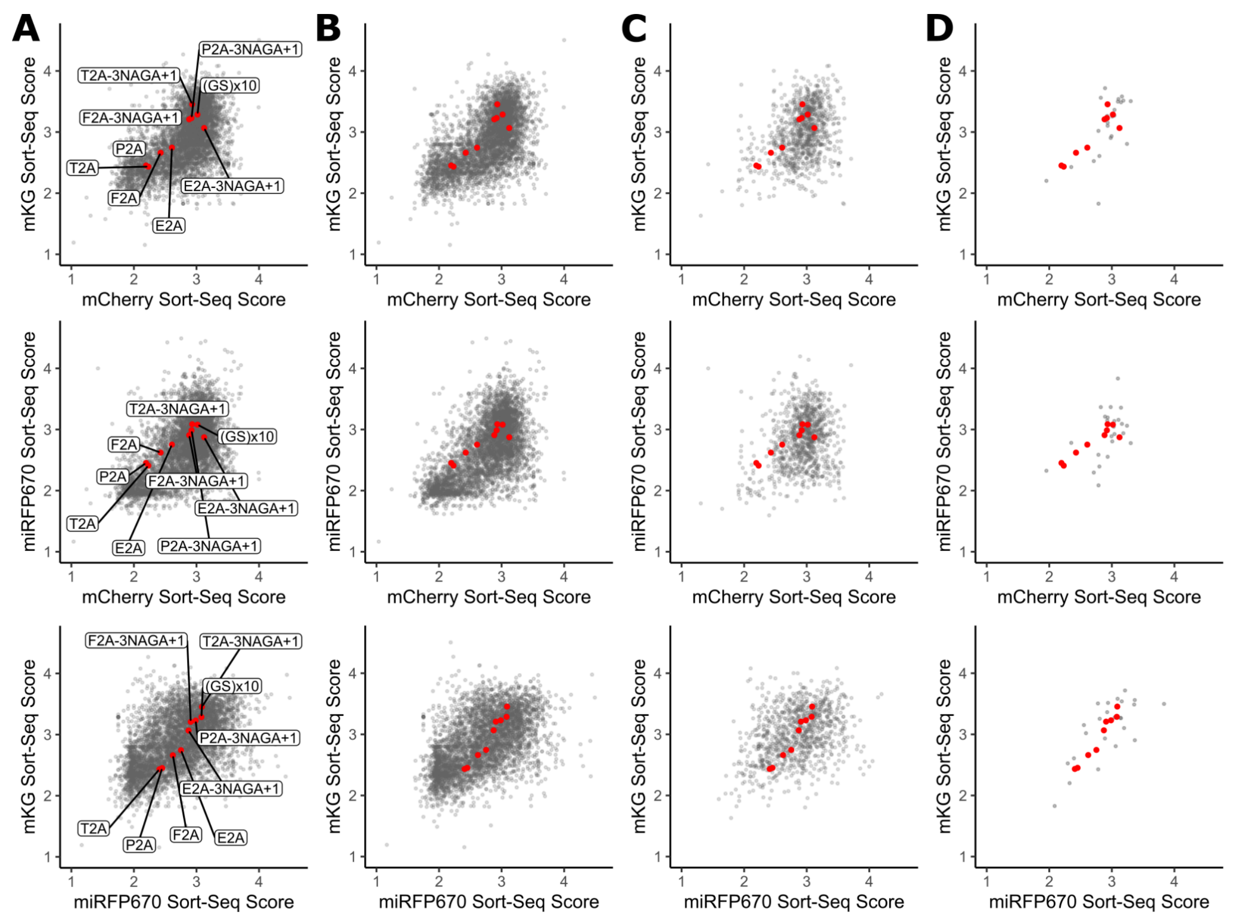


Supplemental Figure 19.

Pairwise comparisons for Sort-Seq color data. Top: mCherry and mKG. Middle: mCherry and miRFP670. Bottom: miRFP670 and mKG. (A) Full Sort-seq run. (B) Eukaryotic-origin sequences. (C) Bacerial-origin sequences. (D) Archaeal-origin sequences. Red dots represent control samples. P2A, T2A, E2A, and F2A are skipping controls. P2A$_{-3}$NAGA$_{+1}$, T2A$_{-3}$NAGA$_{+1}$, E2A$_{-3}$NAGA$_{+1}$, F2A$_{-3}$NAGA$_{+1}$, and (GS)x10 are non-skipping controls.

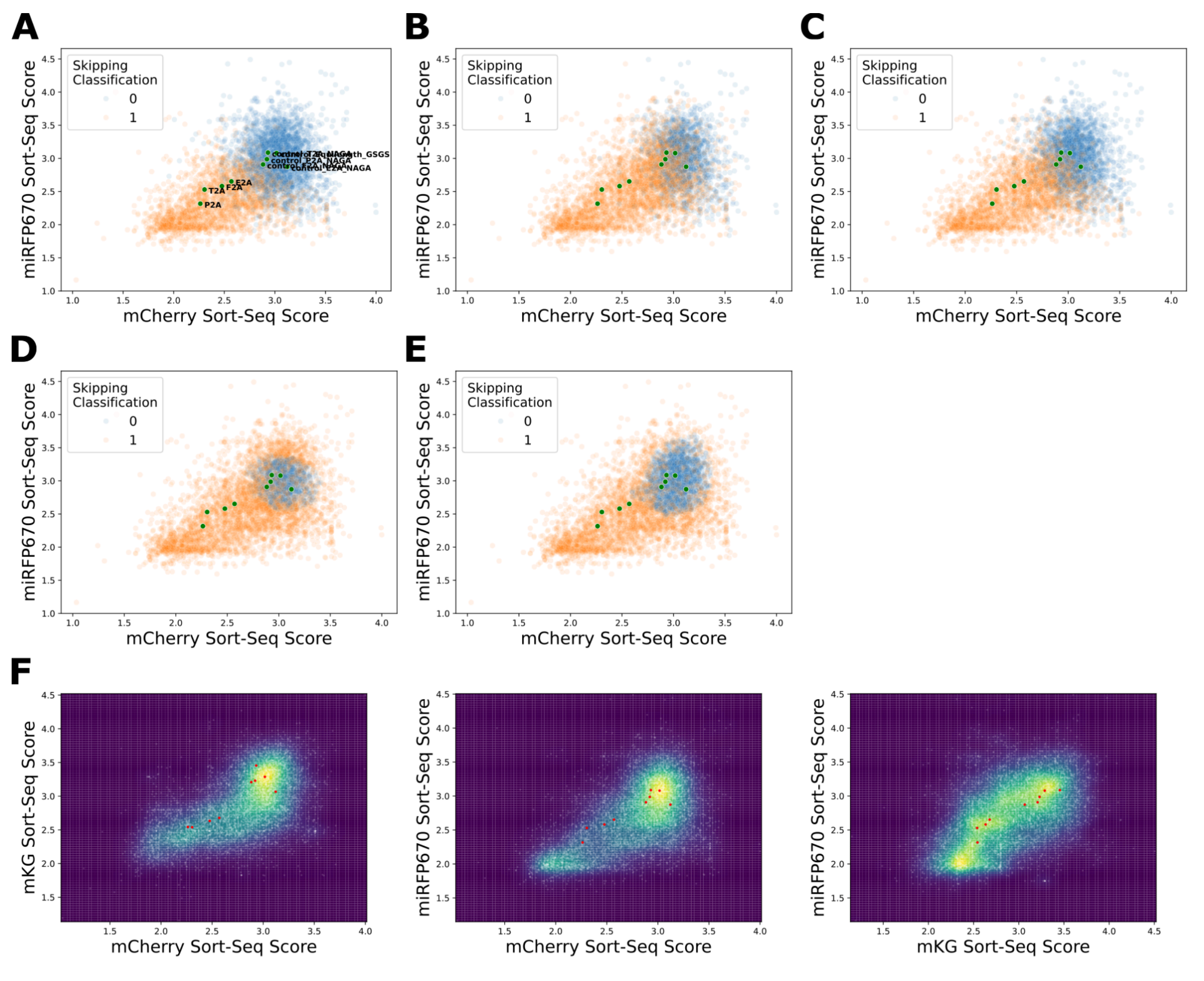


Supplemental Figure 20.

Results of classification models. All models were trained on the mCherry, mKG, and miRFP670 Sort-Seq scores for the controls, with skipping as a binary output.. (A) Fuzzy Clustering. (B) Support Vector Machine with linear kernel. (C) Support Vector Machine with polynomial kernel. (D) Support Vector Machine with radial basis function kernel. (E) Logistic Regression including the kernel density estimate as an input. (F) Kernel density estimate for the data cloud, shown as all pairwise fluorophore comparisons.

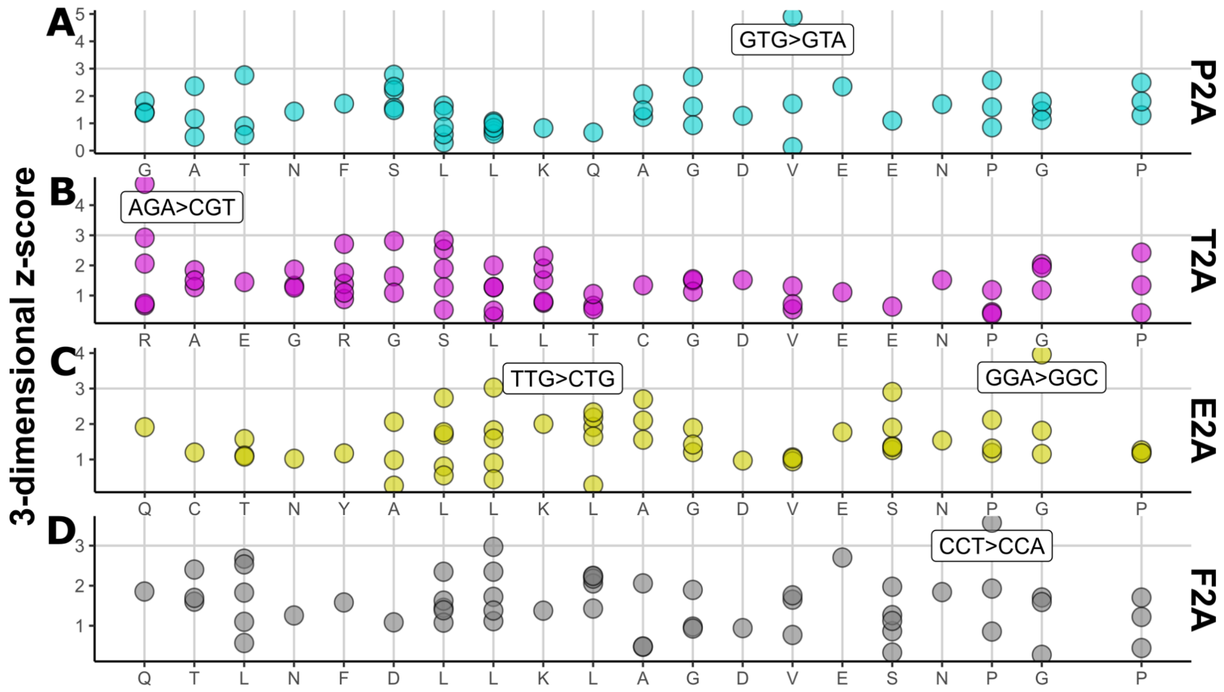


Supplemental Figure 21.

Z-scores of single-residue synonymous variants by residue. (A) Single-residue synonymous variants of P2A. (B) Single-residue synonymous variants of T2A. (C) Single-residue synonymous variants of E2A. (D) Single-residue synonymous variants of F2A.

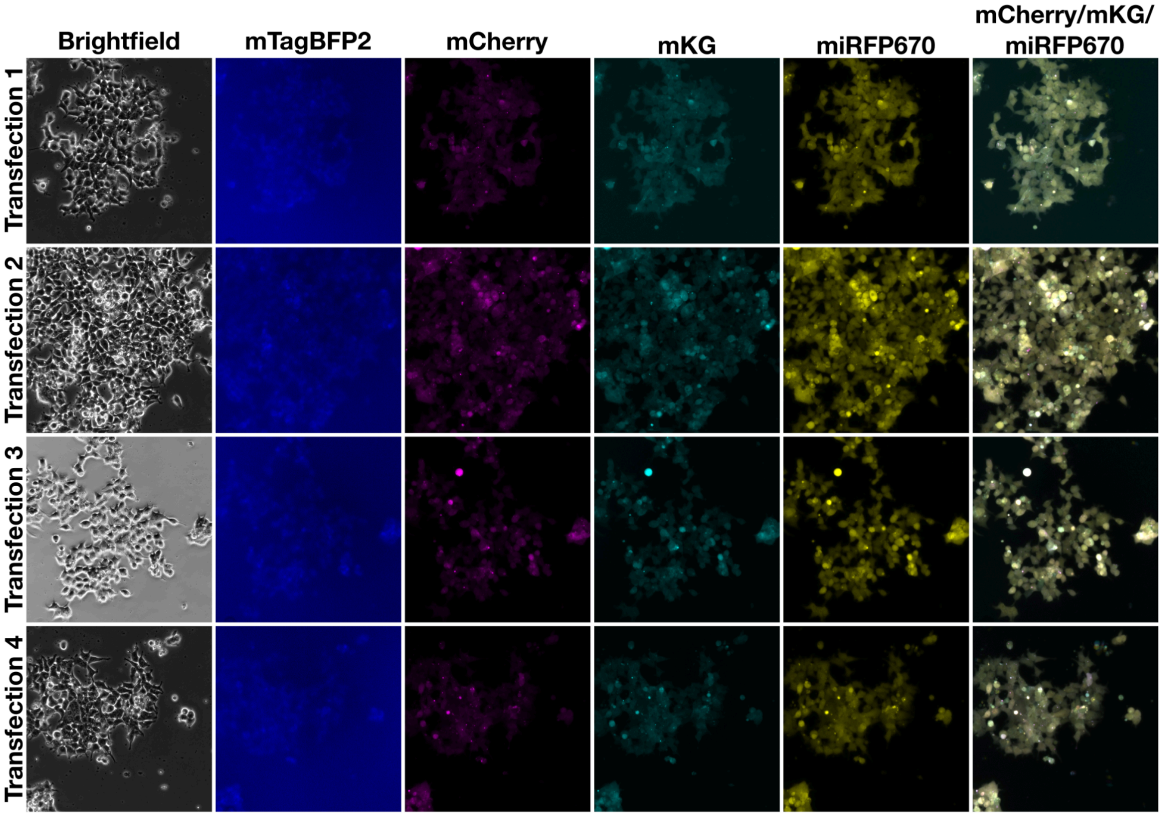


Supplemental Figure 22.

Microscope images of cells with integrated TR containing GATNFSLLKQAGDVEENPGP (P2A) where each codon is replaced with a synonymous variant inserted at the 2A insertion site.

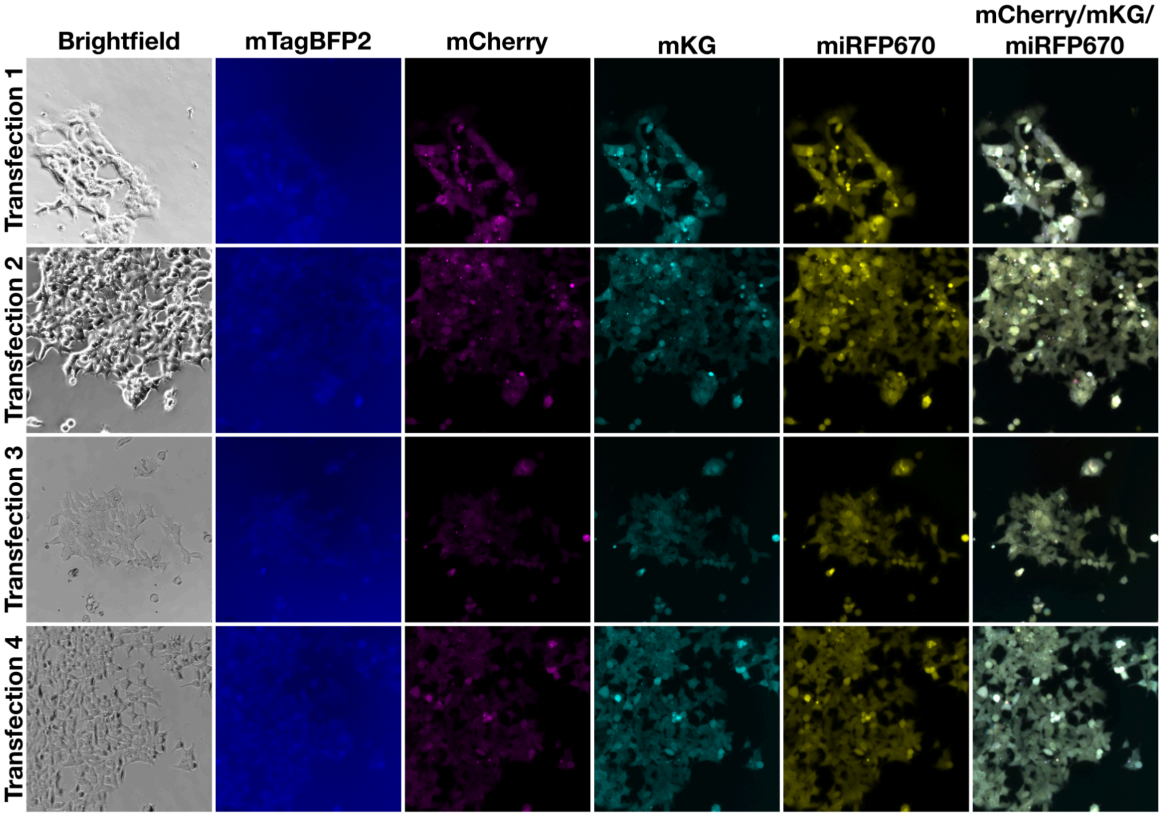


Supplemental Figure 23.

Microscope images of cells with integrated TR containing RAEGRGSLLTCGDVEENPGP (T2A) where each codon is replaced with a synonymous variant inserted at the 2A insertion site.

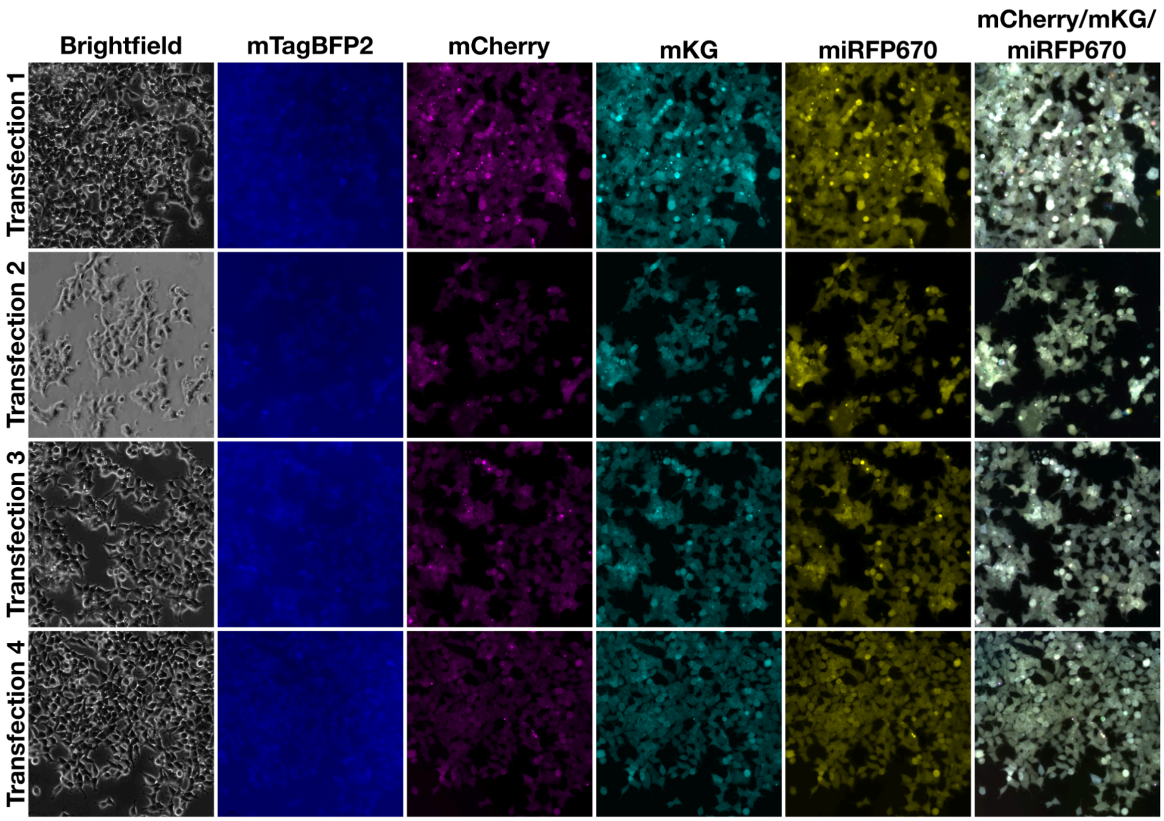


Supplemental Figure 24.

Microscope images of cells with integrated TR containing QCTNYALLKLAGDVESNPGP (E2A) where each codon is replaced with a synonymous variant inserted at the 2A insertion site.

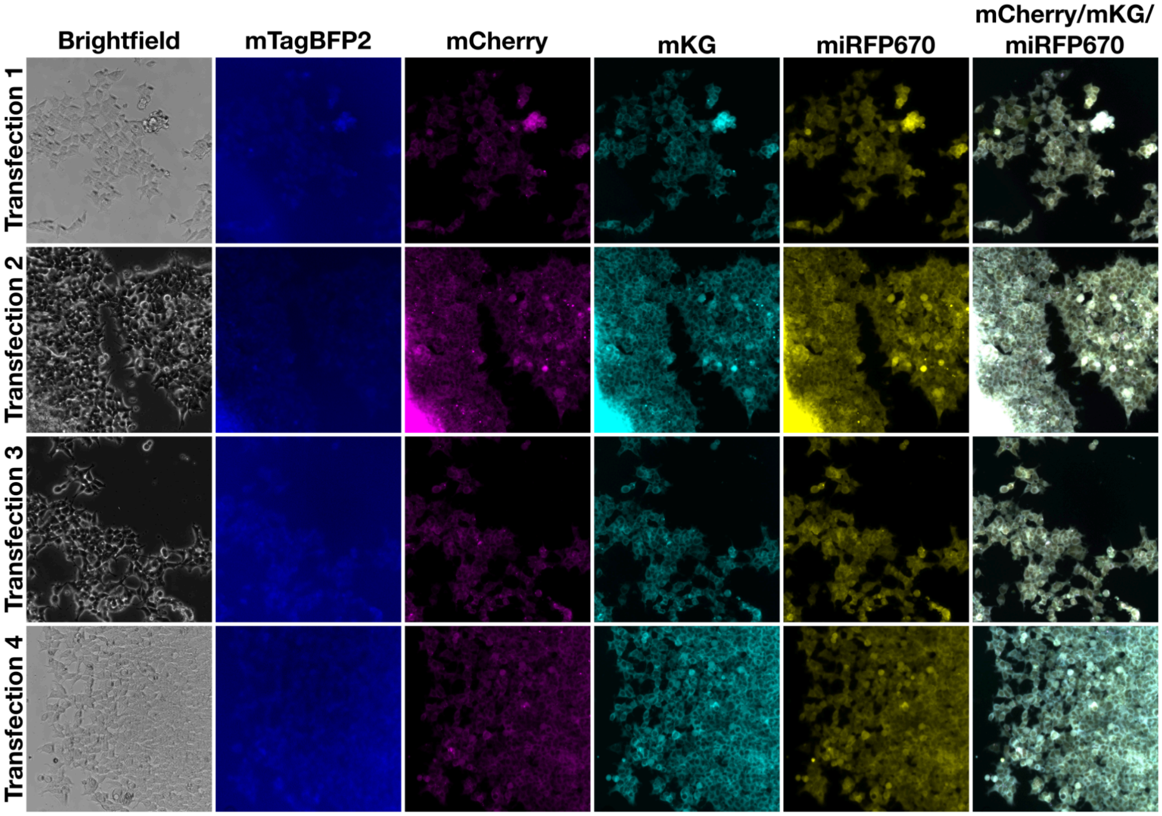


Supplemental Figure 25.

Microscope images of cells with integrated TR containing QTLNFDLLKLAGDVESNPGP (F2A) where each codon is replaced with a synonymous variant inserted at the 2A insertion site.

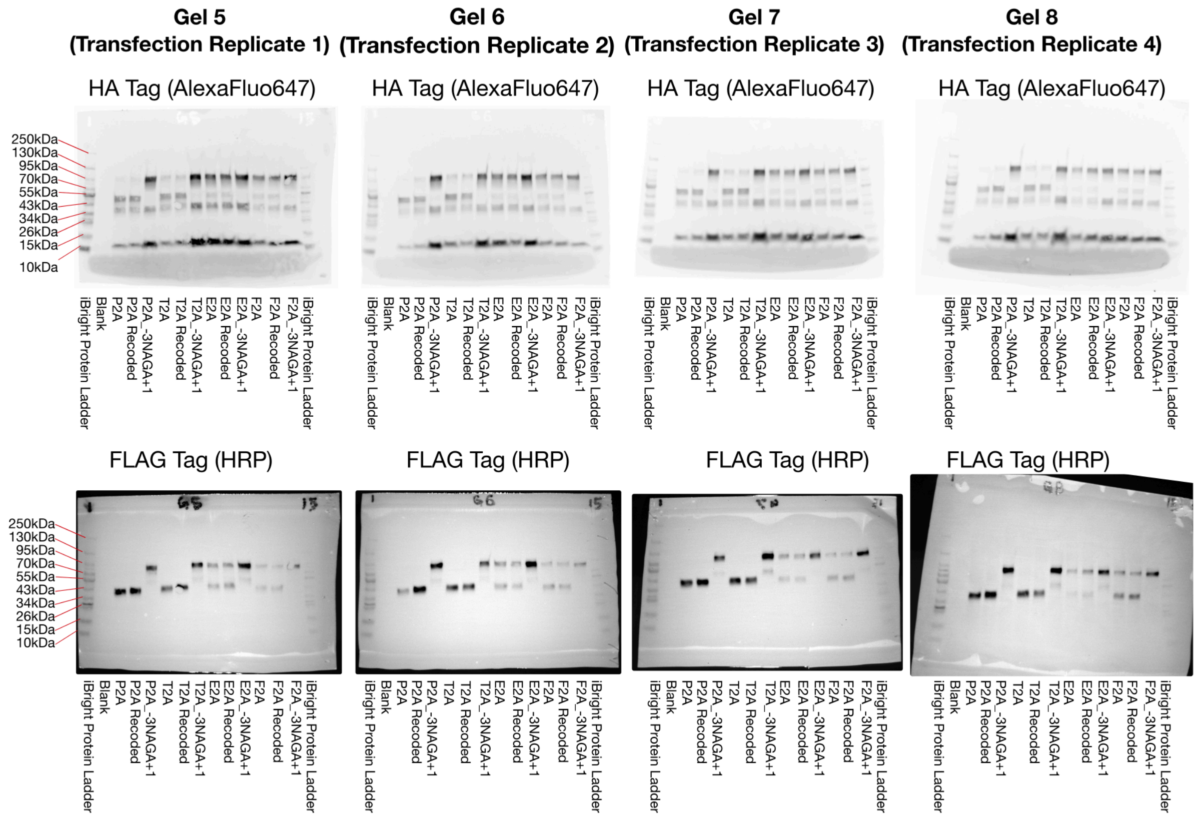


Supplemental Figure 26.

Samples for recoded variants of P2A, T2A, E2A, and F2A as tested in Figure 5 and Supplemental Figure 21-24. Samples of P2A, T2A, E2A, and F2A, as well as the $_{-3}$NAGA$_{+1}$ variants as used in Figure 2 are included for comparison.

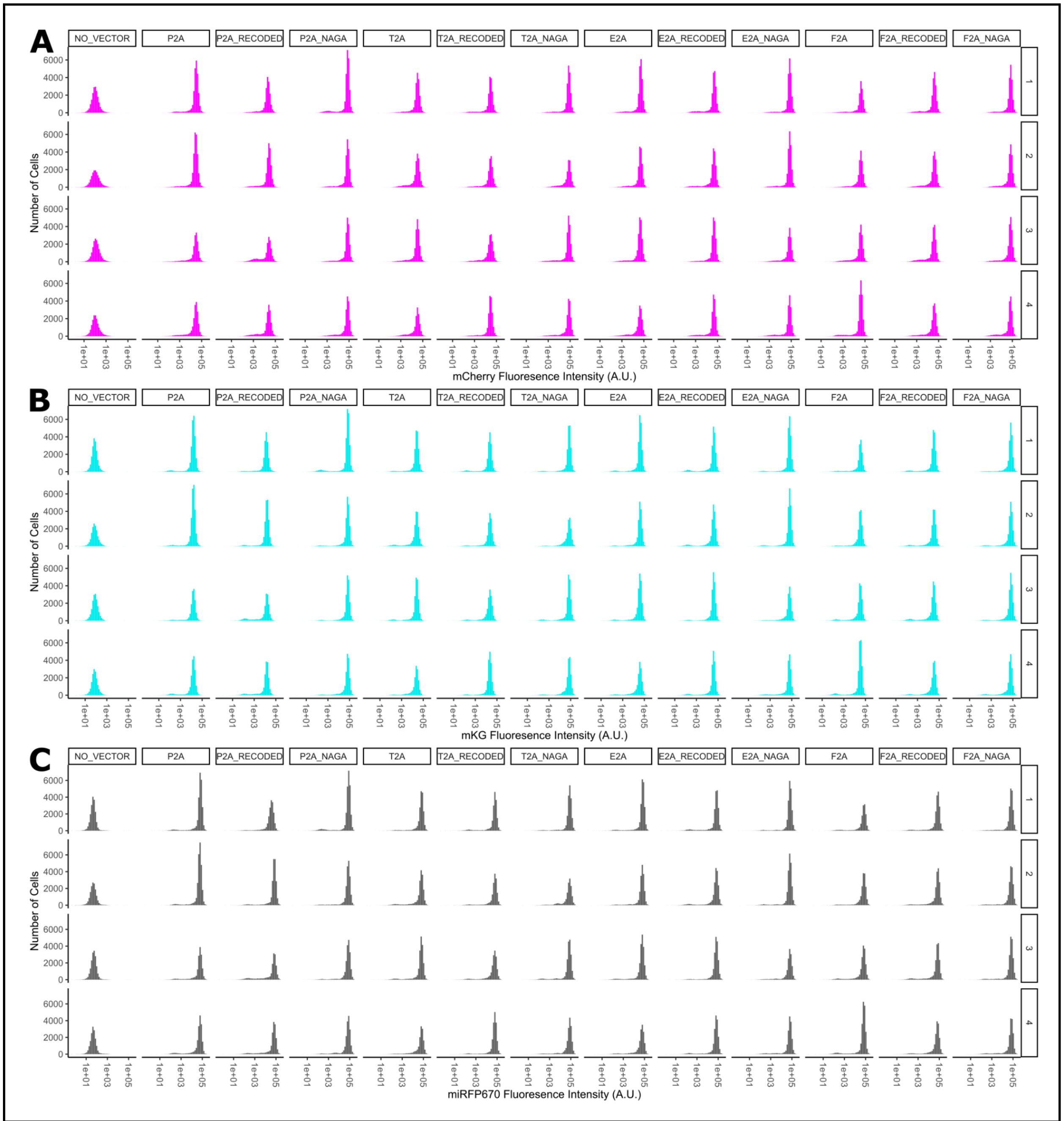


Supplemental Figure 27.

Flow cytometry distributions for fully recoded variants of P2A, T2A, E2A, and F2A. (A) mCherry distribution, (B) mKG distribution, and (C) miRFP670 distribution for samples used for Figure 5. Distributions for original coding sequences, cells without a vector, and $_{-3}$NAGA$_{+1}$ variants are included for reference.

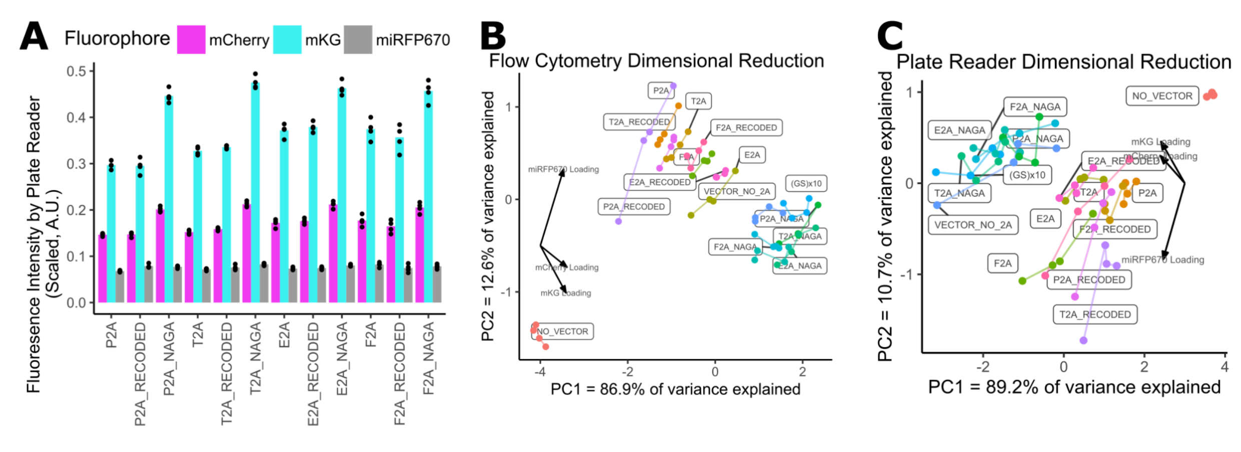


Supplemental Figure 28.

Additional Measurements of Recoded Variants of P2A, T2A, E2A, and F2A. (A) Fluorescence intensity of cell population normalized to cell count, with application of cube root scaling. (B) Principle Component Analysis of control samples and fully recoded variants of P2A, T2A, E2A, and F2A based on results from Flow Cytometry measurements. (C) Principle Component Analysis of control samples and fully recoded variants of P2A, T2A, E2A, and F2A based on results from plate reader measurements.

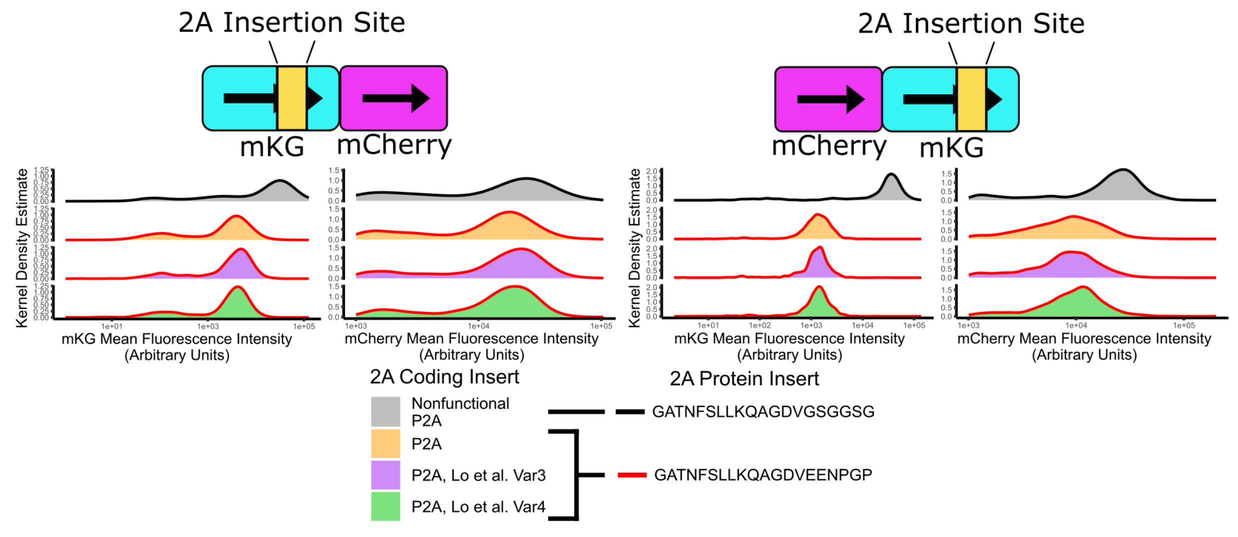


Supplemental Figure 29.

We were unable to recapitulate the loss of skipping phenotype reported in Lo et al. 2015. When we compare the coding sequence of functional P2A, P2A with the coding sequence of Var3 (which is reported to skip) and P2A with the coding sequence of Var4 (which is reported to not skip), we observe matching fluorescent phenotypes.

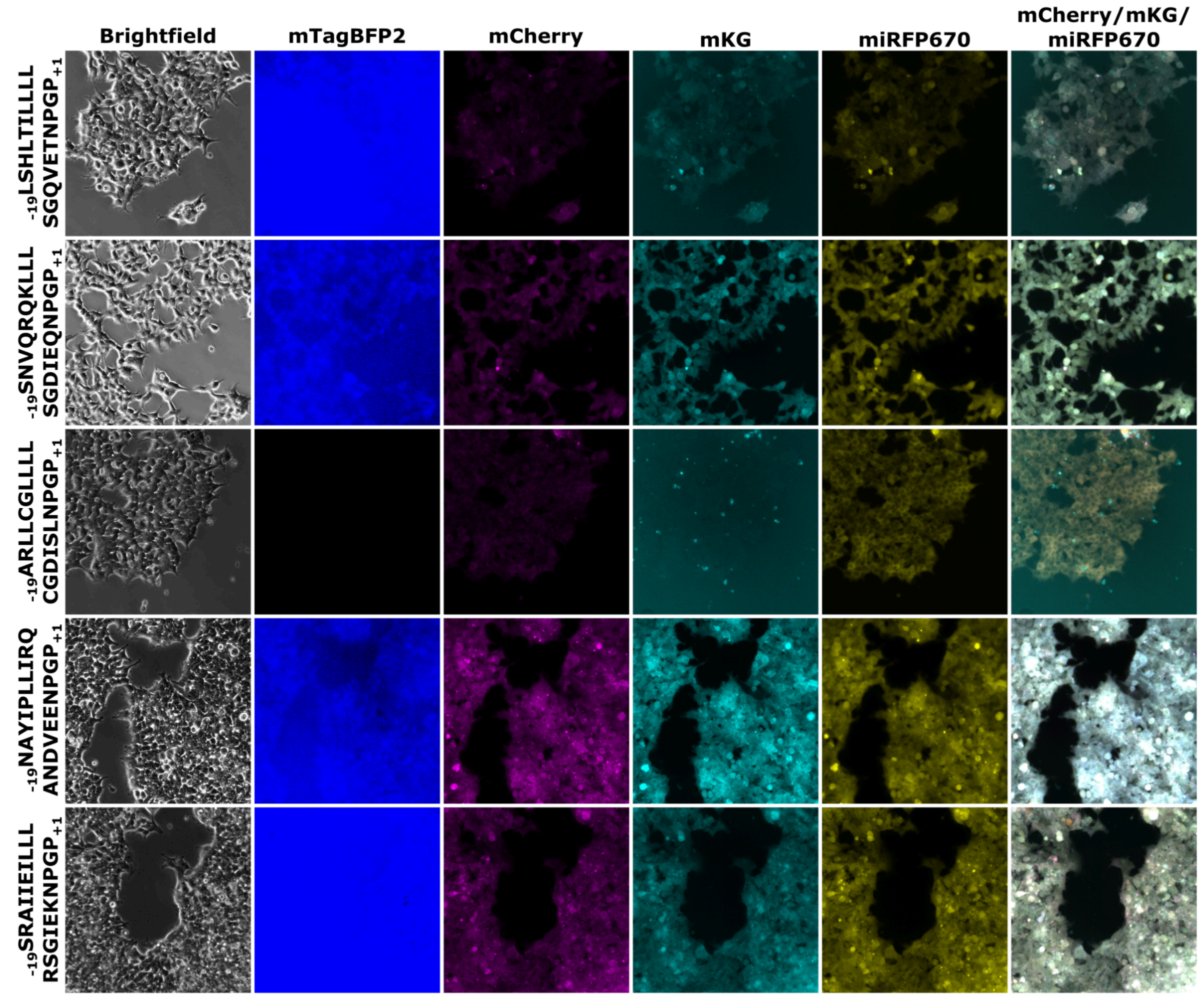


Supplemental Figure 30.

Microscope images of cells with integrated trifluorescent construct containing $_{-19}$LSHLTILLLLSGVETNPGP$_{+1}$, $_{-19}$SNVQRQKLLLSGDIEQNPGP$_{+1}$, $_{-19}$ARLLCGLLLLCGDISLNPGP$_{+1}$, $_{-19}$NAYIPLLIRQANDVEENPGP$_{+1}$, or $_{-19}$SRAIIEILLLRSGIEKNPGP$_{+1}$.

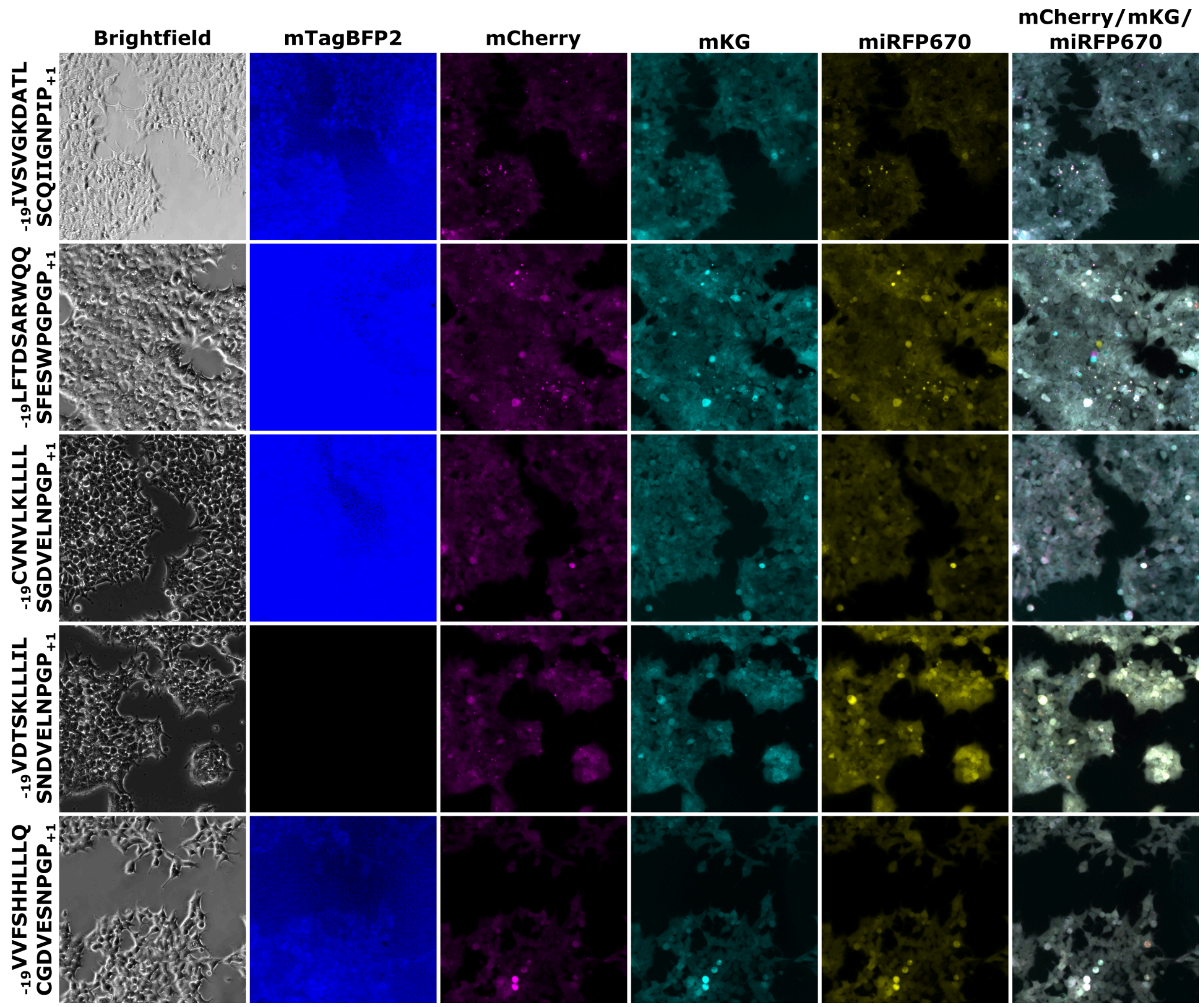


Supplemental Figure 31.

Microscope images of cells with integrated trifluorescent construct containing $_{-19}$IVSVGKDATLSCQIIGNPIP$_{+1}$, $_{-19}$LFTDSARWQQSFESWPGPGP$_{+1}$, $_{-19}$CVNVLKLLLLSGDVELNPGP$_{+1}$, $_{-19}$VDTSKLLLILSNDVELNPGP$_{+1}$, or $_{-19}$VVFSHHLLLQCGDVESNPGP$_{+1}$.

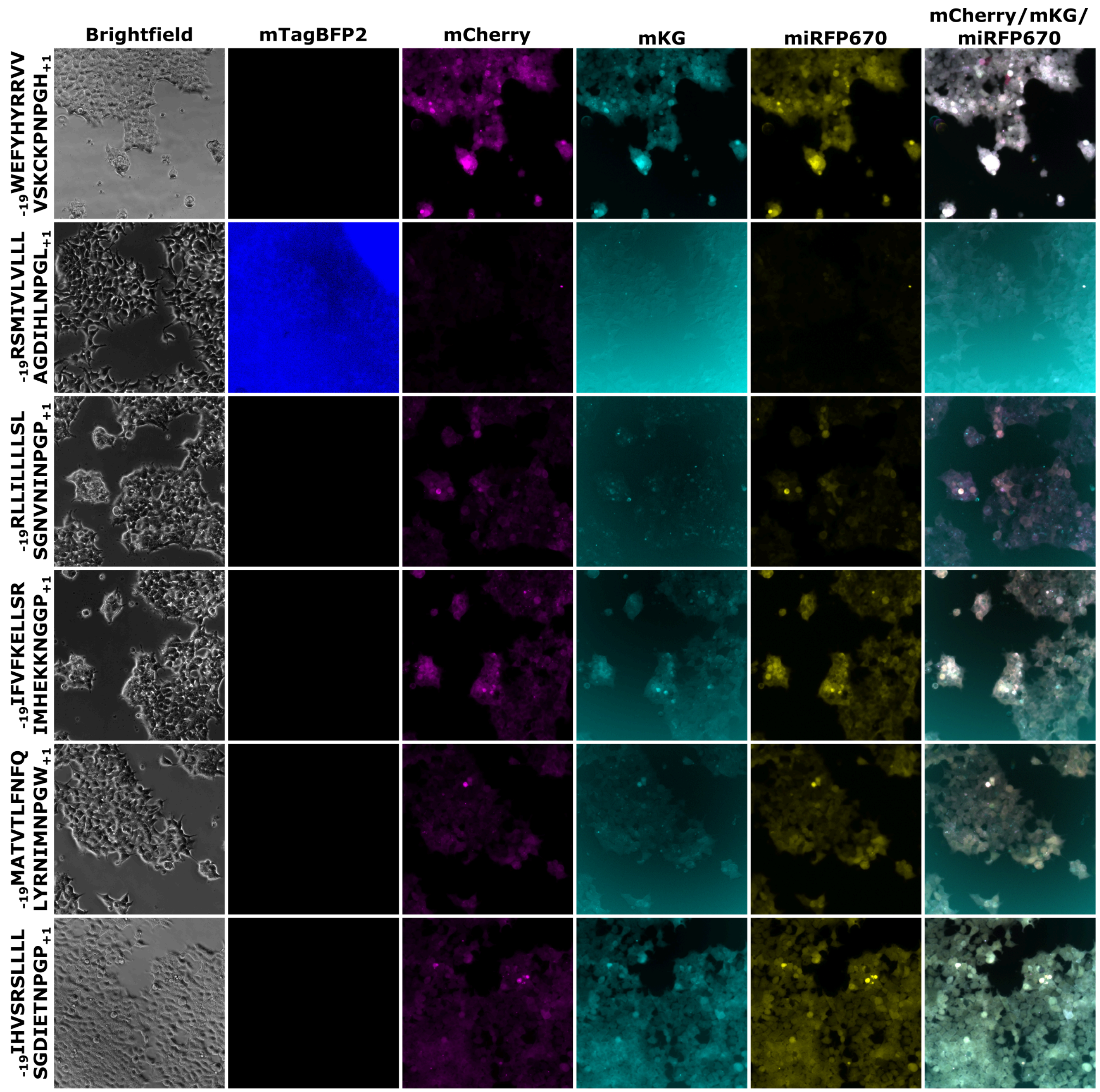


Supplemental Figure 32.

Microscope images of cells with integrated trifluorescent construct containing $_{-19}$WEFYHYRRVVVSKCKPNPGH$_{+1}$, $_{-19}$RSMIVLVLLLADGIHLNPGL$_{+1}$, $_{-19}$RLLILLLLSLSGNVNINPGP$_{+1}$, $_{-19}$IFVFKELLSRIMHEKKNGGP$_{+1}$, $_{-19}$MATVTLFNFQLYRNIMNPGW$_{+1}$, or $_{-19}$IGVSRSLLLLSGDIETNPGP$_{+1}$.

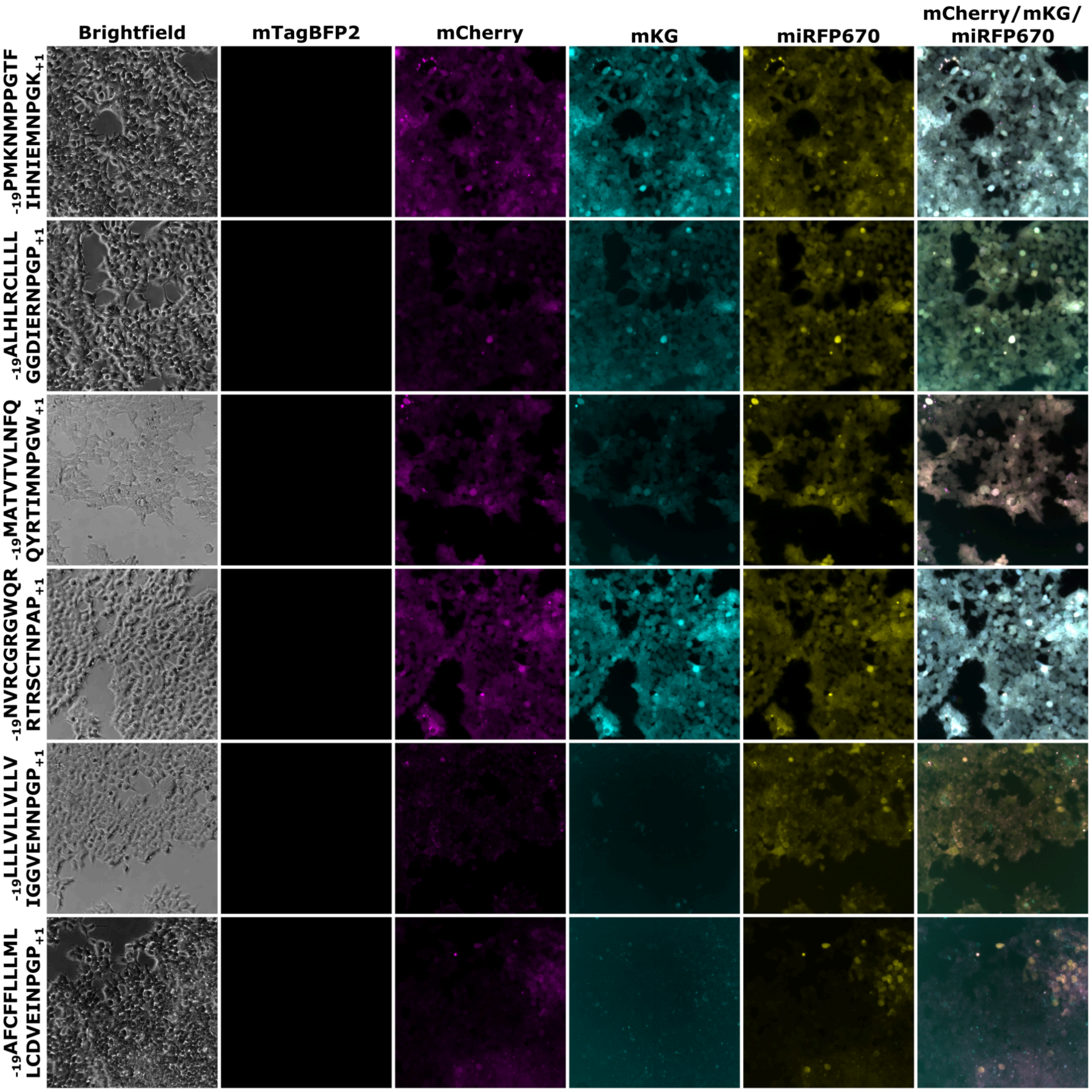


Supplemental Figure 33.

Microscope images of cells with integrated trifluorescent construct containing $_{-19}$PMKNMPPGTFIGNIEMNPGK$_{+1}$, $_{-19}$ALHLRCLLLLGGDIERNPGP$_{+1}$, $_{-19}$MATVTVLNFQQYRTIMNPGW$_{+1}$, $_{-19}$NVRCGRGWQRRTRSCTNPAP$_{+1}$, $_{-19}$LLLVLLVLLVIGGVEMNPGP$_{+1}$, or $_{-19}$AFCFFLLLMLLCDVEINPGP$_{+1}$. $_{-19}$AFCFFLLLMLLCDVEINPGP$_{+1}$ is an artificial 2A-like protein designed based on positional residue enrichment in all classified functionally active 2A-like peptides.

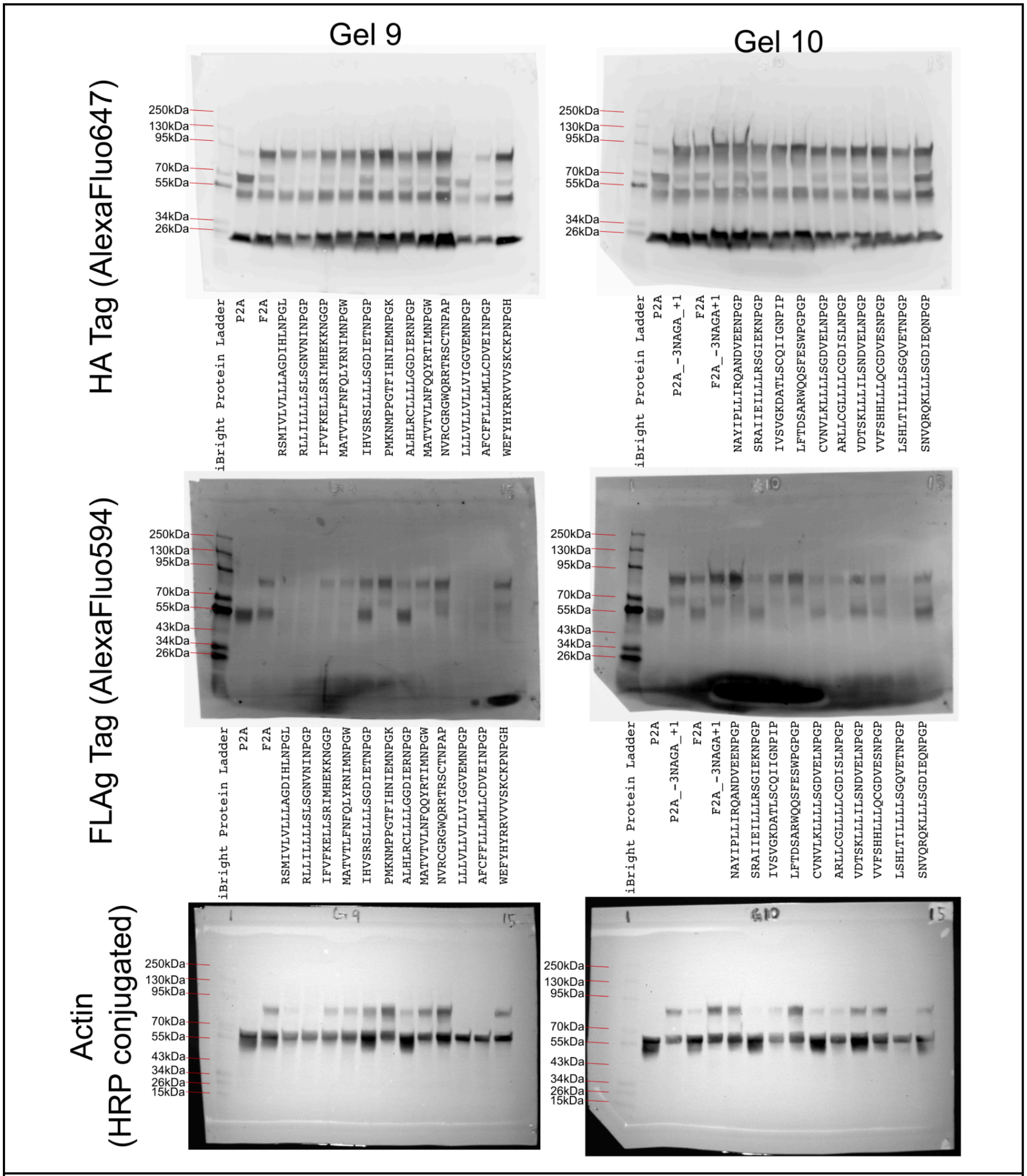


Supplemental Figure 34.

Full Western Blots for samples measured in Figure 7. Samples were loaded with equal amounts of protein. The anti-Actin antibody appeared to have conjugated with the FLAG tag as well as Actin, leading to a seeming additive band pattern.

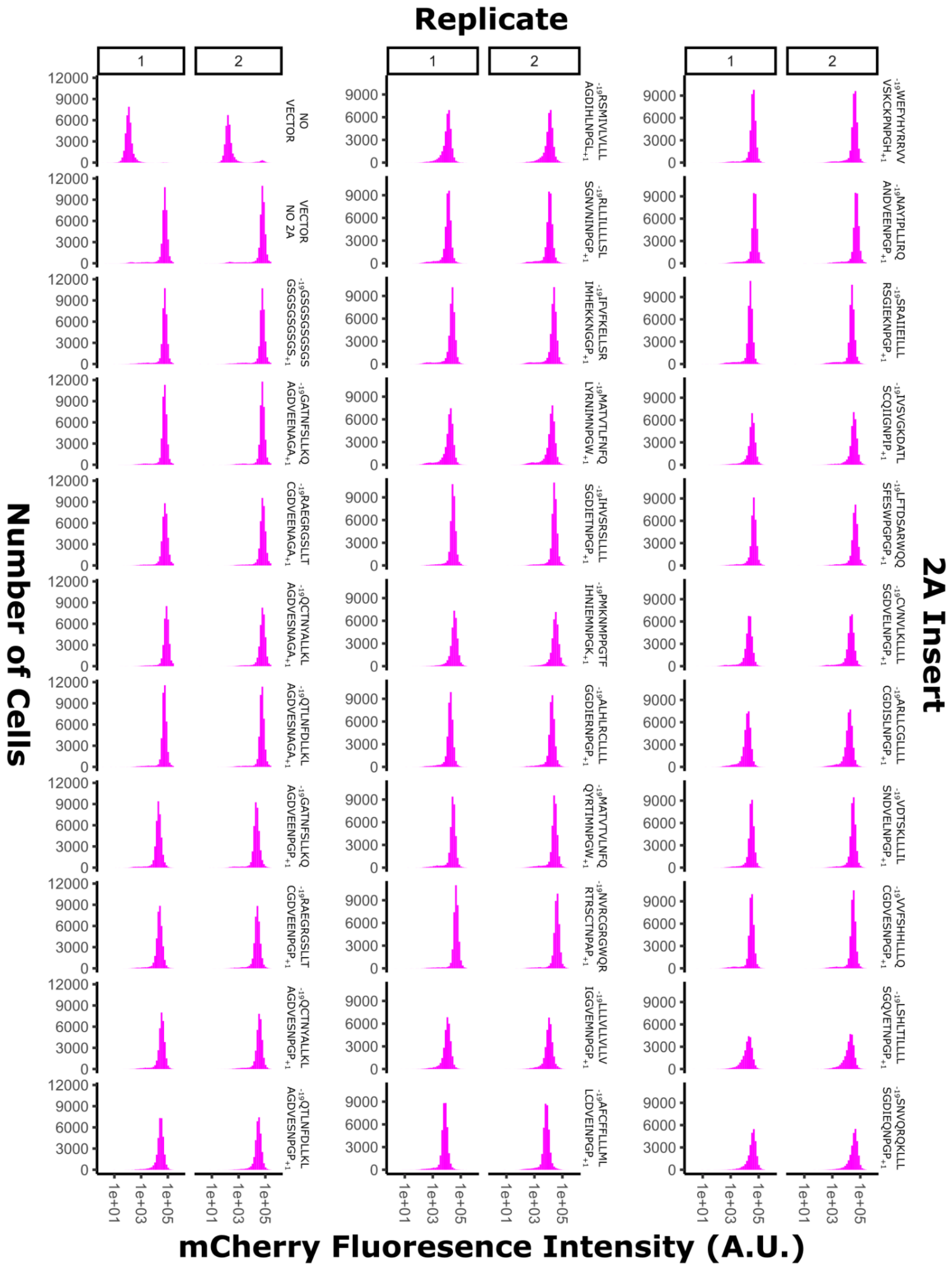


Supplemental Figure 35.

mCherry flow cytometry distributions for samples from Figure 7. Replicates are technical replicates.

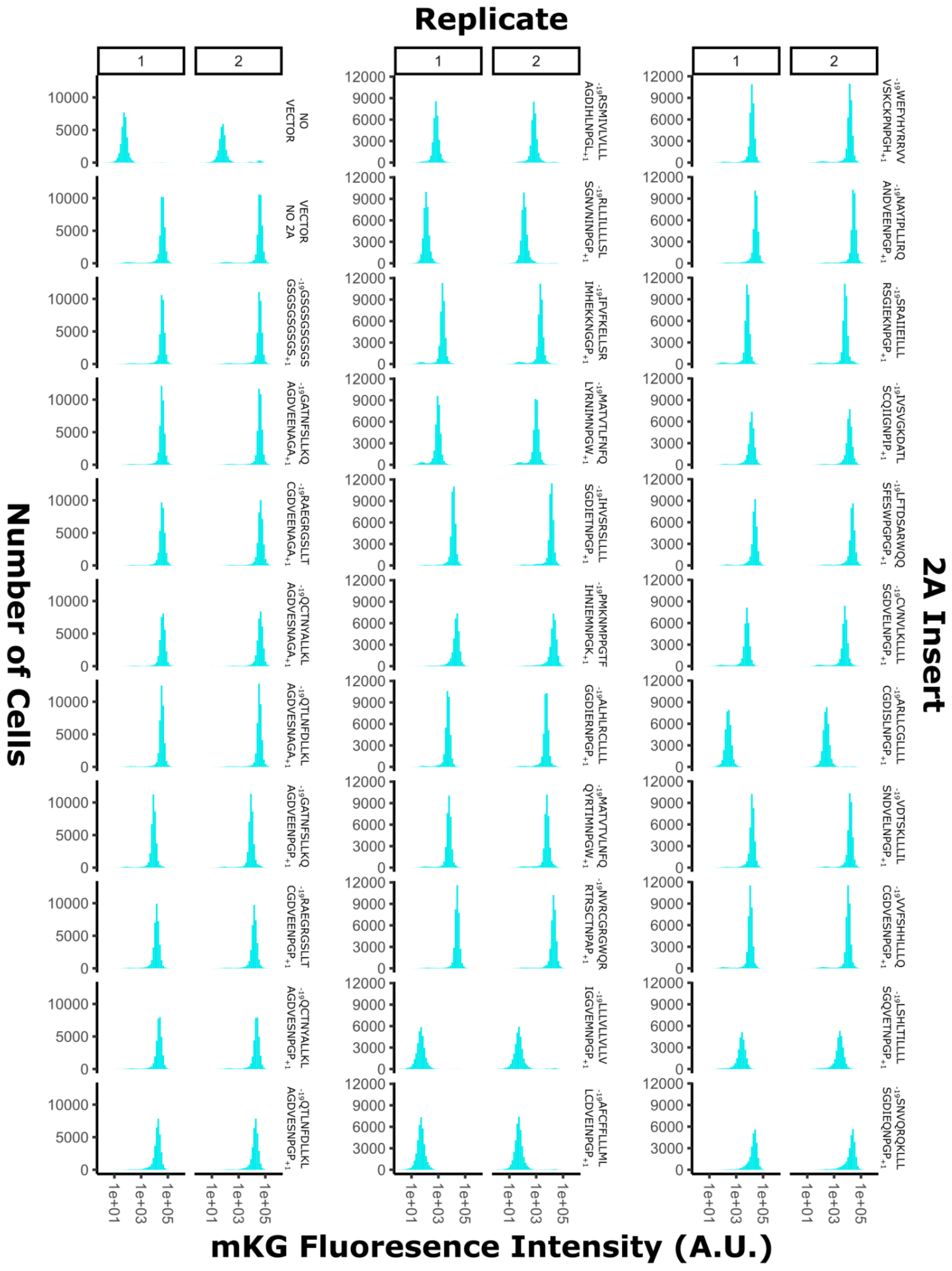


Supplemental Figure 36.

mKG flow cytometry distributions for samples from Figure 7. Replicates are technical replicates.

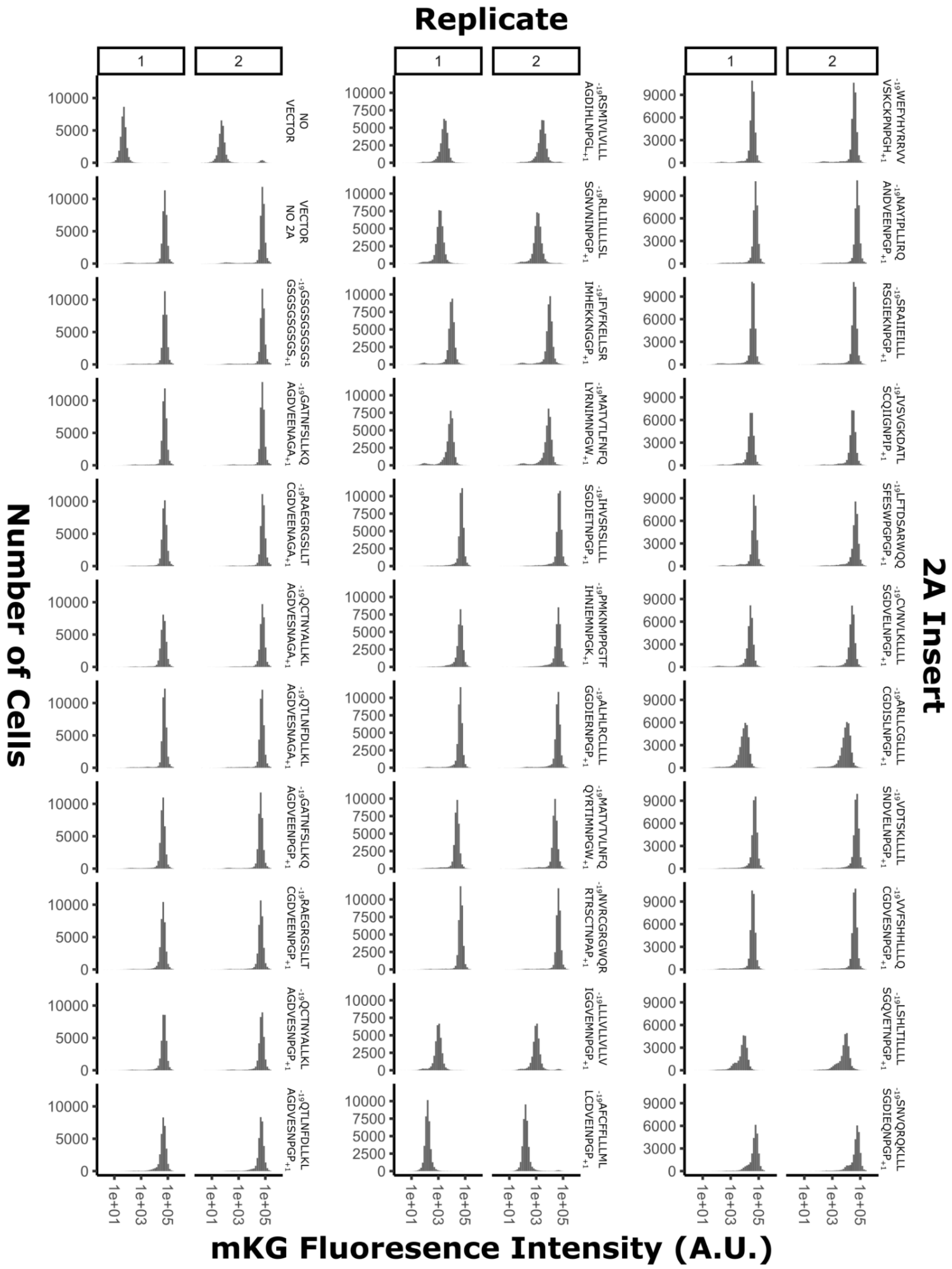


Supplemental Figure 37.

miRFP670 flow cytometry distributions for samples from Figure 7.  Replicates are technical replicates.

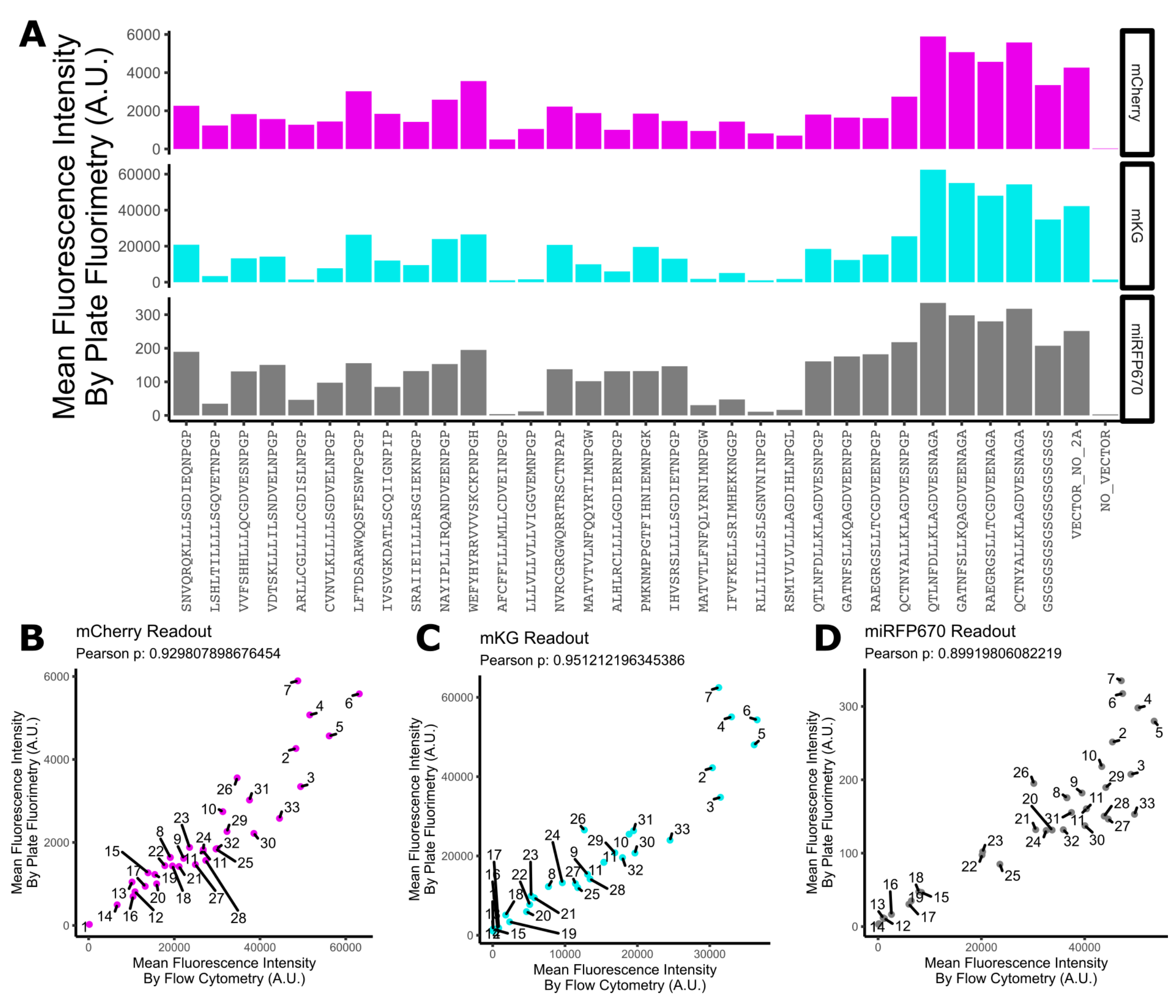


Supplemental Figure 38.

Plate Fluorimetry results for samples from Figure 7. (A) Plate Fluorimetry results for mCherry (top), mKG (middle), and miRFP670 (bottom). (B) Correlation of flow cytometry and plate fluorimetry results for mCherry shows effectively identical results based on fluorescence measurement approach. (C) Correlation of flow cytometry and plate fluorimetry results for mKG shows effectively identical results based on fluorescence measurement approach. (D) Correlation of flow cytometry and plate fluorimetry results for miRFP670 shows effectively identical results based on fluorescence measurement approach.

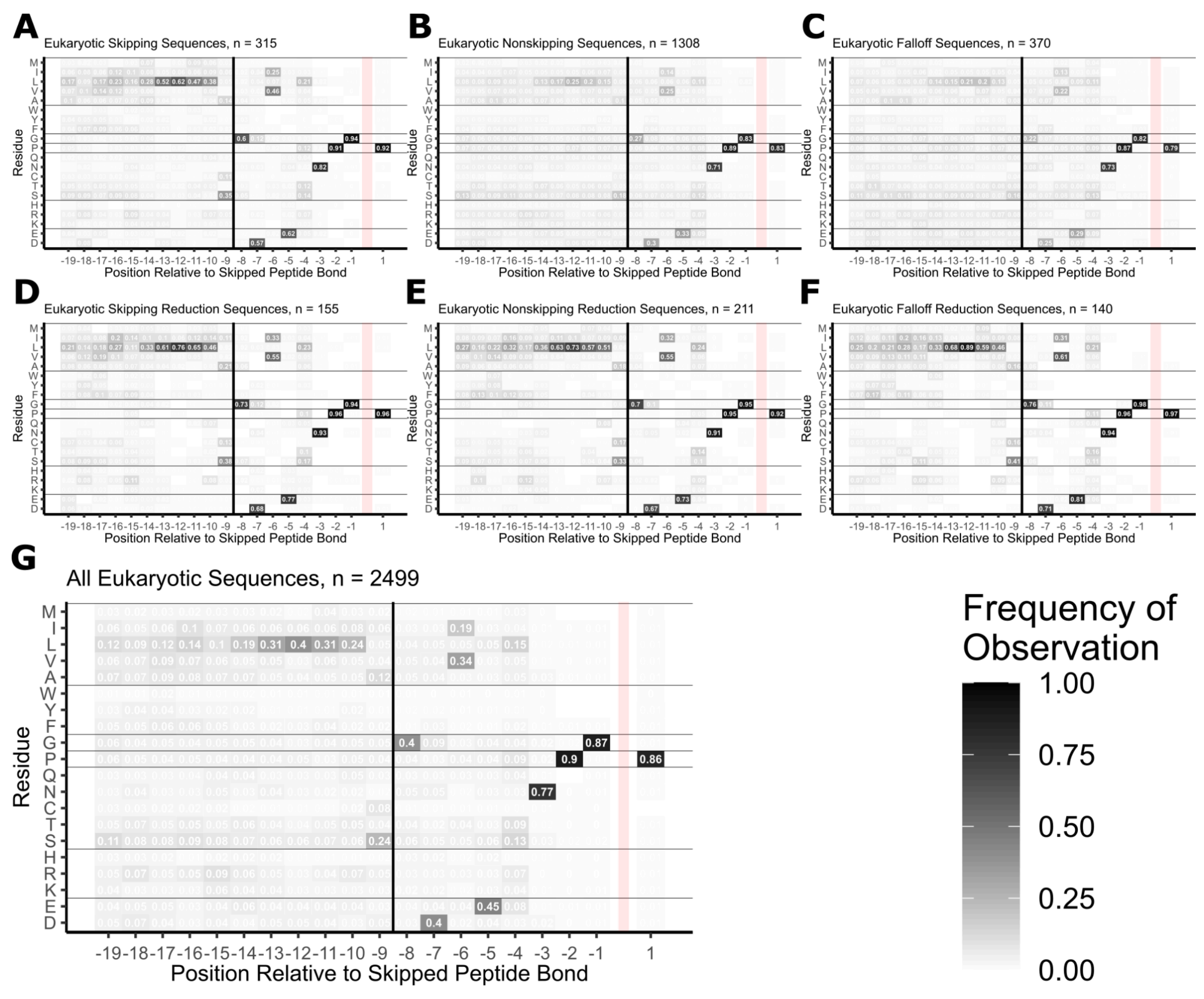


Supplemental Figure 39.

Eukaryotic residue-position usage traces. (A) Skipping peptides only. (B) Nonskipping peptides only. (C) Falloff peptides only. (D) Skipping, reduction peptides only. (E) Nonskipping, reduction peptides only. (F) Falloff, reduction peptides only. (G) all peptides pooled.

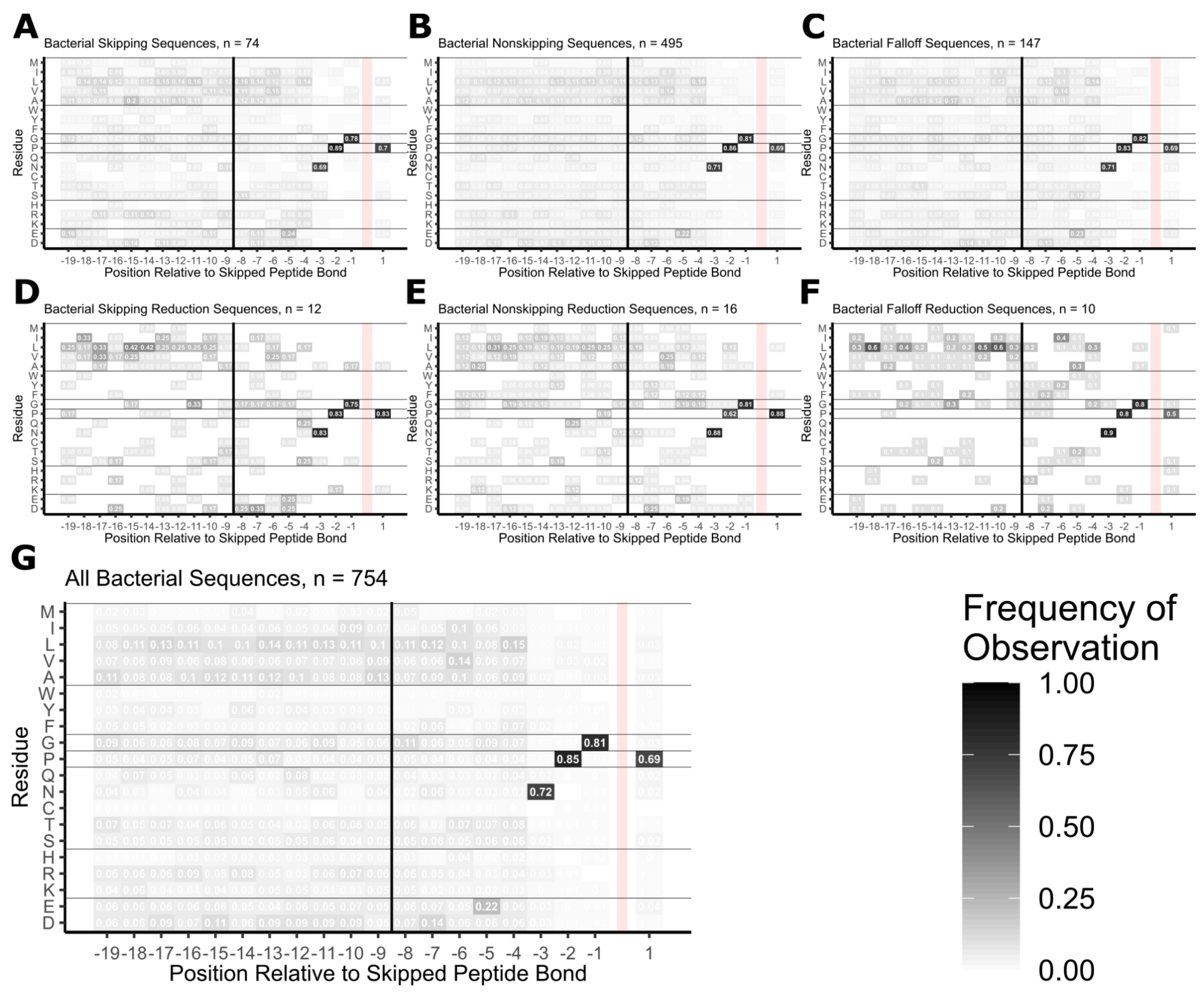


Supplemental Figure 40.

Bacterial residue-position usage traces. (A) Skipping peptides only. (B) Nonskipping peptides only. (C) Falloff peptides only. (D) Skipping, reduction peptides only. (E) Nonskipping, reduction peptides only. (F) Falloff, reduction peptides only. (G) all peptides pooled.

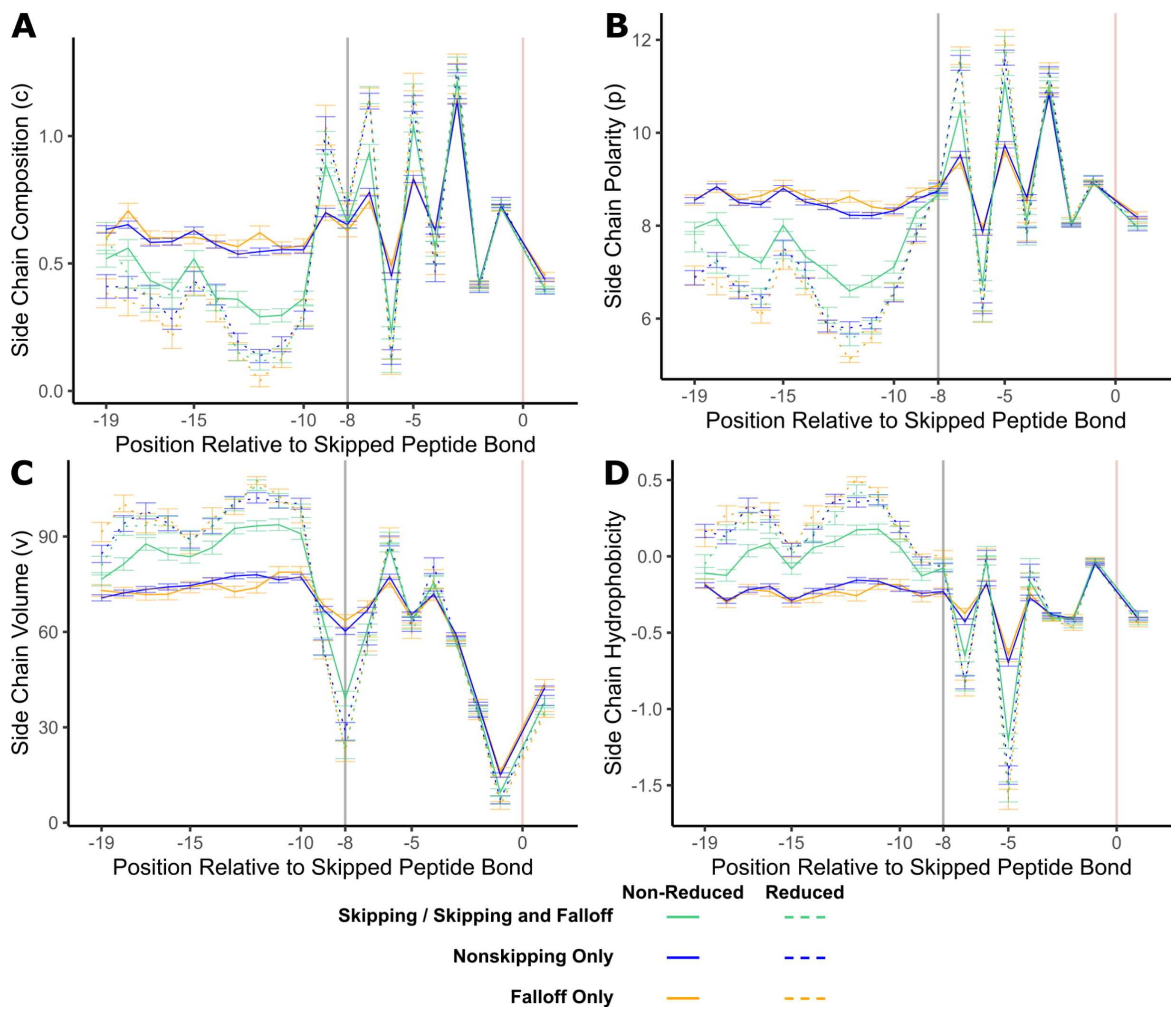


Supplemental Figure 41.

Biochemical properties for each skipping class. All samples were assigned residue biophysical properties according to the (A) composition *c*, (B) polarity *p*, or (C) volume *v* as noted in Grantham, 1974, as well as (D) the $\Delta G_x^{residue}$ for interacting with lipid membranes from Wimley and White, 1996. The mean value for each position was calculated and traced for each class. Error bars represent the standard error of the mean.

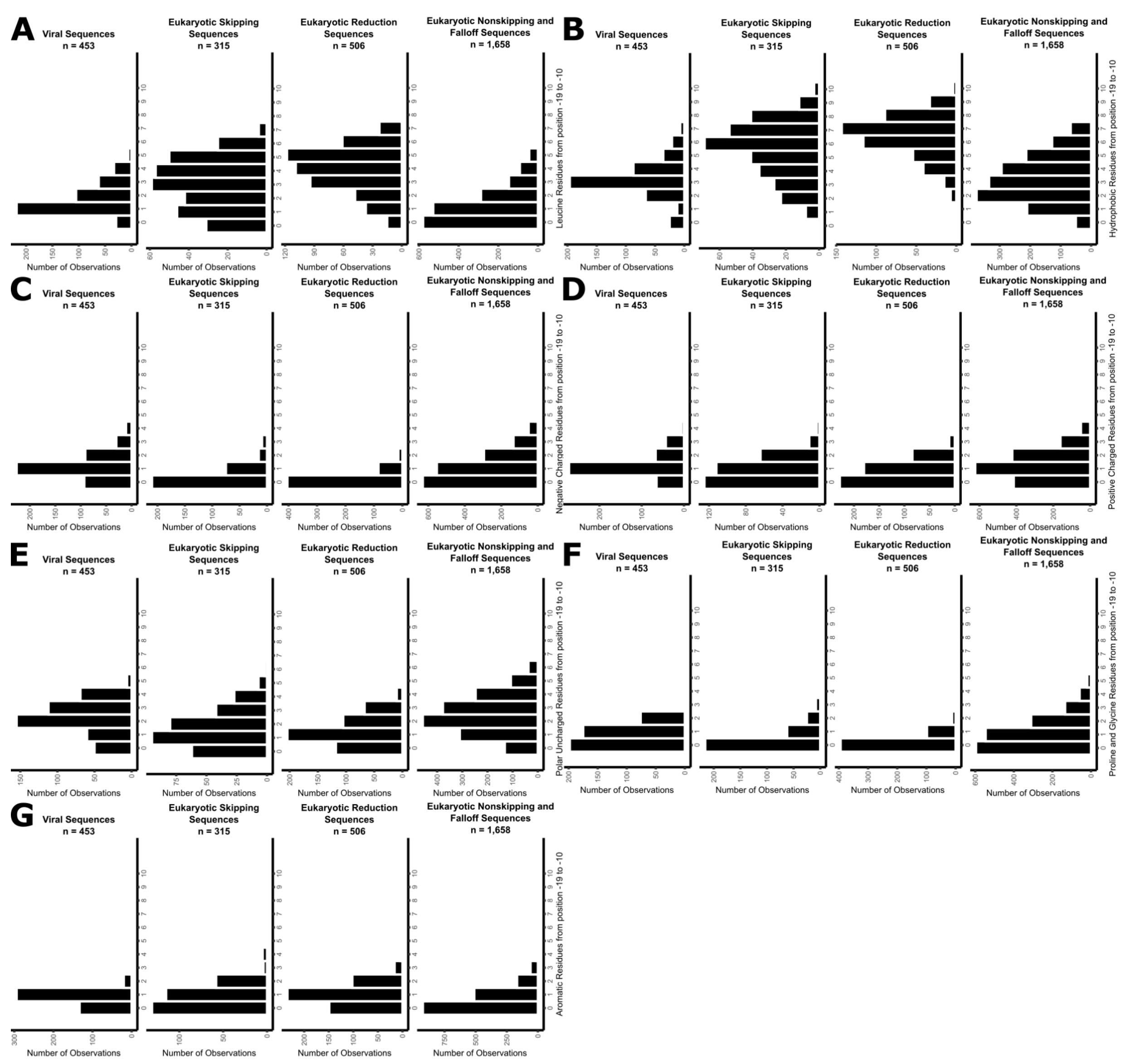


Supplemental Figure 42.

Residue use counts by peptide class and residue type from position -19 to -10 relative to the skipped peptide bond. For each panel of 4, the left graph is viral peptides (not tested), center left is eukaryotic skipping classification, center right is eukaryotic reduction classes pooled, and left is eukaryotic nonskipping and falloff classes pooled. (A) Leucine residues only. (B) Hydrophobic residues alanine, valine, isoleucine, leucine, and methionine. (C) Negative charged residues aspartate and glutamate. (D) Positive charged residues lysine, arginine, and histidine. (E) Polar uncharged residues serine, threonine, cysteine, asparagine, and glutamine. (F) Proline and glycine specifically. (G) Aromatic residues phenylalanine, tyrosine, and tryptophan.

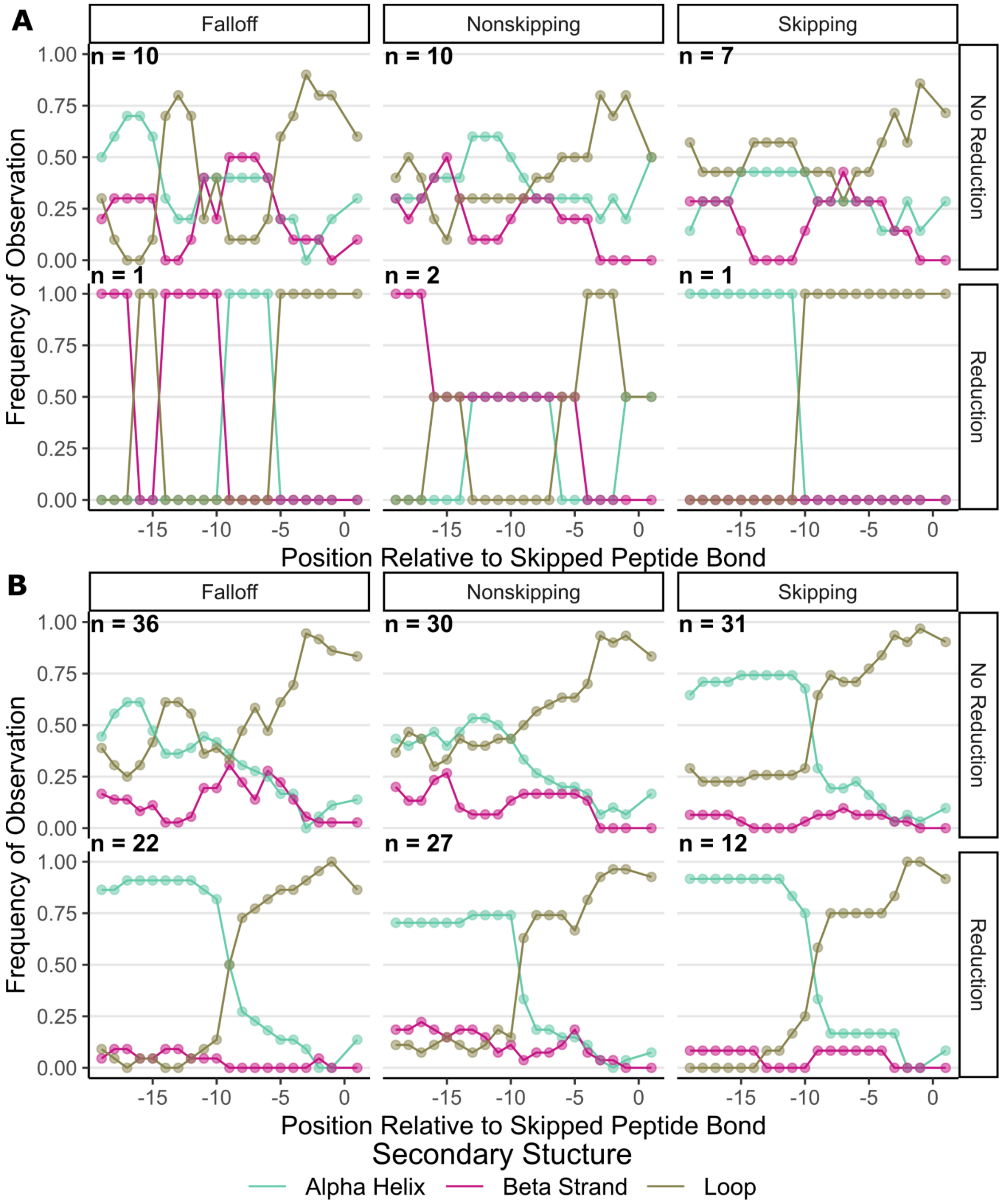


Supplemental Figure 43.

Additional secondary structure traces generated from structural prediction models. Structures for 344 proteins were predicted using AlphaFold 3, and DSSP was used to determine secondary structure of each position. Distinct 2A peptide and overall structure call (i.e. LINE1 non-LTR ORF1p-like) pairings were used to remove pseudo-duplicate data. (A) Bacterial and Archaeal peptides that were predicted only. (B) All peptides pooled, including peptides from panel A and from Figure 8H.

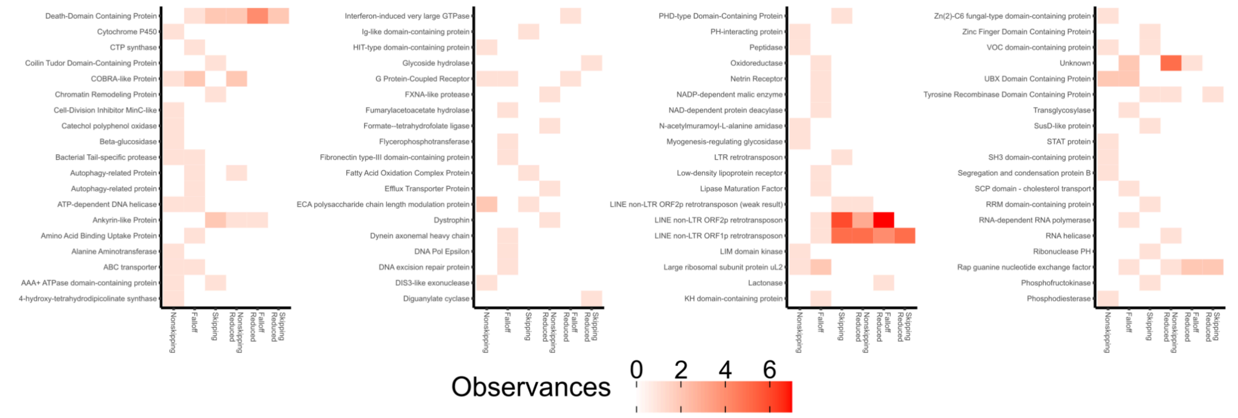

Supplemental Figure 44.

Expanded results from Figure 9A. Structure call frequency by skipping class reclassification. Most structures were only observed 1 or 2 times.
